# Development of Bioisosteric *Iboga*-alkaloids as Antinociceptive and Anxiolytic Agents with Neuroprotective Effects

**DOI:** 10.1101/2025.02.24.639821

**Authors:** Abhishek Gupta, Tuhin Bhattacharya, Subhamoy Pratihar, Sabnur Parvage, Swrajit Nath Sharma, Arnab Sarkar, Akash De, Sanmoy Karmakar, Sanjit Dey, Surajit Sinha

**Author notes:** A.G. & T.B. contributed equally.

## Abstract

The clinical importance of iboga alkaloids lies in their efficacy in reversing drug addiction and modulating drug tolerance. However, due to safety concerns, their use is restricted to appropriate medical supervision. These alkaloids often cause severe hallucinogenic effects due to differential binding to various brain receptors and cardiotoxicity by blocking the human ether-a-go-go-related gene (hERG) potassium channel. To create safer analogs, our group previously synthesized various benzofuran-containing iboga analogs with good opioid binding selectivity and excellent antinociceptive property. However, the present manuscript disclosed a step-economical and cost-effective synthesis of modified ibogaine/ibogamine analogs (**C1, C2, C3 & C4**) with bio-isosteric replacement of the indole scaffold with a benzofuran moiety, and comparing their antinociceptive/anxiolytic activity with their natural counterparts. Among the synthesized iboga analogs, the *Endo*-iboga analogs (**C2 & C4**, epimers of **C1** and **C3**, respectively) not only exhibited superior anti-inflammatory and oxidative stress-relieving activity, but also effectively improved restricted locomotor activity in a formalin-induced acute pain model in mice. These *Endo*-iboga analogs significantly elevated the levels of inhibitory neurotransmitters (GABA and dopamine) and brain-derived neurotrophic factor (BDNF) compared to their *Exo*-counterparts or previously published benzofuran-containing iboga analogs lacking the tetrahydroazepine ring. Amongst, **C2** and **C4**, the latter exhibited superior cardiac safety profile in C2C12 cells (IC_50_ = 235 µM) and showed no adverse effects on rat hearts during *in vivo* ECG tests, indicated by no significant QTc prolongation. Overall, the development of bioisosteric iboga analogs, particularly **C4**, demonstrated significant potential for acute pain management without notable cardiotoxicity, representing a breakthrough in pain therapy innovation.

## 1. Introduction

The management of acute or chronic pain, post-traumatic stress and surgical injury requires intensive supportive care to restore the appropriate health condition of a patient [1]. Till date, opioid and nonopioid analgesics remain as the two major categories of medications for the effective first-line treatment of severe pain and psychiatric disorder [2–4]. Despite having the potential in treating pain and mental illness, opioids side-by-side exert different systematic side effects and eventually develop drug dependence, tolerance, addiction and withdrawal symptoms [5–7].

In this context, Ibogaine, a monoterpene indole alkaloid isolated from the root bark of West African shrub *Tabernanthe iboga* (apocynaceae family) was reported as the potent psychedelic compound which had the capability of reversing substance use disorder (SUD) and pain alleviation even after a single dose of administration [8–11]. Such a long-lasting effect of ibogaine was mainly attributed to its long-lived active metabolite noribogaine, generated via O-demethylation when metabolized by CYP2D6 enzyme [12,13]. The actual reason for the exertion of psychedelic behavior of ibogaine still remains elusive because it targets multiple brain receptors and neurotransmitters simultaneously [14]. The differential binding of ibogaine with a wide variety of brain receptors and transporters like κ-opioid receptor (KOR), µ-opioid receptor (MOR), nicotinic α3β4, serotonin 5-HT2A, sigma (σ1 and σ2) and N-methyl-D-aspartate (NMDA) receptors, perhaps rendered for its chronic hallucinogenic side effects [15–18]. Along with this, the irreversible blockage of human ether-a-go-go-related gene (hERG) potassium channel and prolonged QT interval and development of TdP arrhythmias remained as the other major risk factors associated with ibogaine when administered at low micromolar concentration [19–23]. The limited number of case studies (mainly anecdotal reports) and lack of extensive clinical research had raised questions about ibogaine’s efficacy and safety as an antiaddictive and analgesic agent in humans [24–27]. This major shortcoming, however, motivated scientists to develop safer iboga congeners with fewer side effects, while retaining their anti-addictive properties and reducing the simultaneous targeting of multiple brain receptors [28–31].

In this regard, our previous disclosure of benzofuran-containing iboga analogs with no tetrahydroazepine ring showed efficient KOR and MOR binding affinity in the radiolabeled ligand displacement assay [32,33]. This seminal work demonstrated that the biological activity of such benzofuran-containing iboga scaffold was still retained in the absence of the seven membered O-tetrahydroazepine ring. Further validation of this proof-of-concept was justified when different functional group-modified benzofuran analogs were investigated as anti-nociceptive as well as neuromodulatory agents in formalin-induced acute pain model in mice [34].

In connection with this, the present manuscript disclosed the step-economical synthesis of modified ibogaine and ibogamine analogs, featuring bio-isosteric replacement of indole moiety with benzofuran scaffold, and their corresponding antinociceptive as well as anxiolytic effects in formalin-induced acute pain model in mice. The purpose of such bioisosteric replacement was to address the stability issues associated with the natural compounds, such as aerial oxidation and sensitivity to heat and light [35]. Importantly, the toxicity of ibogaine in human hERG channels is mainly attributed to its interaction through the indolyl –NH and the tertiary ‘N’ present in the isoquinuclidine ring. The former exhibited H-bonding interaction with Ser624 residue in the pore loop of the receptor, while the latter having pKa = 8.1, preferably remained in the protonated state under physiological pH, causing the blockage of the channels from the intracellular side and thereby inhibiting repolarization [20]. As a result, we hypothesized to replace the indole functionality of natural ibogaine/ibogamine with its bio-isosteric benzofuran scaffold while keeping the rest of the skeleton intact (**Figure 1**).

**Figure 1:**
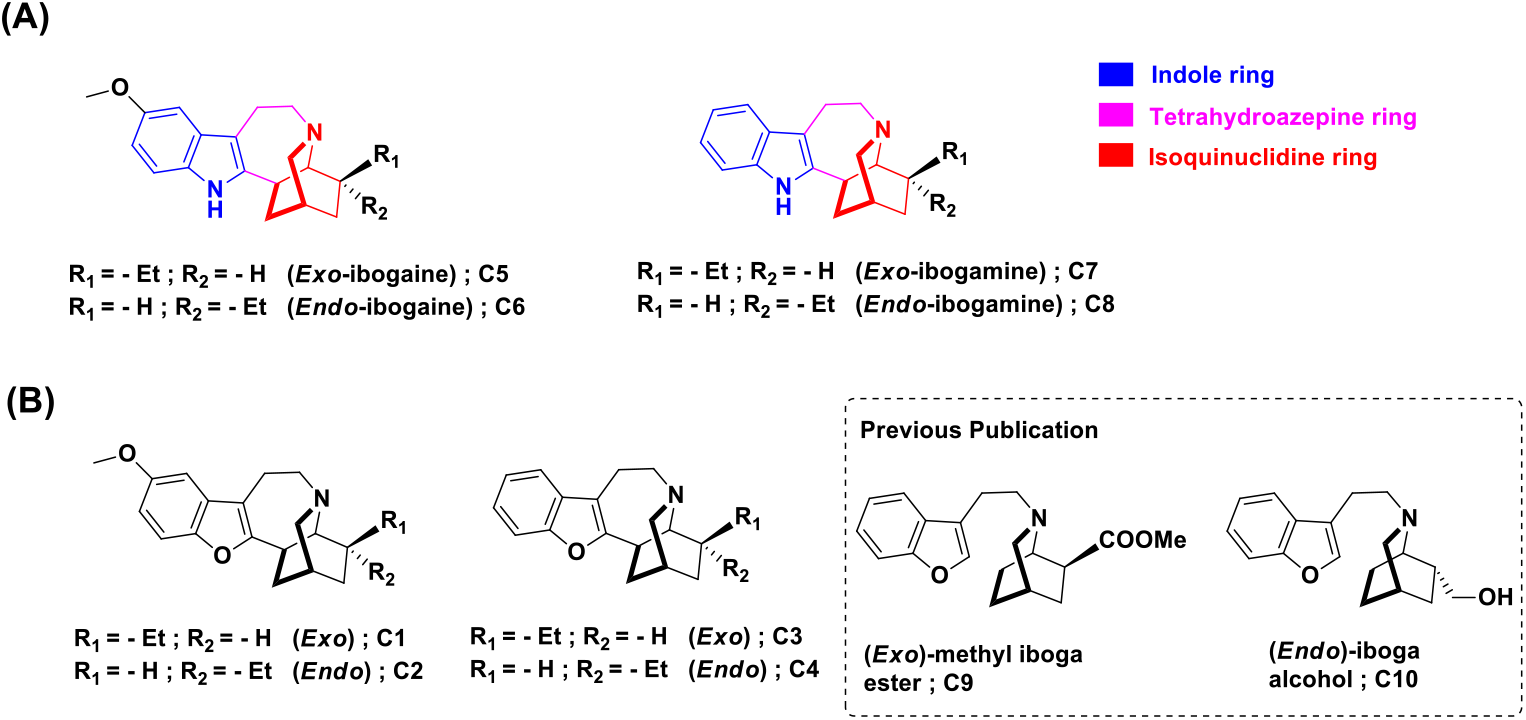
Chemical structure of **(A)**. natural ibogaine (**C5, C6**) and ibogamine (**C7, C8**) & **(B)**. bioisosteric benzofuran-containing iboga-analogs (**C1**-**C4**). **C9 & C10**: Previously published iboga analogs with abolished tetrahydroazepine ring.

Consistent with our idea, we practically synthesized four benzofuran-containing iboga analogs: **C1, C2, C3** and **C4**. All modifications were designed by ensuring that the structural connection between the three fused rings (benzofuran, tetrahydroazepine, and isoquinuclidine) would remain intact. Having synthesized these four analogs, we speculated that our proposed modification would not attenuate the overall amphipathic character of the above-mentioned molecules, a parameter crucial for penetrating blood-brain-barrier (BBB) *in vivo* [36]. In our present set of experiments, the incorporation of the *Endo* and *Exo*-isomers of natural ibogaine and ibogamine along with two previously published iboga scaffolds (i.e. *Exo*-methyl iboga ester and *Endo*-iboga alcohol) served as the appropriate choice of controls. Altogether, such cumulative incorporation of the compounds in a single set, would help us to find the most potent iboga analog in a single shot. The denotation of the above included controls was as follows: **C5 :** (*Exo*)-ibogaine, **C6 :** (*Endo*)-ibogaine, **C7 :** (*Exo*)-ibogamine, **C8 :** (*Endo*)-ibogamine, **C9 :** (*Exo*)-iboga methyl ester and **C10 :** (*Endo*)-iboga alcohol.

**Scheme 1:**
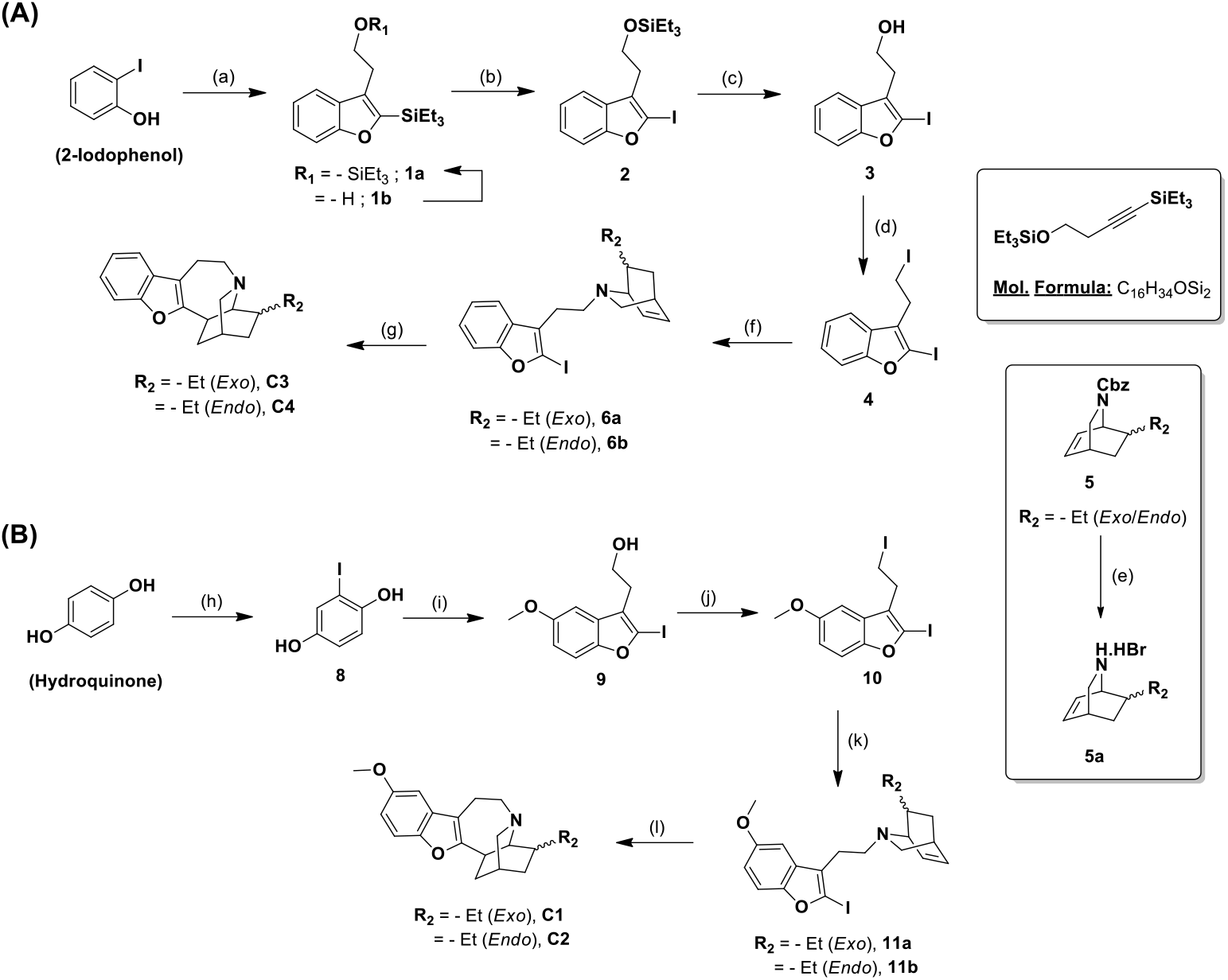
Synthetic strategy for the forward synthesis of bioisosteric benzofuran-containing **(A)**. ibogamine & **(B)**. ibogaine analogs. *Reagents & conditions:* (a). (i) C_16_H_34_OSi_2_, Na_2_CO_3_, LiCl, Pd(OAc)_2_, DMF, (ii) TESCl, Imidazole, DCM, 93% (over 2 steps); (b). NIS, CH_3_CN, 75%; (c). TBAF, THF, 69%; (d). I_2_, PPh_3_, Imidazole, DCM, 91%; (e). HBr (∼33% in AcOH), DCM, (quantitative); (f). **5a**, Cs_2_CO_3_, CH_3_CN, 69% (including **6a & 6b**); (g). PPh_3_, Pd(OAc)_2_, HCOONa, DMF, 72% (including **C3 & C4**); (h). (i) TBDMSCl, Imidazole, DMF, (ii) NIS, TFA, CH_3_CN, (iii) TBAF, THF, 57% (over 3 steps); (i). (i) C_16_H_34_OSi_2_, Na_2_CO_3_, LiCl, Pd(OAc)_2_, DMF, (ii) TESCl, Imidazole, DCM, (iii) NIS, CH_3_CN, (iv) TBAF, THF, (v) K_2_CO_3_, MeI, DMF, 38% (over 5 steps); (j). I_2_, PPh_3_, Imidazole, DCM, 85%; (k). **5a**, Cs_2_CO_3_, CH_3_CN, 72% (including **11a & 11b**); (l). PPh_3_, Pd(OAc)_2_, HCOONa, DMF, 81% (including **C1 & C2**).

All the synthesized bioisosteres of iboga analogs, along with their natural counterparts, were initially screened for pain-alleviating and anxiolytic activities using the formalin-induced acute pain model in mice. The results demonstrated that, among the sets, the *Endo*-analogs of the benzofuran-containing ibogaine or ibogamine (**C2** and **C4**, respectively) along with the natural one (**C6** and **C8**) exclusively retained the antihyperalgesic or antinociceptive activity when compared with their corresponding *Exo*-counterparts just after the formalin administration. The effects of the analogs were evaluated in terms of the measurement of (1) paw licking/withdrawal activity, (2) tail flick latency and (3) mechanical allodynic activity of the formalin-treated mice throughout the different phases or time points. The *Endo*-compounds (**C2, C4, C6** and **C8**) seemed to be effective among the sets, as they were not only able to increase the overall locomotor activity, but also had the capability of elevating the exploratory activity of the formalin-induced pain model in mice with excellent anxiolytic properties. Additionally, **C2, C4, C6** and **C8** were able to scavenge local ROS generation in paw regions post-treatment with formalin and elevate the anti-inflammatory response by preventing the increase of substance P, CGRP, and neurokinin 1 receptor (NK1R) at the spinal level. However no such driscriminatory effect between these *Endo*-analogs (**C2, C4, C6** and **C8**) and their respective *Exo*-counterparts (**C1, C3, C5** and **C7**) was noted when their effects on relative protein expression of p65 were investigated in paw lysates. While examining the neuromodulatory effects of these analogs in L4-L6 spinal region, elevated levels of dopamine/GABA along with reduced level of spinal serotonin (5-HT) and glutamate, were observed in all cases of **C2, C4, C6** and **C8** treatments. However, in terms of restoring the relative protein expression of brain derived neurotrophic factor (BDNF), **C2** along with **C1** and **C3** turned out to be most effective among the series and their extent of upregulation of BDNF expression were almost comparable to that of the untreated control. In fact, such restorative effect completely outperformed the individual post-treatments with **C9 & C10**. Amongst the potent *Endo-*iboga compounds (**C2 & C4**), **C4** showed more cytocompatibility in C2C12 cells and exhibited no deleterious effect on the corrected QT interval (i.e. QTc) in rats when analyzed by ECG tests. Overall, this research, utilizing a formalin-induced acute pain model in mice, identified **C4** as the most promising *Epi-*iboga scaffold, which held excellent antinociceptive and anti-inflammatory property with minimal cardiotoxicity. The examination of its neuromodulatory mechanism also indicated that it holds superior potential for future therapeutic applications.

## 2. Results and Discussion

### 2.1. Chemical synthesis of bioisosteric iboga analogs

The chemical synthesis of the bioisosteric benzofuran-containing ibogaine (**C1 & C2**) and ibogamine (**C3 & C4**) analogs was initiated using commercially available hydroquinone and 2-iodophenol, respectively, adopting the previously published methodology, which mainly involved the following synthetic steps: (1) Larock heteroannulation, (2) Vinylic ipso-iodination, (3) Removal of silyl protection group, (4) Homoallylic iodination, (5) Base-mediated coupling with isoquinuclidine ring and lastly, (6) Reductive Heck coupling [37]. The Di-iodo benzofuran compounds (**4** and **10**) were synthesized from their respective starting materials in five and nine synthetic steps with ∼44% and ∼20% yield, respectively, and then, were individually subjected to base-mediated alkylation with compound **5a**. The successful coupling of the isoquinuclidine ring with the benzofuran motif mainly relied on exploiting the difference of reactivity between the aliphatic (sp3) and vinylic (sp2) iodide. The *N*-alkylated intermediates, thus generated for ibogaine (**11a & 11b**) and ibogamine (**6a & 6b**) were next subjected to reductive Heck coupling. This process successfully accomplished the installation of the tetrahydroazepine ring, connecting the benzofuran motif with the isoquinuclidine skeleton. The total synthesis of the benzofuran-containing ibogaine and ibogamine was achieved in 12 and 8 synthetic steps with an overall yield of ∼11% and ∼20%, respectively (including both the *Exo* and *Endo*-isomers) (**Scheme 1**). Cumulatively, the relative ease in chemical synthesis and adoption of comparatively lesser number of synthetic steps via cost-effective approach, made this methodology amenable to bulk scale production of bioisosteric iboga analogs. The other iboga compounds (**C5**-**C10**), highlighted in **Figure 1**, were also synthesized following our previously published reports [34, 37]. The inclusion of these compounds in our experiments as appropriate controls was essential for assessing the biological efficacy of **C1**-**C4**. In this context, it is worth mentioning that Sames *et al*. recently reported a benzofuran-containing iboga analog, termed oxanoriboga alkaloid, which had a lower probability of causing cardiac arrhythmias and exhibited higher reversal activity against opioid dependence in animal models than natural ibogaine/noribogaine [38].

### 2.2. Acute pain model and experimental design

The formalin-induced acute pain model is an established method for assessing a compound’s analgesic effect in laboratory mice [39]. After injecting formalin into the paw, two separate phases of reaction are usually seen: (1) acute phase 1 and (2) tonic phase 2. Phase 1 pain behaviors develop due to the direct chemical stimulation of peripheral nociceptors. In contrast, phase 2 pain behaviors result from a sustained peripheral expression of phase 1 induced spinal cord hyperexcitability and ongoing stimulation of nociceptors by inflammatory and ROS mediators. In evaluating the therapeutic efficacy of the synthesized iboga analogs in a formalin-induced acute pain model, the timing of intraperitoneal (IP) treatment immediately after pain induction, was crucial, resembling a curative clinical setup. The allodynic behavior was recorded at multiple time points up to 24 h post-formalin injection, whereas the locomotor activity, anxiety-like response and inflammatory mediators were measured 24 h post-treatment.

### 2.3. Antihyperalgesic, antinociceptive and antiallodynic activity of iboga analogs after formalin administration

Although natural ibogaine was well-documented in the literature for exhibiting antinociceptive and antiaddictive properties in morphine-tolerant mice model, it was ineffective in significantly reducing nociception in the acute pain model by itself [40, 41]. In this regard, the objective of the present work was to ascertain the independent antinociceptive qualities of the newly synthesized bioisosteric iboga analogs Therefore, formalin-induced paw licking, tail flick latency, and paw withdrawal behaviors were evaluated to determine the pain-relieving activity of the compounds (**Figure 2**). Manual Von Frey monofilaments were applied vertically to the unrestrained animals with increasing force to measure mechanical allodynia or tactile hypersensitivity at multiple time points following formalin injection, as previously described (**Figure 3**).

**Figure 2:**
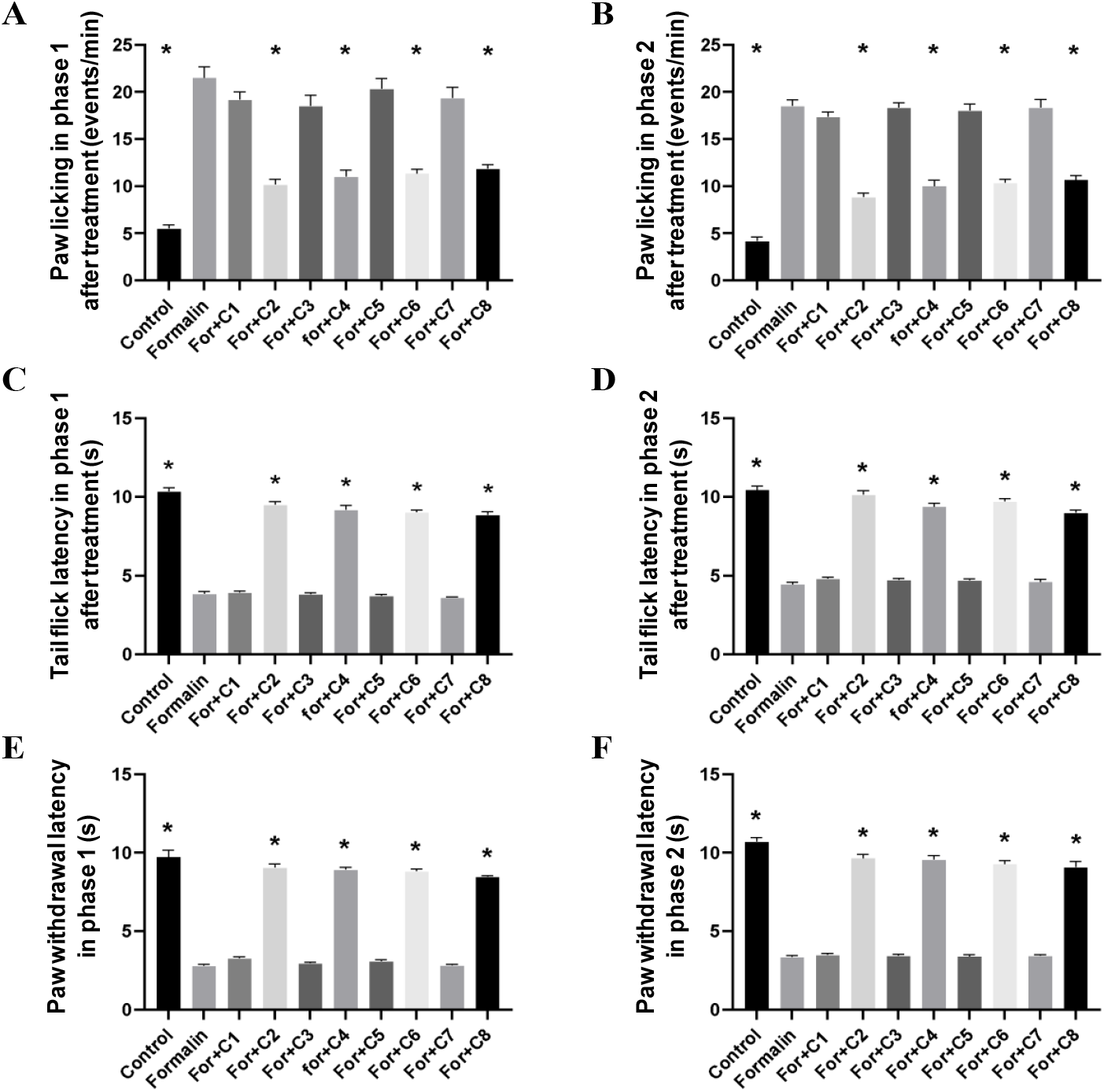
Measurement of paw licking events, tail flick and paw withdrawal latency. **(A)** Paw licking events in phase 1 after formalin and iboga analog treatment. **(B)** Paw licking events in phase 2 after formalin and iboga analog treatment. **(C)** Tail flick latency (measured in seconds) in phase 1 after formalin and iboga analog treatment. **(D)** Tail flick latency (measured in seconds) in phase 2 after formalin and iboga analog treatment. **(E)**. Paw withdrawal latency (measured in seconds) in phase 1 after formalin and iboga analog treatment. **(F)**. Paw withdrawal latency (measured in seconds) in phase 2 after formalin and iboga analog treatment. Iboga analogs were administered at a dose of 30 mg/kg via IP injection (n = 6). Data represented as Mean ± SEM, \**P* < 0.05, * indicated significant difference with formalin-treated group.

**Figure 3:**
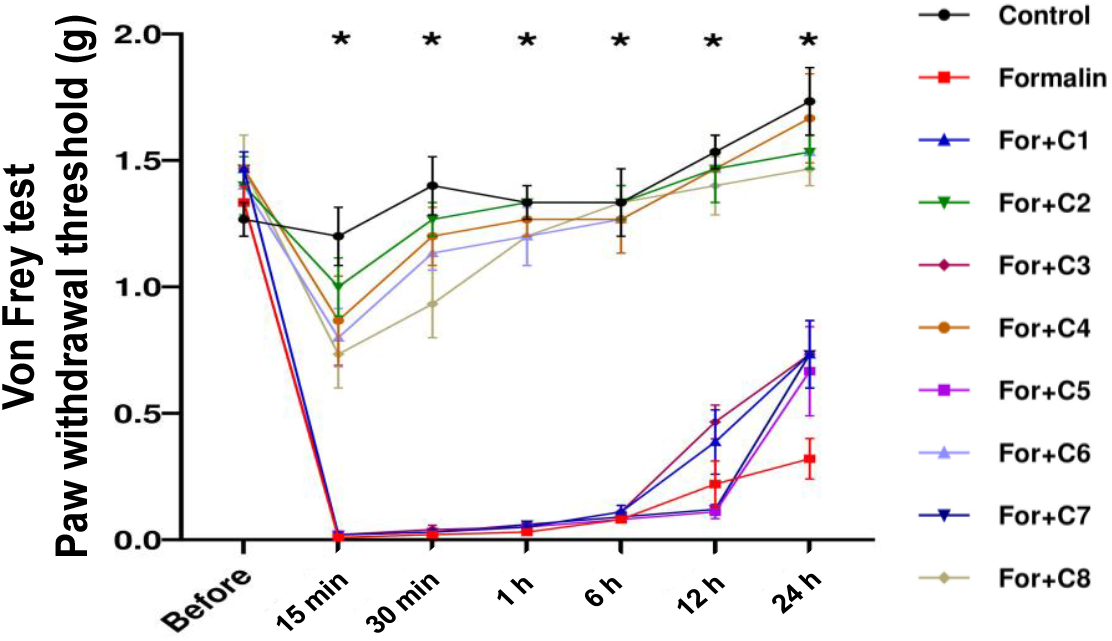
Characterization of mechanical allodynia after formalin induced acute pain. Paw withdrawal threshold (expressed in g) was recorded for a period of 15 min, 30 min, 1 h, 6 h, 12 h, 24 h post-formalin and iboga analog treatment. Trendlines were drawn for each treatment group at multiple time points. Two-way ANOVA followed by Tukey’s multiple comparison tests were performed to find any significant difference. Iboga analogs were administered at a dose of 30 mg/kg via IP injection (n = 6). Data represented as Mean ± SEM, \**P* < 0.05, * indicated significance difference between control, **C2, C4, C6** and **C8** with formalin-treatment at the given time points.

The biphasic response, triggered by the introduction of formalin, resulted in increased paw licking events (PLE) during both phases of the treated mice. *Endo-*ibogaine and ibogamine derivatives from both the structural configuration (benzofuran/natural), i.e., **C2, C4, C6** and **C8**, respectively, showed significant inhibition of paw licking events in comparison to formalin-treated group (*P* < 0.05) **(Figure 2A & 2B)**. In between the two structural derivatives, the benzofuran analogs (**C2** and **C4**) showed a better antinociceptive effect than that of natural **C6** and **C8**. On the other hand, the *Exo*-analogs (**C1, C3, C5** and **C7**) failed to show any significant antinociceptive effects in formalin-induced nociception (*P* > 0.05). To evaluate the effects of iboga analogs on heat-induced hyperalgesia, tail flick test and paw withdrawal test were conducted, where the tail flick latency (TFL) and paw withdrawal latency (PWL) were measured in seconds. The paw withdrawal and tail flick behaviors, which indicate sensitivity to unpleasant stimuli, demonstrated acute pain response in both stages when triggered by a spinal reflex. *Endo-*iboga derivatives, namely, **C2, C4, C6** and **C8** showed significant hyperalgesic behavior (*P* < 0.05) in both phase 1 and phase 2, where both TFL (**Figure 2C & 2D**) and PWL (**Figure 2E & 2F**) were restored at an extent comparable to that of the untreated control mice. On contrary, *Exo*-iboga analogs, i.e., like **C1, C3, C5** and **C7** failed to exert any significant differential effects in this regard. The paw withdrawal threshold (expressed in g) was significantly lowered (*P* < 0.05) in the formalin-treated group at all time points (15 min, 30 min, 1 h, 6 h, 12 h, and 24 h) (**Figure 3**). However, *Endo*-iboga analogs (**C2, C4, C6** and **C8**) treatment did not result in a significant decrease in the paw withdrawal threshold due to formalin injection at any observed time points, indicating the absence of allodynia. Consistent with previous findings, *Exo*-iboga analogs (**C1, C3, C5** and **C7**) also failed to show any significant anti-allodynic effects across all time points.

### 2.4. Amelioration of compromised locomotor activity and anxiety-like behavior of mice by iboga treatment

When animals experience pain, it is said that they repress their ambulatory behavior, which is associated with mobility [42]. To investigate whether the increased latencies in the tail flick and hot plate tests were due to actual antinociception or a general lack of responsiveness, a locomotor activity test was conducted in an open field. Consequently, the antidepressant efficacy of the analogs was assessed using an open field test, where the animals’ locomotor activity was observed in a circular open field, both before and after receiving treatments with formalin and iboga analogs (**Figure 4**). After 24 h of formalin treatment, there was a decrease observed in the measurement of locomotor activity including mean speed (MS, measured in m/s) (**Figure 4E**) and distance travelled (DT, measured in m) (**Figure 4C**), which was not evident in the *Endo*-iboga epimers (**C2, C4, C6**, and **C8**)-treated groups. Here, again the *Exo*-iboga analogs (**C1, C3, C5** and **C7**) failed to ameliorate the pain-induced restricted locomotor activity 24 h after formalin treatment. This study also proved that the iboga analogs had no deleterious effects on locomotion, confirming that they were devoid of any adverse effects on muscle and central nervous system (CNS). Overall, the formalin treatment in mice grossly reduced the spontaneous movement and distance travelled, whereas the *Endo*-iboga epimers effectively restored the movement to an extent like untreated control.

**Figure 4:**
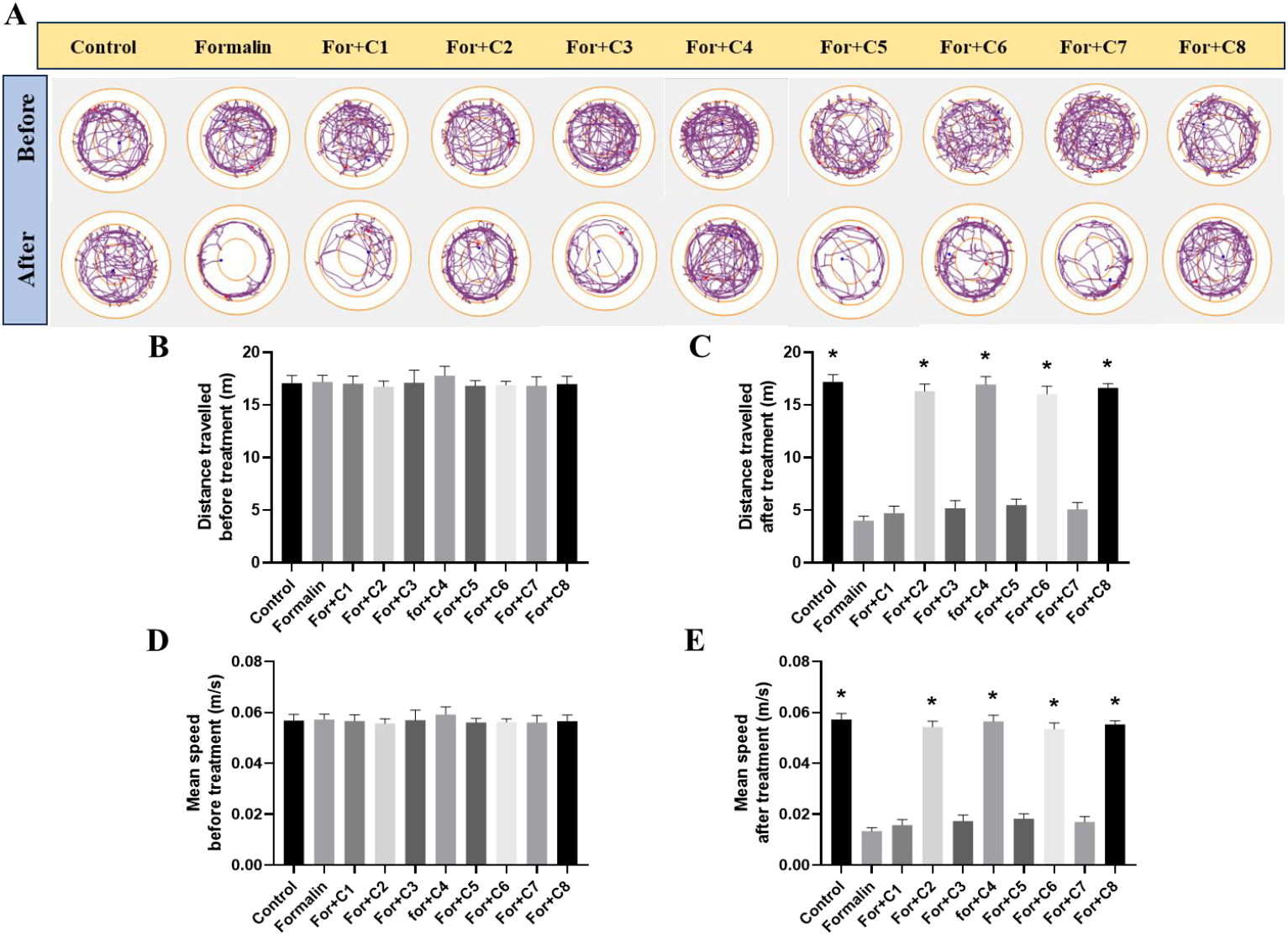
Evaluation of locomotor activity study in an open field. **(A)**. Track plot showing locomotor activity before and after iboga analog treatment in open field. **(B)**. Distance traveled (DT, measured in meter) before formalin or iboga treatment. **(C)**. Distance traveled after formalin and iboga analog treatment. (D). Mean speed (MS, measured in meter/second) before formalin or iboga treatment. **(E)**. Mean speed after formalin and iboga analog treatment. Iboga analogs were injected at 30 mg/kg dose via IP injection (n = 6). Values were Mean ± SEM, \**P* < 0.05, * indicated significant difference with formalin-treated group.

Anxiety-like behavior in mice increased when they were subjected to external chemical stimuli (i.e. formalin) to induce acute pain. Therefore, anxiety-like behavior in an elevated plus maze was used to assess the anxiolytic effect of the compounds [43].

Before and after receiving treatments with formalin and iboga analogs, the movements of the mice in the maze’s open and closed arms were observed (**Figure 5**). Important measurements of anxiolytic behavior included the number of entries (NE) and time spent in seconds (TS) in the open arm. After receiving formalin, the animal’s movement in the open arm the plus maze was restricted significantly (*P* < 0.05), as evidenced by a heat map of the animal’s movement (**Figure 5A**), while the *Endo*-iboga analog (**C2, C4, C6** and **C8**) therapy clearly preserved the animal’s ability to travel around the open arm of the plus labyrinth (**Figure 5C & 5E**). Here also, the IP treatment with *Exo*-iboga analogs failed to significantly counteract the potent anxiogenic action of formalin injection (*P* > 0.05).

**Figure 5:**
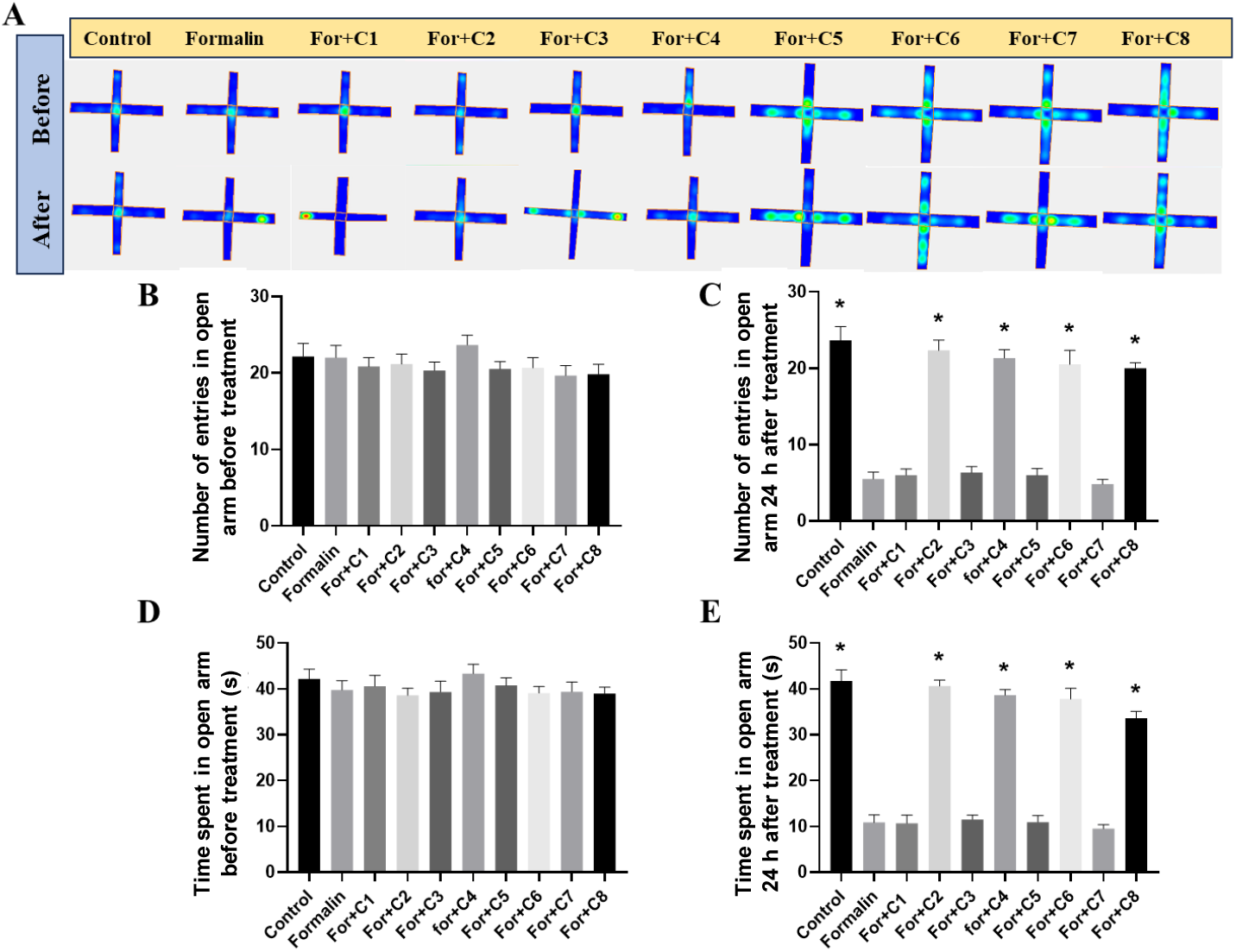
Evaluation of anxiety-like behavior in elevated plus maze. **(A)**. Heat map of anxiety-like activity measured before and after iboga analog treatment in Elevated Plus Maze (EPM). **(B)**. Number of entries in the open arm before formalin or iboga treatment. **(C)**. Number of entries in the open arm after formalin and iboga treatment. **(D)**. Time spent in the open arm (measured in seconds) before formalin/iboga treatment. **(E)**. Time spent in the open arm (measured in seconds) after formalin and iboga treatment. Iboga analogs were injected at 30 mg/kg dose intraperitoneally (n = 6). Data were Mean ± SEM, \**P* < 0.05, * indicated significant difference with formalin treated group).

### 2.5. Reduction of oxidative stress by iboga analogs after formalin administration

To assess whether the iboga analogs could reduce oxidative stress induced by post-formalin treatment, their antioxidant properties in the paw tissue lysate were evaluated replicating the earlier report (**Figure 6**) [44]. Reduced glutathione level (expressed in μmoles/mg of protein), an important marker for inherent antioxidant capacity of cells, was significantly (*P* < 0.05) elevated after treatment with all iboga analogs (both *Endo* and *Exo*) over the formalin-treated group (**Figure 6A**). The metalloprotein Superoxide dismutase 1 (SOD1), which shields cells from oxidative damage, also showed increased activity (measured in U/mg of protein) (*P* < 0.05) after administration of iboga analogs in contrast to formalin treatment (**Figure 6B**). Catalase (CAT) activity, measured in terms of μmoles of H_2_O_2_ reduced/mg of protein, is essential for neutralizing reactive oxygen species (ROS) through the breakdown of H_2_O_2_. Treatment with all the iboga analogs significantly increased the catalase activity (*P* < 0.05), which was much higher than animals-treated with formalin alone (**Figure 6C**). In contrast to the formalin-treated groups, the *Endo*-iboga analog-treated groups did not show any significant increase in TBARS (thio-barbituric acid reactive substrates), indicative of lipid peroxidation (LPO) (**Figure 6D**) and their extent of restoration was almost similar to that of untreated control.

**Figure 6:**
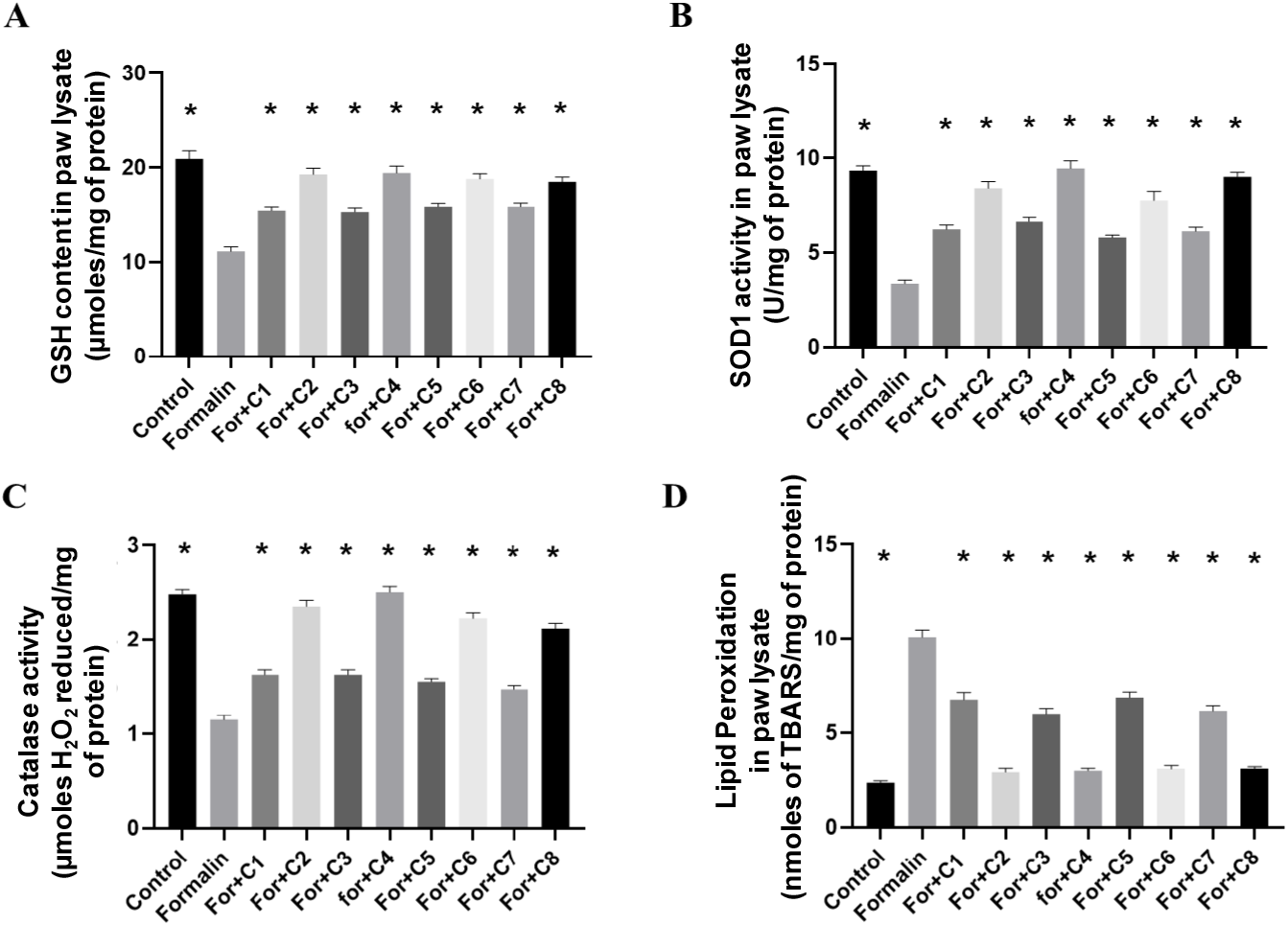
Evaluation of ROS scavenging activity in paw lysate. **(A)**. Reduced glutathione levels (GSH, measured in μmoles/mg of protein) in paw lysate after treatment. **(B)**. Superoxide dismutase 1 (SOD 1) activity (measured in U/mg of protein) in paw lysate after treatment. **(C)**. Catalase activity (measured in μmoles of H_2_O_2_ reduced/mg of protein) in paw lysate after treatment. **(D)**. Lipid Peroxidation (LPO, measured as nmoles of TBARS/mg of protein) in paw lysate after treatment. Iboga analogs were injected at 30 mg/kg dose via IP injection (n = 6). Data were Mean ± SEM, \**P* < 0.05, * indicated significant difference with formalin treated group).

### 2.6. Effects of iboga analogs on inflammatory onset in paw tissue

Usually, the anti-inflammatory effects of the compounds were typically evaluated by monitoring the time-dependent release of several pro-inflammatory chemicals, triggered by the redox imbalance caused by formalin, ultimately leading to an inflammatory cascade [45]. Substance P, an undecapeptide neurotransmitter, is known to trigger an inflammatory pain response by activating TRPA1 (Transient Receptor Potential Ankyrin 1) and causes a subsequent calcium influx [46]. At the same time, it also plays a crucial role in the transmission of pain information to the CNS.

The expression level of Substance P (pg/μg of protein) in the paw lysate was significantly elevated after formalin treatment compared to the untreated control mice. However, it returned to levels similar to the untreated control only with the injection of *Endo*-iboga compounds (**C2, C4, C6** and **C8**) (**Figure 7A**). Conversely, the *Exo*-iboga analogs failed to downregulate the level of expression of Substance P (*P* > 0.05). The neurokinin-1 receptor (NK1R) is a G protein-coupled receptor (GPCR) and plays an important role in pain transmission and several other physiological processes [47]. The level of expression of NK1R was significantly downregulated upon treatment of *Endo-*iboga analogs and became almost comparable to that of the untreated control (**Figure 7C**). Calcitonin gene-related peptide (CGRP) is a neuropeptide which also plays a crucial role in neuronal pain transmission, modulation and inflammation [48]. To evaluate the effects of iboga analogs, the expression level of CGRP in paw lysate was tested. The findings revealed that iboga analogs significantly downregulated CGRP expression compared to the formalin-treated mice (**Figure 7B**). As reported earlier, elevated levels of Substance P can induce phosphorylation of p65, thereby affecting its nuclear localization [49]. In the present study, immunoblot analysis clearly showed that formalin injection promoted the nuclear localization of p65, the master regulator of inflammatory onset in cells. However, this upregulated expression was significantly reduced following post-treatment with all iboga analogs, when administered locally in the paw tissue (*P* < 0.05) (**Figure 7D**).

**Figure 7:**
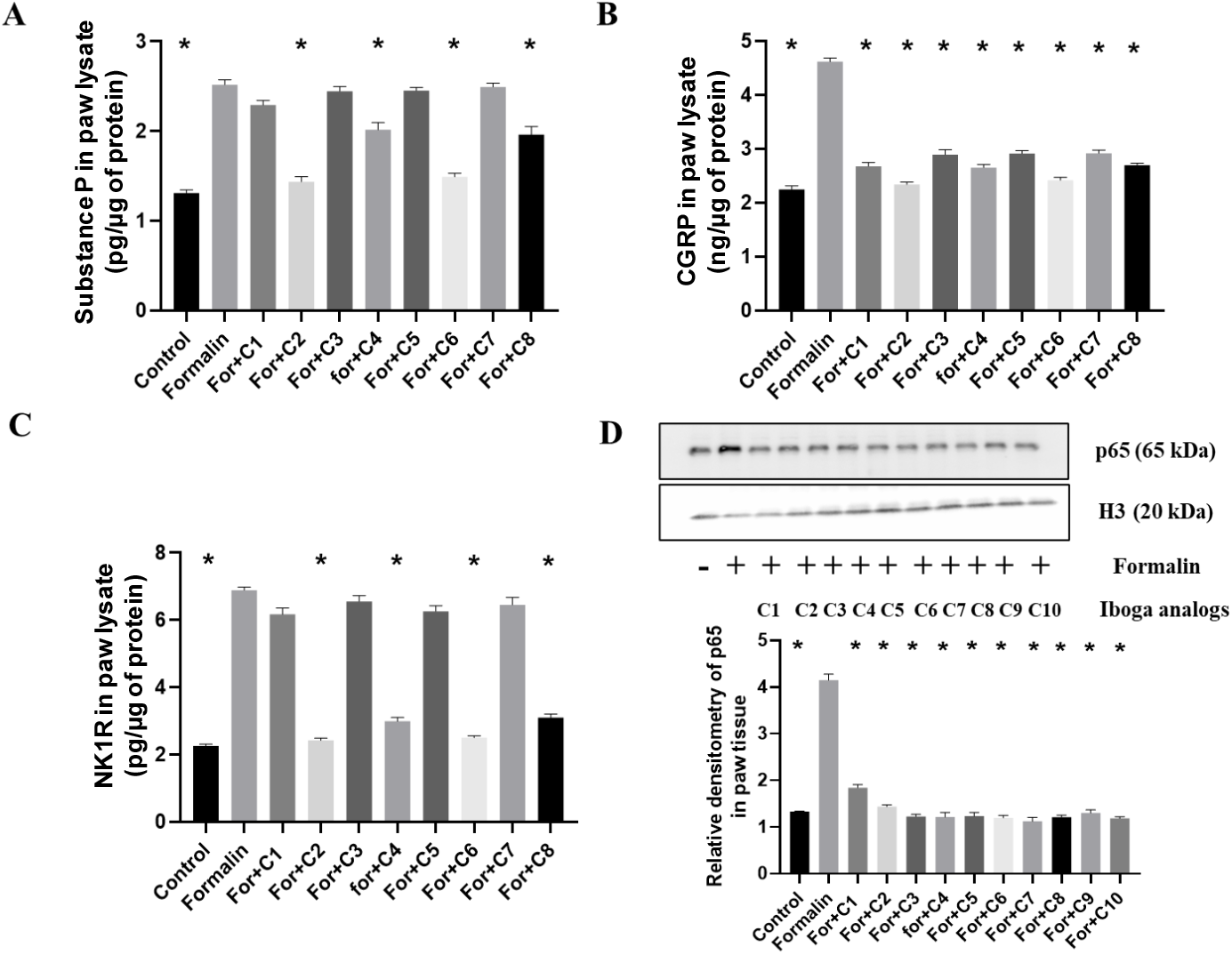
Evaluation of inflammatory markers in paw tissue. **(A)**. Substance P levels (measured as pg/μg of protein) in paw lysate after treatment. **(B)**. Calcitonin gene related peptide (CGRP, measured as ng/μg of protein) in paw lysate after treatment. **(C)**. Neurokinin 1 receptor (NK1R) in paw lysate (measured as pg/μg of protein) after treatment. **(D)**. Immunoblot of nuclear p65 from paw tissue after formalin and iboga analog treatment with densitometric analysis. Iboga analogs were injected at 30 mg/kg dose via IP injection (n = 6). Values were Mean ± SEM, \**P* < 0.05, * indicated significant difference with formalin treated group).

### 2.7. Neuromodulatory effects of iboga analogs on neurotransmitters and neurotrophic factor in spinal L4-L6 region

The spinal L4-L6 area, where the sciatic nerves from paw tissue terminate, is typically impacted by neuropathic pain or paw inflammation. Moreover, the previous reports of acute pain showed the evidence for spinal glutamatergic neuron activation cascade. To ascertain whether the excitatory neurotransmitter glutamate and the inhibitory neurotransmitters dopamine and GABA are modulated by iboga chemicals or not, we wished to evaluate the effects of the synthesized compounds (**Figure 8**). Furthermore, the roles of spinal serotonin (5-HT) and brain-derived neurotrophic factor (BDNF) in acute pain were assessed in an acute inflammatory test model in mice induced by formalin, as previously shown.

**Figure 8:**
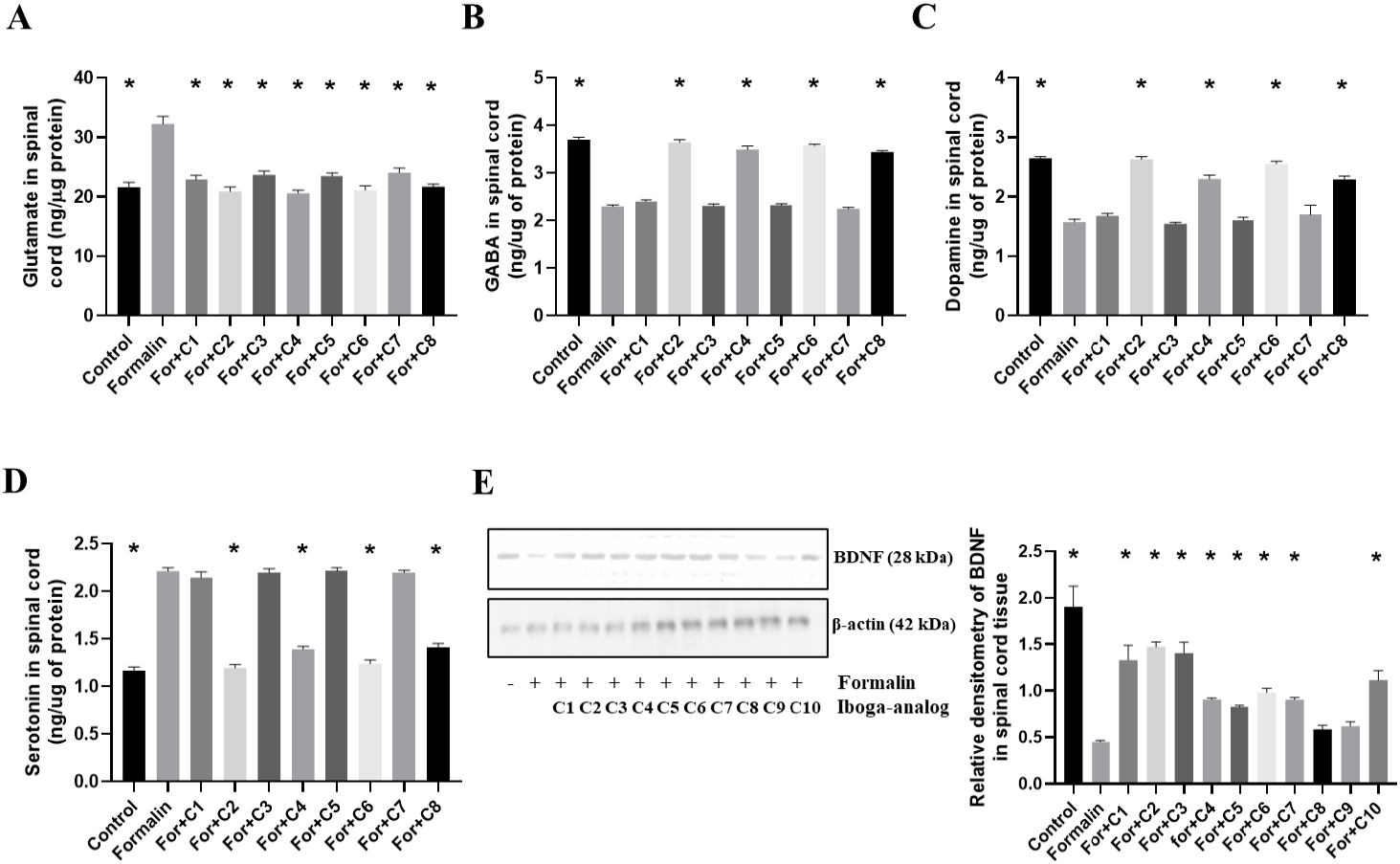
Evaluation of the levels of neurotransmitters and BDNF in spinal L4–L6 segment lysate. **(A)**. Glutamate levels (measured as ng/μg of protein) after treatment. **(B)**. γ-Aminobutyric acid (GABA, measured as ng/μg of protein) levels after treatment. **(C)**. Dopamine levels (measured as ng/μg of protein) after treatment. **(D)**. Serotonin levels (measured as ng/μg of protein) after treatment. **(E)**. Immunoblot analysis of BDNF expression and densitometric analyses after treatment. Iboga analogs were injected at 30 mg/kg dose intraperitoneally (n = 6). Data were Mean ± SEM, \**P* < 0.05, * indicated significant difference with formalin treated group.

Elevated level of spinal glutamate was observed in case of formalin treatment [50]. However, all the iboga analog-treatments significantly downregulated the increased glutamate levels (*P* < 0.05), reducing them to a level comparable to the untreated control (**Figure 8A**). On the other hand, a marked decrease in GABA and dopamine levels was observed following formalin injection [51,52]. Upon treatment of the *Endo*-iboga epimers (**C2, C4, C6** and **C8**), significant elevation was noted both in the level of GABA and dopamine (**Figure 8B & 8C**), closely resembling those of the untreated control. However, all the *Exo*-iboga analogs failed to upregulate GABA and dopamine levels in the spinal region after formalin injection. Additionally, the *Endo*-iboga analogs demonstrated a significant downregulation (*P* < 0.05) in serotonin levels after formalin injection (**Figure 8D**) [53]. As somatosensory regulation of pain signals in spinal cord involves BDNF, a decreased level of BDNF was found after formalin injection, which was again significantly upregulated with the treatments with iboga compounds [36]. This experimental result depicted the positive role of iboga analogs in neural growth and plasticity against the formalin-induced acute inflammatory pain (**Figure 8E**).

### 2.8. Toxicological evaluation of iboga analogs

For a comprehensive understanding of the safety profile of the synthesized iboga analogs, the *in vitro* cytotoxicity (MTT-based viability assay) and *in vivo* cardiotoxicity (serum CK-MB and LDH activity) were measured after treating with the iboga compounds. The mitochondrial dehydrogenase-based MTT assay was carried out in the C2C12 (mouse skeletal muscle) cell line, which has the potential to differentiate into cardiomyocytes and other cell types. The findings indicated that the potent *Endo*-iboga analogs, **C2** (IC_50_ = 60 µM) and **C4** (IC_50_ = 235 µM), were ∼6 and 1.5-fold more safe than their corresponding natural *Exo*-counterparts, **C5** and **C7**, respectively (**Supporting Figure S1**). However, in this series, **C10** (IC_50_ = 325 µM) turned out to be the safest analog as compared to other iboga congeners.

To determine the cardiotoxic effects of the synthesized iboga compounds, serum CK-MB and LDH activity were measured (**Supporting Figure S2 & S3**). Here also, the potent iboga-analogs (**C2** and **C4**) turned out to be safest, as compared to their natural *Exo*-counterparts (**C5, C7**), showing no significant increase in serum CK-MB and LDH levels. In contrast, **C5** and **C7** showed the elevated serum CK-MB and LDH levels, potentially contributing to the development of cardiomyopathy. Furthermore, a histological study of mice cardiac tissue (Hematoxylin and Eosin stained) was performed to identify any cardiomyopathy associated with **C2** and **C4**, compared to **C5** and **C7** (**Supporting Figure S4**). The histological sample analysis demonstrated the degeneration of the ideal syncytial arrangement of myocytes in the groups treated with **C5** and **C7**, as compared to **C2, C4** and control-treated groups. The cumulative experiments thus highlighted the potential safety of benzofuran-containing iboga epimers, compared to their natural *Exo*-counterparts.

### 2.9. Electrocardiogram (ECG)-based cardiovascular toxicity study in vivo

Since the interaction of natural ibogaine (**C5**) with hERG potassium channels, causes delayed repolarization and prolonged action potential, the direct cardiotoxicity assessment of the synthesized iboga analogs (**C2, C4**, and **C5**) was examined *in vivo* through ECG recordings (**Supporting Figure S5**). The corrected QT interval (QTc, expressed in seconds or milliseconds) for each respective group was measured at both the start (Day 0) and the end (Day 14) of the experiment (**Figure 9**). The experimental result demonstrated that **C5** exhibited the highest QTc (0.154 ± 0.004 s) compared to its untreated control group (0.140 ± 0.005 s), followed by **C2** (0.157 ± 0.006 s) and **C4** (0.137 ± 0.005 s) with their respective controls (0.148 ± 0.009 s for **C2** and 0.135 ± 0.003 s for **C4**) (**Figure 9A**). A more precise statistical analysis showed that the percentage increment in QTc follows the trend: **C5** (10%, \*\*\**P* < 0.001) > **C2** (6.08%, \**P* < 0.05) > **C4** (1.48%, ns). Considering the connection between prolonged QTc intervals and the increased risk of life-threatening cardiac arrhythmias/sudden cardiac death, we may conclude that in our case, **C4** is the safest analog among the series followed by **C2** and **C5**. Indeed, this practical experimentation validates that our bioisosteric modification in the iboga scaffold is worth considering for potential reduction of cardiotoxicity *in vivo*.

**Figure 9:**
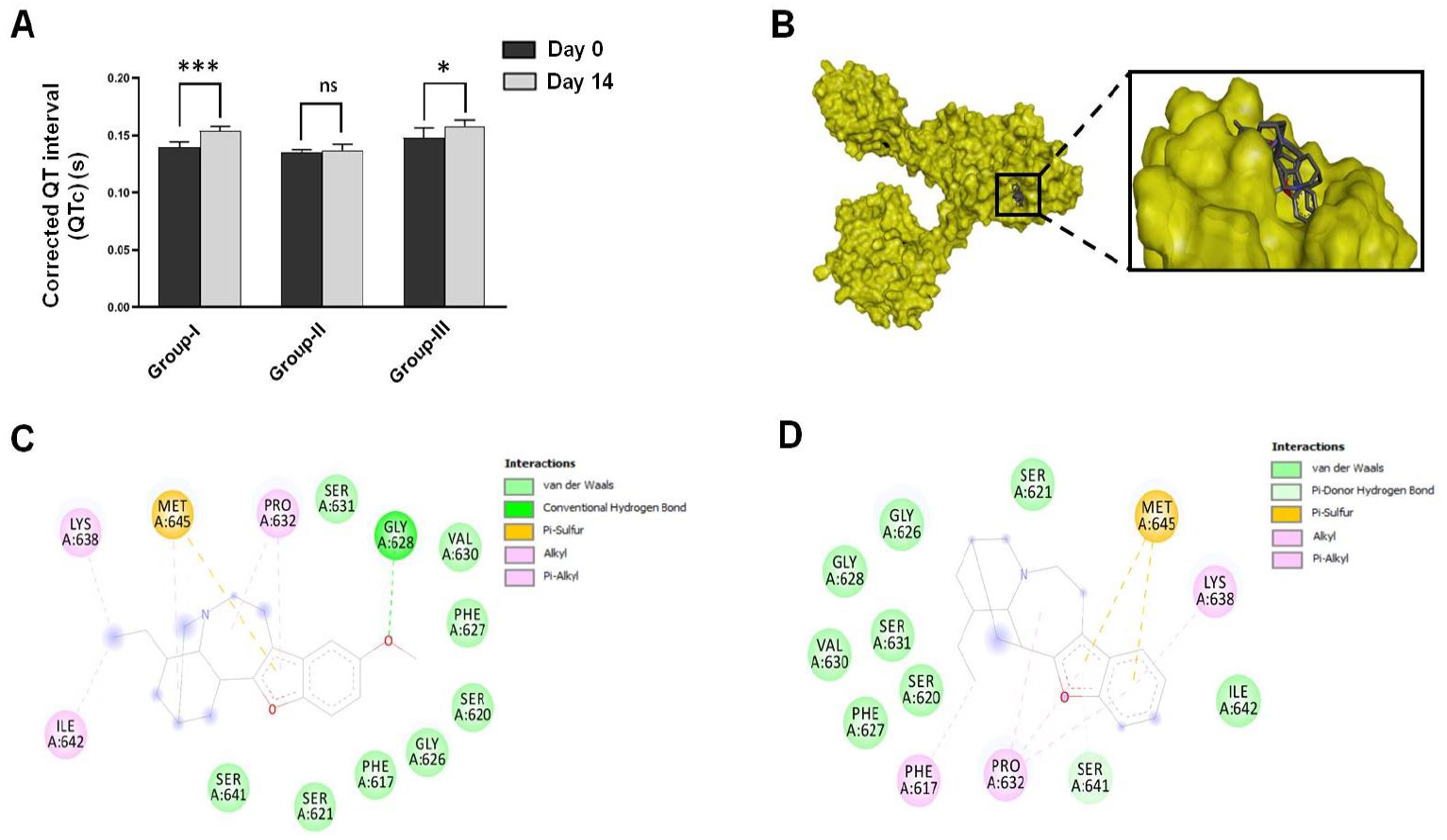
*In vivo* and *in silico* cardiotoxicity assessment of **C2, C4 & C5. (A)**. Corrected QT interval (QTc, measured in seconds) of three different groups of rats at both start (Day 0) and end (Day 14) of experiment. Group-I, Group-II and Group-III represented **C5, C4 & C2**-treated groups, respectively, each experiencing a fixed dose of 15 mg/kg (n = 6). All compounds were administered via conventional oral gavage technique till 14 consecutive days. Data represented as Mean ± SEM. \**P* < 0.05, \*\*\**P* < 0.001, *** indicated significant difference between Day 0 & Day 14 treatment. **(B)**. *In silico* binding of **C2 & C4** with the active site of hERG channel. **(C). & (D)**. represented the interactions of **C2** and **C4** with the amino acid residues of the hERG active site by molecular drug docking.

### 2.10. In silico toxicity prediction

Unlike the interaction between **C5** and S624 residue, our *in silico* study of the synthesized iboga analogs **C2** and **C4** revealed that both compounds lacked the vital hydrogen (H-) bonding interaction with the serine equivalents of S624 residue in hERG channel (**Figure 9C & 9D**). This lack of H-bonding interaction diminished their stabilities in the active site of the hERG channel, providing a possible explanation for why **C2** and **C4** exhibited reduced QTc prolongation in rats. Overall, **C2** formed six hydrophobic interactions with the P632, K638, I642 and M645 residues, and most importantly, one H-bonding interaction with the G628 residue, exerted by its -OMe group. On the contrary, **C4** exhibited the same number of hydrophobic interactions with the F617, P632, K638 and M645 residues, but lacked the above-mentioned H-bonding interaction with the G628 residue due to the absence of the -OMe group. The lack of this H-bonding interaction in **C4** somewhat again destabilized its structure within the active site of the hERG channel, thereby resulting for its higher binding energy (−6.14 kcal/mol) compared to **C2** (−6.49 kcal/mol). Thus, this *in silico* molecular docking study offers a potential rationale for the cardiac safety of our synthesized bioisosteric *Endo*-iboga epimers (particularly **C4**, lacking the H-bond acceptor like OMe).

## 3. Conclusion

Our prior work on the benzofuran-iboga scaffolds with or without tetrahydroazepine ring laid the foundation for their potential use as antinociceptive and anxiolytic agents for acute pain management in mice for the first time [32–34, 37]. These promising results motivated us to design bioisosteric scaffolds of natural ibogaine/ibogamine, ensuring that the scaffold’s biological potency would remain intact while providing stability against light and air, and circumventing the cardiotoxicity associated with the indolyl -NH of the natural scaffold. Consistent with this idea, four different benzofuran-containing iboga-alkaloids had been made and furthermore were individually subjected to biological screening for investigation of their capabilities in ameliorating formalin-induced pain sensation in mice. To our surprise, particularly the *Endo*-analogs of bioisosteric/natural iboga scaffolds (**C2, C4 & C6** and **C8**, respectively) turned out to be effective as compared to their corresponding *Exo*-counterparts (**C1, C3 & C5** and **C7**, respectively). The potency of the *Endo*-analogs was clearly manifested by in terms of both the reduction in the paw licking activity and the elevation in paw withdrawal and tail flick latency in formalin-induced acute pain model in mice. The compromised locomotor activity and anxiety-like behavior were also completely reversed after the administration of **C2 & C4**, and became almost closer to the state of untreated cases. Along with this, significant ROS quenching activity and superior anti-inflammatory effect (manifested by the reduced level of expression of substance P, CGRP, NK1R and p65) of **C2 & C4**, further depicted the potential of the *Endo*-iboga scaffolds. Upon post-treatment of the iboga compounds, the inhibitory (dopamine, GABA) and excitatory neurotransmitter (glutamate and serotonin) were significantly regulated in reciprocal manner with respect to each other. The neurotrophic modulation, as indicated by elevated BDNF expression, was restored to a major extent in case of **C2 & C4**-treatment after formalin administration. Moreover, the *in vitro* compatibility of *Endo*-iboga analogs in the C2C12 cell line and their *in vivo* ability in restoring the serum CK-MB and LDH levels to the extent of the untreated control, without inducing cardiomyopathy, corroborated the safety profile of these compounds over their natural *Exo*-counterparts. Amongst, **C2** and **C4**, the latter showed no significant QTc prologation in rats unlike the case of **C5**, when analyzed by ECG, and thus revealing its cardiac safety. This nonsignificant QTc prolongation might be attributed to the absence of H-bonding interaction between **C4** and the serine equivalents of S624 residue of the active site of hERG channel. In summary, the ease of chemical synthesis, along with the pain-relieving activity and good neuroprotective and cardio safety profile of **C4**, suggest its potential as a clinically useful therapeutic agent. In this context, further development of several newer safer analogs is currently underway, and will be reported in due course.

## 4. Materials and methods

### 4.1. Materials

Phosphate buffer saline, reduced glutathione (GSH), thiobarbituric acid (TBA), trichloroacetic acid (TCA), sodium dodecyl sulfate (SDS), acrylamide, bis-acrylamide, ammonium persulfate, TEMED, glacial acetic acid, glycine, Tris, hydrogen peroxide, dimethyl sulfoxide, copper sulfate, ethanol, Bovine serum albumin (BSA) and all other fine chemicals and reagent were purchased from Merck SA (Darmstadt, Germany). All the mentioned reagents were of molecular biology grade. Mouse specific substance P, CGRP, neurokinin 1 receptor (NK1R), glutamate, γ-Aminobutyric acid (GABA), and serotonin were procured from MyBiosource (San Diego, CA, USA). Primary antibodies for p65, β-actin, H3 and HRP tagged secondary antibodies were procured from Cell Signaling Technology (Danvers, Massachusetts, USA). Primary antibody for BDNF was procured from Novus Biologicals (Centennial, USA). ECL reagent was procured from GE healthcare. Polyvinylidene fluoride (PVDF) membrane was purchased from Merck (Darmstadt, Germany). CK-MB kit (Catalogue No. 1102070210) and LDH kit (Catalogue No. 1102160025) were procured from Coral Clinical Systems, Goa, India. Mouse myoblast cell line (C2C12) was purchased from NCCS cell repository and stored according to the supplier’s protocol.

### 4.2. In vivo experimental study

The Institutional Animal Ethics Committee (IAEC), University of Calcutta, Kolkata, India, which is registered as the Committee for Control and Supervision of Experiments on Animals (CCSEA), a statutory committee of the Department of Animal Husbandry and Dairying (DAHD), Ministry of male Fisheries, Animal Husbandry and Dairying (MoFAH & D), Government of India, approved all experimental protocols and methods. Swiss albino mice, aged 6-8 weeks and weighing 22-25 g, were obtained from a CCSEA registered supplier for the current study. They were housed in standard conditions, with 25 ± 2 ºC and 50% humidity, and were fed and watered freely throughout the day, following a 12-hour light-dark cycle as previously documented in studies on acute inflammatory pain. Before testing, the animals had at least one week of acclimatization. The Institutional Animal Ethics Committee (IAEC), University of Calcutta, Kolkata, India, has duly approved the number of animals used in this study and their care. The standard Animal Research: Reporting of in vivo Experiments (ARRIVE) requirements were adhered to by the researcher for this experimental study in an animal model. A total of 60 animals were randomly assigned to ten experimental groups (n = 6) as follows: **(i) Control:** 20 μL PBS (vehicle) in right hind paw + 70 μL PBS (vehicle) intraperitoneally. **(ii) Formalin:** 20 μL 2.5% (v/v) formalin in right hind paw + 70 μL PBS (vehicle) intraperitoneally. **(iii) Formalin & iboga analog treated groups:** 20 μL 2.5% (v/v) formalin in right hind paw + 70 μL iboga analog at a dose of 30 mg/kg intraperitoneally.

The chronological denotation of the synthesized iboga analogs were mentioned accordingly as depicted above. Soon after the administration of formalin in the right hind paw, the treatments with either PBS or iboga compounds (at a dose of 30mg/kg body weight after dissolution in 50% DMSO-H_2_O) were achieved via IP. Immediately after completion of the treatments, the non-invasive hyperalgesic and spontaneous behaviors of the treated mice were studied. Anxiety-related behaviors, locomotor activity, and allodynic behaviour of the mice were measured from formalin administration upto a period of 24 hours. Following that, all mice were put to death in accordance with CCSEA approved protocols utilizing isoflurane, and hind paw, spinal L4–L6 segments were separated, kept at 80 °C, and used for additional research. For preparation of tissue homogenate, soft tissue from the isolated hind paw of mice was removed and homogenized in RIPA lysis buffer (10mM Tris-HCl, 1mM EDTA, 0.5mM EGTA, 1% Triton X-100, 0.1%sodium deoxycholate, 0.1% SDS, 140mM NaCl, 1mM PMSF - for whole cell lysates), cytosolic extraction buffer (10mM HEPES, 60mM KCl, 1mM EDTA, 0.075% NP-40, 1mM DTT, 1mM PMSF – for cytosolic fraction isolation) and nuclear extraction buffer (20 mM HEPES, 1.5mM MgCl_2,_ 0.42M NaCl, 0.2mM EDTA, 25% v/v glycerol – for nuclear fraction isolation) in presence of protease and phosphatase inhibitors. Finally, homogenates were centrifuged at 4 ºC and 10000 g for 30 mins. In case of preparation of spinal cord lysate, similar protocols were followed with RIPA lysis buffer.

### 4.3. Determination of Tail flick and paw withdrawal latency

To assess the nociceptive responses and determine the antinociceptive activities of iboga analogs, a tail-flick test was performed. The time taken to flick the tail against radiant heat was recorded and represented as tail-flick latency. Three independent tail flicks/mouse were recorded at a given time point and the average values were presented. The hot plate test assessed the sensorimotor activity of each of the mice during painful stimuli when placed on a metal test plate preheated to 52 ± 1 °C. Paw withdrawal latency was recorded as the time elapsed until the mouse licked or flicked its hind paw.

### 4.4. Paw licking behavior study

Mouse paw licking behavior was recorded in phase 1 (up to 10 min) and phase 2 (10 min onwards) after formalin infusion. The recorded events were measured as licking per minute for each mouse. Thus, the antinociceptive behavior of iboga analogs was recorded in both the first and second phases of formalin-induced nociception as reported earlier.

### 4.5. Von Frey monofilament test

The Von Frey monofilament test was used to determine the tactile sensory threshold of the hind limb of mice to examine the pain-induced allodynic response and its inhibition by iboga analogs. The mice were housed on a transparent plexiglass cylinder-enclosed wired mesh table for fifteen minutes before the experiment. After confirming that the mice were quiet, Von Frey filaments were applied to the lateral portion of the paw, starting at 0.001 g and going up to 4 g and the subsequent reaction of mice were recorded.

### 4.6. Study of locomotor activity in open field test (OFT)

The OFT in a circular arena with closed walls was performed following the specifications as reported by others with slight modifications [34]. The animals were set free to the circular open field arena where the field of study was made by opaque plastic materials with a circular (40 cm in radius) field having a 17 cm high continuous wall in the periphery. During study, mice were placed at the center and allowed to explore the field for 5 minutes. The exploratory activity of mice was video recorded using the ANY-Maze software for the central region of the mice. For locomotor activity, the average speed and distance travelled in the open field was evaluated before and after specific treatments.

### 4.7. Evaluation of anxiety-like behavior in elevated plus maze (EPM) test

The EPM tests were performed following the protocols described in earlier studies with suitable modifications. The apparatus was made up of wood with two open arms (30 × 5 cm each) and two closed arms (30 × 5 × 15 cm each) arranged in such a manner that the two arms of each type were opposite to each other. The maze was elevated 40 cm above the floor with wooden legs. Each animal was placed at the center of the maze facing any of the open arms. Through a test period of 5 minutes, the exploration of mice was video recorded considering the central point by ANY-Maze software. The number of entries into the open arms and the time spent in the open arms were automatically counted by the software as the measures of the anxiety-like behavior following treatments.

#### 4.7.1. Estimation of reduced glutathione (GSH) level

The sulfhydryl group of GSH reacts with DTNB and produces a yellow colored 5-thio-2-nitrobenzoic acid (TNB). Measurement of the absorbance of TNB provides accurate estimation of GSH in the sample. After applying 0.1 mL of 25% TCA to the paw homogenate, the precipitate was centrifuged at 3900 g for 10 minutes to pellet the material. By mixing 1 mL of the supernatant with 2 mL of 0.5 mM DTNB, produced in 0.2 M phosphate buffer (pH 8), the free endogenous sulfhydryl group was measured in a total volume of 3 mL. Following a reaction, the GSH and DTNB created a yellow complex and thereafter the absorbance was measured at 412 nm.

#### 4.7.2. Catalase activity

Catalase enzyme activity was determined by monitoring the decrease in absorbance resulting from the elimination of H_2_O_2_ by the action of catalase. Absorbance of H_2_O_2_ was measured at 240 nm to measure the catalase activity. The absorbance was then measured after the homogenate was added, showing that the catalase had eliminated the H_2_O_2_. Paw tissue homogenate, 100 mM hydrogen peroxide, and 50 mM potassium phosphate buffer (pH 7.0) were combined in a 1.0 mL reaction volume. After that, the reaction was conducted at 20 ºC, and the catalase activity was calculated using only the first linear rate of absorbance.

#### 4.7.3. SOD 1 activity

The auto-oxidation of pyrogallol was used to evaluate the activity of SOD 1. Pyrogallol’s auto-oxidation was modified during computation. To put it briefly, the paw tissue homogenate was mixed with 62.5 mM tris-cacodylic acid solution and then treated with 4 mM pyrogallol. The initial absorbance of pyrogallol was measured at 420 nm, followed by the absorbance of test samples taken at predetermined intervals at the same wavelength, to track the auto-oxidation of pyrogallol.

#### 4.7.4. Estimation of lipid peroxidation

Thiobarbituric acid reactive substances (TBARS) such as malondialdehyde, formed from the breakdown of polyunsaturated fatty acids, serves as an index for determining the extent of lipid peroxidation level. To ascertain the cellular stress, the generation of thiobarbituric acid reactive substance (TBARS) in the homogenate was quantified using a standard technique. In summary, the homogenate was combined with 15% TCA, 0.35% TBA, and 5 N HCl and incubated for 15 minutes at 95 ºC. Thereafter, following chilling, all the mixes underwent centrifugation, and the supernatant’s absorbance was measured at 535 nm using a suitable blank. Using an assumed value of e = 1.56 × 10^5^ /M/cm, the quantity of lipid peroxidation in each sample was represented in terms of TBARS in nanomoles/gm tissue.

### 4.8. Immunoblot assay

For 10% sodium dodecyl sulphate-polyacrylamide gel electrophoresis (SDS-PAGE), an equivalent amount of protein (40 µg) was added in each lane and transferred to a PVDF membrane. A 5% bovine serum albumin (BSA) solution was used to block the membrane for an entire night at 4 ºC. Monoclonal antibodies against mouse p65, COX-2, TNF-α, IL-6, BDNF, and GDNF protein were used for immunoblotting. For the cytosolic and nuclear extracts, loading controls of β-actin and histone 3 (H3) were employed, respectively. The BioRad Chemidoc (MP imaging system) was utilized to capture the immunoblots, and Molecular Analyst version 1.5 (Bio-Rad Laboratories, Hercules, CA) and Image J (NIH) software were used for normalization and analysis.

### 4.9. Estimation of neurotransmitter

Levels of substance P, CGRP, NK1-R, glutamate, GABA, dopamine, and serotonin were determined from spinal L4-L6 segment lysates using sandwich ELISA techniques following manufacturer’s protocols (MyBiosource, USA). The respective catalogue numbers were as follows: MBS8800512, MBS8800040, MBS7606597, MBS2601720, MBS260709, MBS8807516 and MBS1601042.

### 4.10. In vitro cytotoxicity assay

C2C12 cells were maintained in Dulbecco’s Modified Eagle’s Medium (DMEM) and supplemented with 10% FBS and streptomycin and penicillin. Cells were kept at 37°C in a humidified atmosphere containing 5% CO_2_. Cells were harvested with 0.25% trypsin (Gibco) after reaching 85-90% confluency for 2–3 min at 37 °C and counted before seeding. Cells were then seeded in 96-well plates at a density of 5000 cells/well and incubated for 24 h. After incubation, cells were then treated with different concentrations of **C2, C4, C5, C7**, and **C10** in presence of 0.5% serum and incubated for 96 h. Freshly prepared MTT solution in media was then added and incubated for another 4 h. After dissolution of the formazan crystals in DMSO, the corresponding absorbance was recorded by a BioTek plate reader at 570 nm. Results were analyzed by GraphPad Prism 6.0 software.

### 4.11. CK-MB and LDH activity

Serum CK-MB (creatine kinase myocardial band) activity was measured to find iboga analogs-induced cardiomyopathy following the manufacturer’s protocol provided with the CK-MB kit [54]. This immunoinhibition method followed the anti-agglutinating sera mediated inhibition of CK-M fraction of CK-MM and CK-MB. Then the CK-B fraction was measured by the CK (NAC act.) method. Finally, the CK-MB activity was calculated by multiplying the CK-B fraction by two. Values were expressed in U/L after calculation.

Serum LDH (lactate dehydrogenase) activity was measured to investigate iboga analogs-induced cardiotoxicity, following the manufacturer’s protocol provided with the LDH kit [55]. LDH enzyme in the serum sample catalyzes reduction of pyruvate in presence of NADH to generate NAD. This contemporary oxidation of NADH to NAD was measured by a decrease in absorbance (340 nm) proportional to the serum LDH activity.

All the respective iboga compounds were injected in mice via IP at a dose of 30 mg/kg body weight as previously described in section 4.2. The results were analyzed by GraphPad Prism 6.0 software.

### 4.12. Histopathological studies

For histopathological studies, animals were perfused after post-treatment with respective compounds at an equivalent dose of 30 mg/kg body weight and then fixed with PBS and neutral buffered formalin (10% formaldehyde in saline) followed by immediate washing and fixing of cardiac tissue in neutral buffered saline. The tissue was then embedded in the paraffin wax for block preparation following established protocols. 5 μm thick tissue sections were cut using a rotary microtome (Leica Biosystems) for Hematoxylin and eosin (H&E) staining. Myocardial damages were studied by observing the degeneration of the cardiac myocyte cells from standard striation and functional syncytium like arrangements [54]. All images were captured at an optical magnification of 400X using an Olympus bright field microscope. For visual interpretation of images, at least three images per group were considered.

### 4.13. ECG-Based Cardiotoxicity Assessment In Vivo

All the involved animal experiments were approved by the IAEC of the Department of Pharmaceutical Technology, Jadavpur University. Male Wistar albino rats (8-9 weeks), weighing 130-150 g, were purchased from CCSEA registered supplier (West Bengal, India). To rule out any concurrent infections, rats were monitored for about 10-14 days, before the start of the experiment. The animals were kept in polypropylene cages with well-aerated stainless-steel lids, and subjected to a 12-hour light/dark cycle and ambient temperature typically around 20-24 °C and 40-60% humidity and were fed and watered freely throughout the day. After acclimatization period, rats were divided in three groups (n=6), i.e. **(1) Group-I** (administered with **C5**), **(2) Group-II** (administered with **C4**) & **(3) Group-III** (administered with **C2**). All respective iboga compounds were administered by oral gavage technique at a dose of 15 mg/kg body weight till 14 consecutive days. The rats were anesthetized with ketamine (60 mg/kg) and xylazine (10 mg/kg), as previously reported [56]. An ECG was obtained immediately after anesthesia using a conventional lead for 5 minutes. The ECG signals were collected and analyzed using a BIOPAC MP36 (Goleta, CA, USA). The QT and RR-interval were measured at both the start (Day 0) and end (Day 14) of experiment. For each respective case, QTc (expressed in seconds or milliseconds) was calculated by Bazett’s formula, QTc = QT/√RR

### 4.14. In silico toxicity assessment

The *in silico* toxicity prediction of the synthetic bioisosteric *Endo*-iboga epimers (**C2 & C4**) was assessed by their binding affinities towards hERG protein [57]. The macromolecule with PDB ID: 5VA1 was obtained from the Protein Data Bank (https://www.rcsb.org). The binding conformations of the receptor and ligands were predicted using the molecular docking software AutoDockTools 1.5.6. The docking protocol was established based on previous protocol, with minor modifications [58]. The center and dimensions of the gridbox were determined by calculating the center of the hERG channel and the distance between the two most distant residues, derived from the coordinates of key residues (T623, S624, V625, G648, Y652, and F656) across the four chains. The grid-box was configured to 45 × 45 × 45 with a spacing of 0.5 Å, centered at X = 73.15, Y = 73.15, and Z = 88.90. Additionally, the characteristics of the protein-ligand complex and their mode of interactions were analyzed by Discovery Studio Visualizer [59].

### 4.15. Data analysis and statistics

All data points were displayed as Mean ± SEM. Using Origin Lab 8.5 and GraphPad Prism 6.0, the One-Way and Two-way ANOVA were used to examine the significance of differences between the treated and untreated groups after Tukey’s test. A *P*-value of less than 0.05 was considered significant.

## Author Contributions

S.S & S.D: Conceived the idea, designed the hypothesis and edited the manuscript; A.G: Synthesized the chemical compounds with proper characterizations, performed the HPLC studies and wrote the manuscript; T.B: Executed the antinociceptive and anxiolytic evaluation and wrote the manuscript; S. Pratihar: Assisted in the initial synthesis of compounds; S. Parvage: Performed serum CK-MB & LDH activity and histological analysis of cardiac tissues; S.N.S: Evaluated the *in vitro* cytotoxicity study; A.S & A.D: Executed the molecular docking study and cardiotoxicity evaluation *in vivo*; S.D: Supervised and analysed the cardiotoxicity assessment study and edited the manuscript.

## Acknowledgement

A.G., S. Pratihar and S.N.S. thank IACS for their fellowships; T.B & S. Parvage are thankful to DRDO-LSRB & CSIR, respectively for their fellowships; A.S & A.D jointly thank State Government and ICMR, respectively for their fellowships. All authors have given approval to the final version of the manuscript.

## Associated Content

All synthetic protocols, characterization of compounds, associated biological experiments and raw data of whole western blot are provided in the supplementary information.

