## Supplemental Information for "Development of Bioisosteric *Iboga*-alkaloids as Antinociceptive and Anxiolytic Agents with Neuroprotective Effects"

¶ A.G & T.B contributed equally

### Table of Contents

Page No

General Information ………………………………………………………………. 3

Chemical synthesis for bioisosteric ibogaine & ibogamine analogs ………………...... 4 -10

NMR ($^{1}H$, $^{13}C$, DEPT-135) ………………………………………………….............. 11-28

HRMS (ESI, +ve ion mode) …………………………………………………........... 29-33

HPLC of **C1-C4** …………………………………………………………………… 34

**Figure S1**: MTT-based cellular viability assay………………………......……….. 35

**Figure S2**: Serum CK-MB level study…………………………………………..... 36

**Figure S3**: Serum LDH activity study……………………………………………... 36

**Figure S4**: Cardiotoxicity evaluation from histological samples of heart……......... 37

Whole Western blot images …………………………………………….............. 38

**Figure S5**: ECG-based cardiovascular toxicity assessment of **C2**, **C4** & **C5** ……….. 39

#### General Information

All reagents were purchased from commercial sources (Sigma-Aldrich/SRL/BLDpharm) and utilized without further purification unless specified. All reactions were performed on oven-dried glassware under inert atmosphere condition wherever necessary. Solvents were dried carefully following standard protocols prior to setting up reactions. Distilled solvents were generally used for column chromatography unless otherwise mentioned. Thin layer chromatography (TLC) was performed on sheets of silica gel 60 F254 on aluminium (layer thickness 0.25 mm, Merck). Visualization of TLC spots was achieved in presence of UV light after staining with Ceric Ammonium Molybdate (CAM) or Ninhydrin or Phosphomolybdic Acid (PMA) or Potassium Permanganate ($KMnO_4$) solution as required. All chromatographic separations were accomplished by using silica gels of 60-120/100-200/230-400 mesh with suitable eluents (e.g. MeOH-DCM/Acetone-DCM/EtOAc-Petroleum Ether). Elucidation of the chemical structure of compounds was made by recording NMR spectroscopy in Bruker NMR spectrometers (300 MHz for $^{1}$H and 75 MHz for $^{13}$C or DEPT characterization) using $CDCl_3$ as solvent. Chemical shifts (δ) were measured in ppm with respect to the solvent residual peak or TMS (being present in the solvent as internal standard). The following abbreviations are used to designate multiplicity of NMR signals: s = singlet; d = doublet; t = triplet; q = quartet, dd = doublet of doublet, m = multiplate; br = broad. High resolution mass spectrometry (HRMS) was performed in QTOF I (quadrupole-hexapole-TOF) mass spectrometer with an orthogonal Z-spray-electrospray interface on Micro (YA-263) mass spectrometer (Manchester, UK). Infrared spectra (IR) were recorded on Shimadzu FTIR-8300 spectrometer. High Performance Liquid Chromatography (HPLC) was done on Shimadzu SP-20AD system by Analytical RP $C_{18}$ column. All NMR spectra and HRMS data were processed with MestReNova software. For cell culture, reagents from Invitrogen and HI-MEDIA were used and cell culture dishes were obtained from BD Falcon.

**Triethyl(2-(2-((triethylsilyl)oxy)benzofuran-3-yl)ethoxy)silane (1a):**

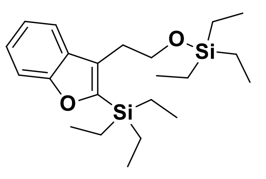

**1a**

To a stirred solution of 2-Iodophenol (1.5 g, 6.82 mmol) in dry DMF (10 ml) was added Triethyl((4-(triethylsilyl)but-3-yn-1-yl)oxy)silane (2.1 g, 6.82 mmol) and was degassed thoroughly for 15 min. $Na_2CO_3$ (3.61 g, 34.1 mmol) and LiCl (261 mg, 6.14 mmol) were then added followed by addition of $Pd(OAc)_2$ (153 mg, 0.68 mmol) and heated at 90°C for 3 h. The reaction mixture was then passed through a pad of celite and extracted with EtOAc (70 ml) and water (30 ml). The aqueous layer was extracted with another 50 ml of EtOAc and the combined organic extract was then collected over anhydrous $Na_2SO_4$ and concentrated in vacuo. The crude product containing both **1a** and **1b** was next purified on silica gel (230-400 mesh) eluting initially with 5% EtOAc in Petroleum Ether to afford **1a** (612 mg, 23%) and then gradually raised 10% EtOAc in Petroleum Ether to afford **1b** (1.45 g, 77%) as deep brown oil. ***TLC→ (EtOAc:Petroleum Ether = 1:19 v/v); $R_f$ = 0.7 (for 1a) & 0.3 (for 1b)***

Compound **1b** (1 g, 3.62 mmol) was dissolved in dry DCM (15 ml) and settled at 0°C during the addition of TESCl (0.67 ml, 3.98 mmol) and imidazole (295 mg, 4.34 mmol). After complete consumption of starting material, the reaction mixture was extracted with DCM (50 ml) and water (25 ml). The organic layer collected over anhydrous $Na_2SO_4$ was then concentrated. The crude product was purified as mentioned early to afford **1a** (1.31 g, 93%) as deep brown oil.

**$^1$H NMR (300 MHz, $CDCl_3$):** δ 7.60 – 7.54 (m, 1H), 7.43 (dt, J = 8.3, 0.8 Hz, 1H), 7.27 – 7.13 (m, 2H), 3.83 (dd, J = 8.3, 7.1 Hz, 2H), 3.06 – 2.96 (m, 2H), 1.05 – 0.98 (m, 9H), 0.98 – 0.89 (m, 15H), 0.63 – 0.57 (m, 6H). **$^{13}$C NMR (75 MHz, $CDCl_3$):** δ 158.10, 157.12, 129.38, 127.55, 124.25, 121.87, 119.84, 111.36, 63.48, 28.67, 7.49, 6.84, 4.50, 4.47, 3.69. **DEPT-135 NMR (75 MHz, $CDCl_3$):** δ 124.32, 121.93, 119.90, 111.42, 63.55, 28.74, 7.55, 6.91, 4.57, 3.75. **IR ($CHCl_3$, $cm^{-1}$):** 2953, 2911, 2875, 1551, 1447, 1414, 1234, 1096, 1004, 808, 736, 721, 610. **HRMS (ESI, + ve ion mode) (m/z):** $[M+Na]^+$ for $C_{22}H_{38}O_2Si_2$: Calcd. 413.2308; Found 413.2154

**Triethyl(2-(2-iodobenzofuran-3-yl)ethoxy)silane (2) :**

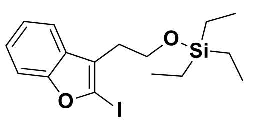

**2**

To a stirred solution of compound **1a** (1 g, 2.56 mmol) in dry $CH_3CN$ (15 ml) was added *N*-Iodosuccinimide (690 mg, 3.1 mmol) and left for overnight. The reaction mixture was then quenched with saturated $Na_2S_2O_3$ solution (10 ml) and concentrated on vacuo and then extracted with EtOAc (50 ml) and water (30 ml). After collecting over anhydrous $Na_2SO_4$, the organic layer was then concentrated in vacuo. The crude product was next purified on silica gel (230-400 mesh) eluting with 5% EtOAc in Petroleum Ether to afford **2** (772 mg, 75%) as yellow oil. ***TLC→ (EtOAc:Petroleum Ether = 1:19 v/v); $R_f$ = 0.8***

**$^1$H NMR (300 MHz, $CDCl_3$):** δ 7.51 (dd, *J* = 5.9, 3.2 Hz, 1H), 7.41 (dd, *J* = 6.0, 3.2 Hz, 1H), 7.19 (dd, *J* = 6.0, 3.2 Hz, 2H), 3.81 (td, *J* = 7.2, 1.2 Hz, 2H), 2.86 (td, *J* = 7.2, 1.0 Hz, 2H), 0.92 (td, *J* = 7.9, 1.3 Hz, 9H), 0.57 (dd, *J* = 8.8, 7.5 Hz, 6H). **$^{13}$C NMR (75 MHz, $CDCl_3$):** δ 158.21, 128.68, 124.48, 124.21, 122.80, 119.01, 110.99, 98.35, 61.96, 29.70, 7.50, 6.86, 4.50, 3.65. **DEPT-135**

**NMR (75 MHz, CDCl$_3$):** δ 124.13, 122.72, 118.93, 110.90, 61.88, 29.62, 6.78, 4.42. **IR (CHCl$_3$, cm$^{-1}$):** 2940, 2905, 2840, 1570, 1456, 1234, 1089, 1029, 978, 808, 736, 705, 602. **HRMS (ESI, + ve ion mode) (m/z):** $[M+H]^+$ for $C_{16}H_{23}IO_2Si$: Calcd. 403.0590; Found 403.0596

**2-(2-Iodobenzofuran-3-yl)ethanol (3) :**

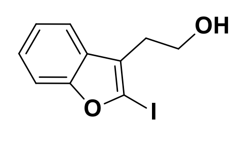


**3**

To a stirred solution of compound **2** (1.5 g, 3.74 mmol) in dry THF (10 ml) was added Tetrabutylammonium fluoride (TBAF) solution (1M in THF) (5.61 ml, 5.61 mmol) and left overnight. Organic solvent was then removed and extracted with EtOAc (50 ml) and water (30 ml). The organic extract was collected over anhydrous $Na_2SO_4$ and concentrated under reduced pressure. The crude product containing was then purified on silica gel (100-200 mesh) eluting initially with 25% EtOAc in Petroleum Ether to afford **3** (740 mg, 69%) as yellow oil. ***TLC→ (EtOAc:Petroleum Ether = 3:7 v/v); $R_f$ = 0.5***

**$^1$H NMR (300 MHz, CDCl$_3$):** δ 7.46 – 7.42 (m, 1H), 7.39 – 7.34 (m, 1H), 7.16 – 7.11 (m, 2H), 3.75 (t, *J* = 6.8 Hz, 2H), 2.78 (t, *J* = 6.8 Hz, 2H), 2.75 – 2.70 (m, 1H). **$^{13}$C NMR (75 MHz, CDCl$_3$)**: δ 157.99, 128.20, 124.23, 123.82, 122.83, 118.63, 110.93, 98.61, 61.42, 29.20. **DEPT-135 NMR (75 MHz, CDCl$_3$):** δ 124.40, 123.00, 118.79, 111.10, 61.59, 29.36. **IR (CHCl$_3$, cm$^{-1}$):** 3580, 3336, 2950, 2878, 1581, 1452, 1275, 1183, 1090, 1044, 1008, 930, 856, 799, 740, 693. **HRMS (ESI, + ve ion mode) (m/z):** $[M+H]^+$ for $C_{10}H_9IO_2$: Calcd. 288.9725; Found 288.0828

**2-Iodo-3-(2-iodoethyl)benzofuran (4) :**

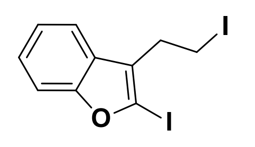


**4**

To a stirred solution of compound **3** (900 mg, 3.12 mmol) in dry DCM (10 ml) was added $PPh_3$ (984 mg, 3.75 mmol), $I_2$ (951 mg, 3.75 mmol) and imidazole (425 mg, 6.25 mmol) at 0°C and then gradually warmed to r.t. for additional stirring for 4 h. The reaction mixture was then quenched with saturated $Na_2S_2O_3$ solution (10 ml) and extracted with DCM (40 ml) and water (30 ml). The organic extract was collected over anhydrous $Na_2SO_4$ and concentrated under reduced pressure. The crude product containing was then purified on silica gel (230-400 mesh) eluting with 1.5% EtOAc in Petroleum Ether to afford **4** (1.1 g, 91%) as yellow oil. ***TLC→ (EtOAc:Petroleum Ether = 2.5:97.5 v/v); Rf = 0.9***

**$^1$H NMR (300 MHz, CDCl$_3$):** δ 7.53 – 7.39 (m, 2H), 7.25 – 7.17 (m, 2H), 3.35 (ddd, *J* = 8.0, 7.1, 1.2 Hz, 2H), 3.20 (ddd, *J* = 8.6, 7.2, 1.2 Hz, 2H). **$^{13}$C NMR (75 MHz, CDCl$_3$):** δ 158.21, 141.77, 127.54, 126.27, 124.57, 124.55, 123.08, 122.64, 119.25, 118.37, 111.76, 111.28, 98.77, 30.36, 28.58, 3.83, 2.38. **DEPT-135 NMR (75 MHz, CDCl$_3$):** δ 124.54, 123.12, 123.07, 122.63, 119.24, 118.35, 111.75, 111.27, 30.34, 28.56, 3.82, 2.37. **IR (CHCl$_3$, cm$^{-1}$):** 3052, 2923, 2844, 1583, 1478, 1452, 1433, 1273, 1171, 1093, 1026, 856, 739, 694, 524, 500. **HRMS (ESI, + ve ion mode) (m/z):** $[M+Na]^+$ for $C_{10}H_8I_2O$: Calcd. 398.8743; Found 398.8745

**7-Ethyl-2-(2-(2-iodobenzofuran-3-yl)ethyl)-2-azabicyclo[2.2.2]oct-5-ene (*Exo* + *Endo*)**

**(6a & 6b) :**

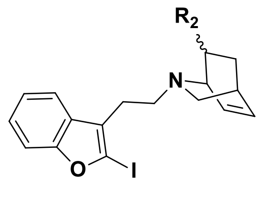

$R_2$ = - Et (*Exo*), **6a**
= - Et (*Endo*), **6b**

The -Cbz deprotection of compound **5** (700 mg, 2.85 mmol) was achieved with the usage of ~33% HBr in AcOH (1.55 ml) dissolved in dry DCM (5 ml). After complete consumption of starting material, the solution was concentrated and repetitively co-evaporated DCM:Petroleum Ether (1:4, 50 ml) to evaporate residual AcOH. Then to the resulting crude compound was added dry $CH_3CN$ (8 ml) followed by addition of $Cs_2CO_3$ (1.4 g, 4.37 mmol) and compound **4** (610 mg, 1.53 mmol) dissolved in dry $CH_3CN$ (5 ml) and refluxed for 12 h. The reaction mixture was then evaporated to dryness and extracted with EtOAc (30 ml) and water (10 ml). The organic extract was collected over anhydrous $Na_2SO_4$ and concentrated under reduced pressure. The crude product was then purified on silica gel (230-400 mesh) eluting with gradually varying percentage of EtOAc (2.5-10%) in Petroleum Ether to afford the pure product containing both **6a** & **6b** (610 mg, 69%) as yellow oil. The characterization of the compounds was made by HRMS and was directly used in the next step.

**HRMS (ESI, + ve ion mode) (m/z):** $[M+H]^+$ for $C_{19}H_{22}NOI$: Calcd. 408.0824; Found 408.0725

**7-Ethyl-6,6a,7,8,9,10,12,13-octahydro-6,9-methanobenzofuro[2,3-d]pyrido[1,2-a]azepine (*Exo* + *Endo*) (C3** & **C4) :**

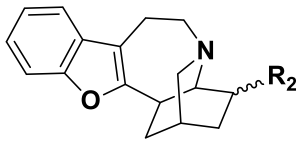

$R_2$ = - Et (*Exo*), **C3**
= - Et (*Endo*), **C4**

To a stirred solution of **6a** & **6b** (200 mg, 0.5 mmol) in dry DMF (5 ml) was added $PPh_3$ (26 mg, 0.1 mmol) and HCOONa (135 mg, 1.96 mmol) and was degassed thoroughly for 15 min. $Pd(OAc)_2$ (11 mg, 0.05 mmol) was then added followed by addition heating at 90°C for 12 h. The reaction mixture was then passed through a pad of celite and extracted with EtOAc (50 ml) and water (20 ml). The aqueous layer was extracted with another 30 ml of EtOAc and the combined organic extract was then collected over anhydrous $Na_2SO_4$ and concentrated in vacuo. The crude product containing both **C3** and **C4**, was next purified on silica gel (230-400 mesh) eluting initially with gradually varying percentage of EtOAc (15% to 30%) in Petroleum Ether to afford the pure products, i.e., **C3** (40 mg, 30%) and **C4** (58 mg, 42%) as deep yellow oil.

***Exo*-product (C3): $^1$H NMR (300 MHz, $CDCl_3$):** δ 7.60 – 7.54 (m, 1H), 7.49 – 7.43 (m, 2H), 7.25 (ddd, *J* = 9.8, 6.6, 1.6 Hz, 1H), 3.86 (qd, *J* = 6.8, 1.6 Hz, 1H), 3.08 – 2.71 (m, 7H), 2.23 (dq, *J* = 11.4, 1.9 Hz, 1H), 2.00 (d, *J* = 11.1 Hz, 1H), 1.83 – 1.65 (m, 2H), 1.65 – 1.47 (m, 2H), 1.43 – 1.29 (m, 1H), 1.28 – 0.96 (m, 4H).**$^{13}$C NMR (75 MHz, $CDCl_3$):** δ 155.31, 128.39, 124.20, 122.34, 119.73, 118.60, 111.51, 103.94, 66.96, 56.76, 56.27, 54.79, 51.63, 35.45, 34.24, 29.04, 26.32, 24.95, 22.91, 17.78. **DEPT-135 NMR (75 MHz, $CDCl_3$):** δ 124.25, 122.38, 119.77, 111.55, 103.98, 67.00, 56.81, 56.31, 54.83, 51.68, 35.49, 29.08, 26.36, 25.00, 22.95, 17.82. **IR ($CHCl_3$, $cm^{-1}$):** 2925, 2854, 2837, 1664, 1582, 1453, 1361, 1274, 1150, 1090, 1081, 1008, 857, 744. **HRMS (ESI, + ve ion mode) (m/z):** $[M+H]^+$ for $C_{19}H_{23}NO$: Calcd. 282.1858; Found 282.1851

***Endo*-product (C4): $^{1}$H NMR (300 MHz, CDCl$_3$):** δ 7.68 – 7.60 (m, 1H), 7.53 – 7.45 (m, 1H), 7.33 – 7.20 (m, 2H), 3.98 – 3.80 (m, 1H), 3.57 – 3.45 (m, 1H), 3.36 – 3.16 (m, 5H), 3.05 (d, *J* = 11.2 Hz, 1H), 2.96 – 2.80 (m, 1H), 2.35 (dd, *J* = 38.2, 11.1 Hz, 2H), 2.01 (d, *J* = 11.1 Hz, 2H), 1.83 (t, *J* = 12.5 Hz, 1H), 1.74 – 1.62 (m, 1H), 1.55 – 1.42 (m, 1H), 1.19 (dd, *J* = 37.7, 4.7 Hz, 3H). **$^{13}$C NMR (75 MHz, CDCl$_3$):** δ 155.27, 127.29, 124.60, 122.76, 119.48, 115.90, 111.55, 102.39, 70.89, 65.82, 55.26, 54.81, 54.66, 52.45, 32.64, 30.84, 29.64, 29.27, 27.38, 24.70, 23.32, 19.88, 17.25. **DEPT-135 NMR (75 MHz, CDCl$_3$):** δ 124.74, 122.90, 119.63, 111.69, 102.53, 65.96, 55.40, 54.95, 54.80, 52.59, 32.78, 30.97, 27.52, 24.84, 23.46, 20.02, 17.40. **IR (CHCl$_3$, cm$^{-1}$):** 2940, 2860, 2832, 1652, 1576, 1440, 1345, 1285, 1189, 1070, 1085, 1008, 860, 742. **HRMS (ESI, + ve ion mode) (m/z):** $[M+H]^+$ for $C_{19}H_{23}NO$: Calcd. 282.1858; Found 282.1839

**2-Iodobenzene-1,4-diol (8) :**

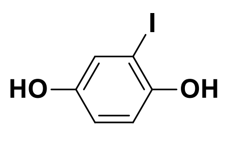

**8**

To a stirred solution of hydroquinone (2.3 g, 20.89 mmol) in dry DMF (15 ml) was added TBDMSCl (6.61 g, 43.86 mmol) and imidazole (3 g, 43.86 mmol) at 0°C and then gradually warmed to r.t. for additional stirring for 4 h. The reaction mixture was then diluted with EtOAc (80 ml) and water (30 ml). The aqueous layer was further extracted with EtOAc (50 ml) and the combined organic phase was collected over anhydrous $Na_2SO_4$ and evaporated to dryness. The crude product (6.93 g, 98%) was directly used in next step without any further purification.

The di-silylated product (7.2 g, 21.26 mmol) was dissolved in dry $CH_3CN$ (30 ml) and to this solution was added *N*-Iodosuccinimide (5.26 g, 23.39 mmol) and TFA (0.65 ml, 8.5 mmol) and allowed to stir for 12 h.The reaction mixture was then quenched with saturated $Na_2S_2O_3$ solution (20 ml) and concentrated in vacuo and then extracted with EtOAc (50 ml) and water (30 ml). After collecting over anhydrous $Na_2SO_4$, the organic layer was then concentrated in vacuo to afford the crude compound (7.4 g, 74%) which was directly used in the next step without any further purification.

To a stirred solution of the above-mentioned crude compound (3.5 g, 7.53 mmol) in dry THF (30 ml) was added tetrabutylammonium fluoride (TBAF) solution (1M in THF) (15.82 ml, 15.82 mmol) and left overnight. Organic solvent was then removed and extracted with EtOAc (50 ml) and water (30 ml). The organic extract was collected over anhydrous $Na_2SO_4$ and concentrated under reduced pressure. The crude product containing was then purified on silica gel (100-200 mesh) eluting with 30% EtOAc in Petroleum Ether to afford the product **8** (1.4 g, 78%) as slight orange-white solid. ***TLC→ (EtOAc:Petroleum Ether = 3:7 v/v); $R_f$ = 0.5***

**$^{1}$H NMR (300 MHz, CDCl$_3$):** δ 7.15 (d, *J* = 2.8 Hz, 1H), 6.77 (d, *J* = 8.7 Hz, 1H), 6.70 (dd, *J* = 8.7, 2.7 Hz, 1H). **$^{13}$C NMR (75 MHz, CDCl$_3$)**: δ 150.42, 149.09, 124.78, 117.08, 115.28, 84.66. **DEPT-135 NMR (75 MHz, CDCl$_3$):** δ 124.81, 117.11, 115.30. **HRMS (ESI, + ve ion mode) (m/z):** $[M+H]^+$ for $C_6H_5IO_2$: Calcd. 236.9412; Found 236.9413

**2-(2-Iodo-5-methoxybenzofuran-3-yl)ethanol (9) :**

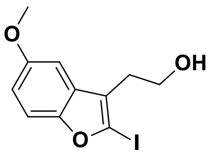

9

Larock heteroannulation was performed as mentioned earlier starting from the ortho-iodinated hydroquinol (1 g, 4.24 mmol) to yield the mono-silylated (224 mg, 13%) and di-silylated (805 mg, 65%) products. All the free -OH groups of the mixture of mono- & di-silylated products were subjected to: (i) protection with triethylsilyl group, followed by (ii) ipso iodination with NIS, (iii) TBAF-mediated removal all silyl protecting groups according to earlier mentioned protocol. Then to the crude compound (800 mg) dissolved in dry DMF (4 ml), was added $K_2CO_3$ (730 mg, 5.26 mmol) and MeI (0.5 ml, 7.9 mmol) and left to stir for 4 h. The reaction mixture was then extracted with EtOAc (50 ml) and water (30 ml). The aqueous layer was further extracted with EtOAc (20 ml) and the combined organic phase was collected over anhydrous $Na_2SO_4$ and evaporated to dryness. The crude product was purified on silica gel (100-200 mesh) eluting with 20% EtOAc in Petroleum Ether to afford **9** (512 mg, 38% over 5steps) as red-brown liquid. ***TLC→ (EtOAc:Petroleum Ether = 3:7 v/v); $R_f$ = 0.6***

**$^{1}$H NMR (300 MHz, $CDCl_3$):** δ 7.25 (dd, *J* = 8.9, 0.5 Hz, 1H), 6.85 (dd, *J* = 8.9, 2.6 Hz, 1H), 6.75 (d, *J* = 2.6 Hz, 1H), 3.95 (t, *J* = 6.4 Hz, 2H), 3.84 (s, 3H), 3.07 (t, *J* = 6.4 Hz, 2H), 2.26 (s, 1H). **$^{13}$C NMR (75 MHz, $CDCl_3$)**: δ 156.70, 156.51, 149.20, 131.46, 113.71, 111.74, 103.43, 64.27, 60.66, 56.04, 31.83. **DEPT-135 NMR (75 MHz, $CDCl_3$):** δ 113.69, 111.72, 103.41, 60.64, 56.03, 31.81. **IR ($CHCl_3$, $cm^{-1}$):** 3565, 3352, 2970, 2862, 1575, 1460, 1280, 1186, 1078, 1048, 1052, 960, 840, 769, 720, 650. **HRMS (ESI, + ve ion mode) (m/z):** $[M+Na]^+$ for $C_{11}H_{11}IO_3$: Calcd. 340.9651; Found 340.7354

**2-Iodo-3-(2-iodoethyl)-5-methoxybenzofuran (10) :**

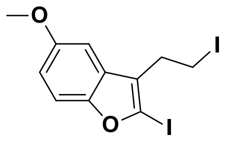

10

Starting from compound **9** (200 mg, 0.63 mmol), following the earlier depicted protocol, compound **10** was synthesized and the purification of which was achieved with 5% EtOAc in Petroleum Ether to afford the pure product **10** (228 mg, 85%) as pale yellow liquid. ***TLC→ (EtOAc:Petroleum Ether = 1:19 v/v); $R_f$ = 0.7***

**$^{1}$H NMR (300 MHz, $CDCl_3$):** δ 7.28 (dd, *J* = 8.9, 0.5 Hz, 1H), 6.90 (dd, *J* = 8.9, 2.6 Hz, 1H), 6.78 (d, *J* = 2.6 Hz, 1H), 3.86 (s, 3H), 3.42 (d, *J* = 1.5 Hz, 4H). **$^{13}$C NMR (75 MHz, $CDCl_3$):** δ 157.24, 156.73, 156.67, 149.17, 131.41, 115.16, 114.24, 111.94, 103.59, 64.45, 56.10, 56.07, 32.60, -0.04. **DEPT-135 NMR (75 MHz, $CDCl_3$):** δ 114.18, 111.87, 103.52, 56.03, 32.54, -0.10. **IR ($CHCl_3$, $cm^{-1}$):** 3032, 2948, 2864, 1572, 1496, 1470, 1442, 1280, 1176, 1083, 1021, 870, 791, 684, 520, 495. **HRMS (ESI, + ve ion mode) (m/z):** $[M+H]^+$ for $C_{11}H_{10}I_2O_2$: Calcd. 428.8848; Found 428.8849

**7-Ethyl-2-(2-(2-iodo-5-methoxybenzofuran-3-yl)ethyl)-2-azabicyclo[2.2.2]oct-5-ene (*Exo* + *Endo*) (11a & 11b):**

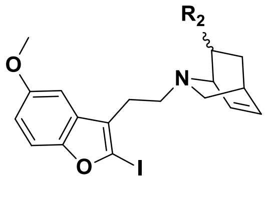

$R_2$ = - Et (*Exo*), **11a**
= - Et (*Endo*), **11b**

The coupling of compound **10** (220 mg, 0.51 mmol) with the -Cbz deprotected isoquinuclidine (200 mg) was performed as mentioned previously. The crude product was then purified on silica gel (230-400 mesh) eluting with gradually varying percentage of EtOAc (5% to 15%) in Petroleum Ether to afford the pure product containing both **11a** & **11b** (230 mg, 72%) as yellow oil. The characterization of the compounds was made by HRMS and was directly used in the next step.

**HRMS (ESI, + ve ion mode) (m/z):** $[M+H]^+$ for $C_{20}H_{24}INO_2$: Calcd. 438.093; Found 438.0792

**7-Ethyl-2-methoxy-6,6a,7,8,9,10,12,13-octahydro-6,9-methanobenzofuro[2,3-d]pyrido[1,2-a]azepine (*Exo* + *Endo*) (C1 & C2) :**

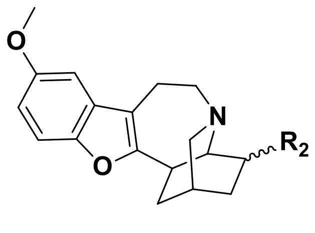

$R_2$ = - Et (*Exo*), **C1**
= - Et (*Endo*), **C2**

Utilizing the same protocol for **C3** & **C4**, starting from the combined mixture of **11a** & **11b** (150 mg), crude product was obtained which upon purification with gradually varying percentages of EtOAc (30% to 50%) in Petroleum Ether yielded **C1** (35 mg, 33%) and **C2** (51 mg, 48%) as deep yellow oil.

***Exo*-product (C1): $^{1}$H NMR (300 MHz, $CDCl_3$):** δ 7.45 – 7.32 (m, 2H), 7.09 (dd, *J* = 5.3, 2.6 Hz, 1H), 6.88 (ddd, *J* = 8.9, 2.6, 0.9 Hz, 1H), 3.91 – 3.82 (m, 3H), 3.19 – 2.91 (m, 4H), 2.43 (d, *J* = 10.5 Hz, 1H), 2.37 – 2.28 (m, 1H), 2.17 (d, *J* = 6.9 Hz, 1H), 2.04 – 1.97 (m, 1H), 1.93 (dd, *J* = 6.2, 3.4 Hz, 1H), 1.85 – 1.57 (m, 3H), 1.50 – 1.37 (m, 1H), 1.29 – 1.21 (m, 3H), 1.12 (d, *J* = 6.5 Hz, 2H), 0.88 (dt, *J* = 13.2, 7.3 Hz, 3H). **$^{13}$C NMR (75 MHz, $CDCl_3$):** δ 156.35, 156.18, 150.40, 142.97, 142.81, 128.81, 128.65, 128.37, 127.87, 117.07, 113.47, 112.18, 103.11, 102.17, 101.94, 77.68, 77.46, 77.26, 76.84, 66.42, 57.95, 56.31, 55.44, 55.00, 52.77, 52.32, 33.95, 32.25, 30.01, 29.90, 28.68, 28.21, 28.16, 25.46, 24.08, 21.26, 19.98, 17.61, 11.21. **DEPT-135 NMR (75 MHz, $CDCl_3$):** δ 142.92, 142.75, 137.96, 128.76, 125.98, 113.41, 112.12, 103.05, 102.10, 66.36, 57.89, 56.44, 56.31, 56.25, 56.13, 55.38, 54.94, 52.70, 52.26, 36.48, 33.89, 32.19, 29.94, 29.84, 29.42, 28.62, 28.15, 25.40, 24.02, 21.20, 19.92, 17.55, 11.15. **IR ($CHCl_3$, $cm^{-1}$):** 2954, 2924, 2853, 1698, 1613, 1474, 1224, 1028, 1175, 1101, 805. **HRMS (ESI, + ve ion mode) (m/z):** $[M+H]^+$ for $C_{20}H_{25}NO_2$: Calcd. 312.1964; Found 312.1940

***Endo*-product (C2): $^{1}$H NMR (300 MHz, $CDCl_3$):** δ 7.47 – 7.28 (m, 1H), 7.16 (dd, *J* = 9.9, 2.6 Hz, 1H), 6.93 – 6.84 (m, 1H), 3.90 – 3.83 (m, 3H), 3.65 – 3.48 (m, 1H), 3.44 – 3.32 (m, 1H), 3.25 – 3.01 (m, 4H), 2.48 – 2.30 (m, 1H), 2.09 – 1.91 (m, 2H), 1.75 – 1.56 (m, 2H), 1.54 – 1.38 (m, 1H), 1.34 – 1.18 (m, 2H), 1.13 (d, *J* = 6.5 Hz, 2H), 0.91 (dq, *J* = 21.2, 7.3 Hz, 3H). **$^{13}$C NMR (75 MHz, $CDCl_3$):** δ 156.05, 150.23, 142.82, 142.70, 137.96, 128.18, 127.78, 125.84, 116.77, 115.83,

113.66, 113.37, 112.09, 111.99, 102.85, 102.13, 102.02, 77.58, 77.36, 77.16, 76.74, 66.17, 57.58, 56.22, 56.22, 56.12, 55.74, 54.98, 54.84, 52.44, 52.10, 36.16, 33.49, 31.78, 29.82, 28.48, 27.98, 27.93, 27.67, 25.18, 23.81, 20.80, 19.74, 17.42, 11.86, 11.03. **DEPT-135 NMR (75 MHz, $CDCl_3$):** δ 142.92, 142.75, 137.96, 132.16, 128.76, 125.98, 113.78, 113.41, 112.24, 112.12, 103.05, 102.10, 101.87, 66.36, 57.89, 56.44, 56.31, 56.25, 56.13, 55.38, 54.94, 52.70, 52.26, 36.48, 33.89, 32.19, 29.94, 29.84, 29.42, 28.62, 28.15, 25.40, 24.02, 21.20, 17.55, 14.27, 11.15. **IR ($CHCl_3$, $cm^{-1}$):** 3061, 2926, 2860, 1663, 1613, 1474, 1436, 1260, 1178, 1118, 1027, 803, 695, 540. **HRMS (ESI, + ve ion mode) (m/z):** $[M+H]^+$ for $C_{20}H_{25}NO_2$: Calcd. 312.1964; Found 312.1937

❖ $^{1}H$ NMR (300 MHz, $CDCl_3$) of compound **1a** :

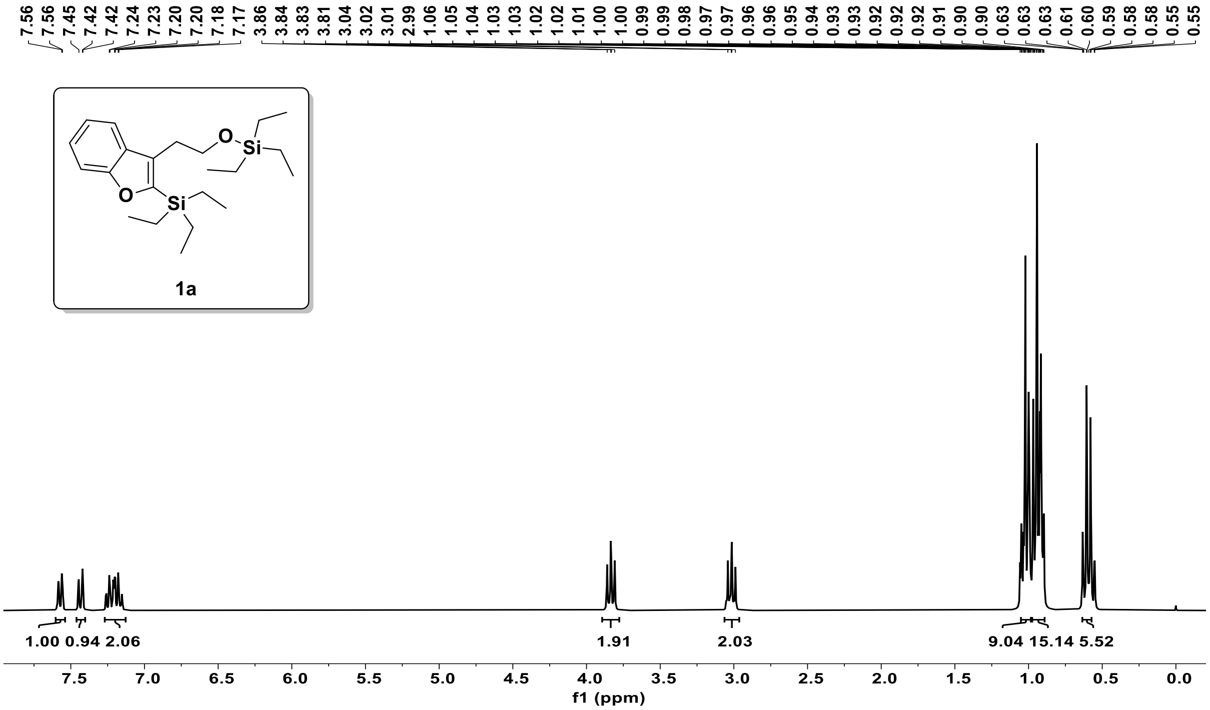


❖ $^{13}C$ NMR (75MHz, $CDCl_3$) of compound **1a** :

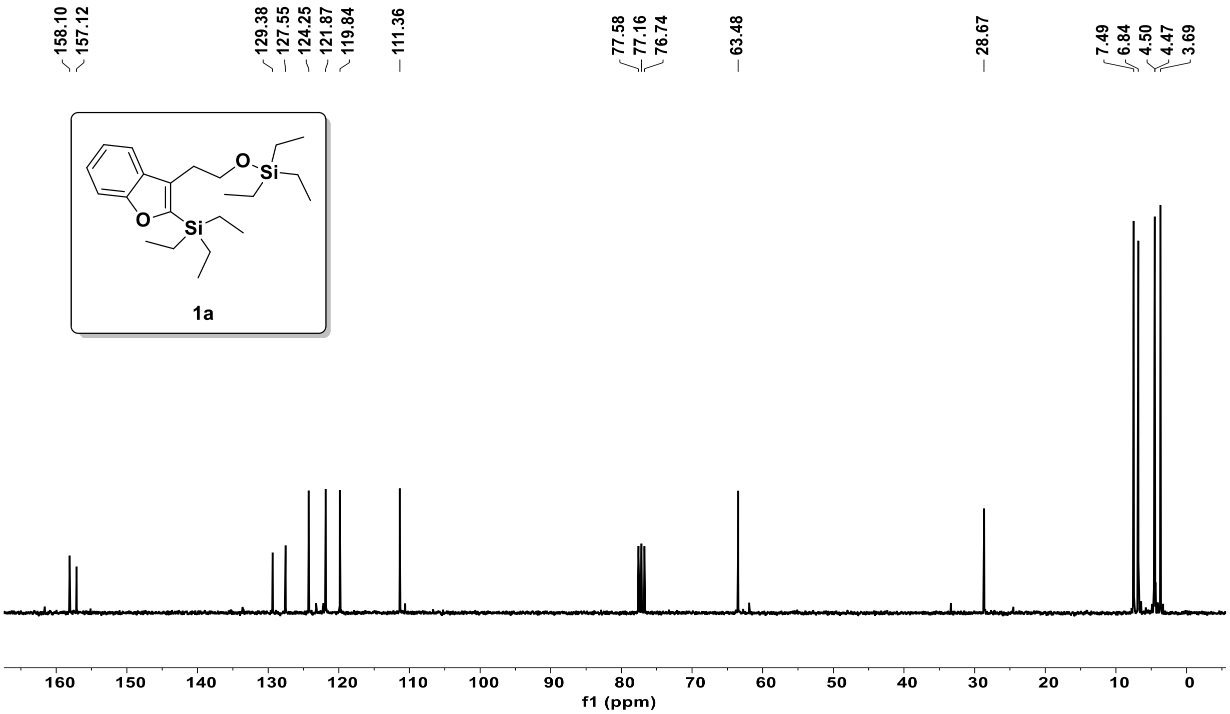

❖ DEPT-135NMR (75MHz, $CDCl_3$) of compound **1a**:

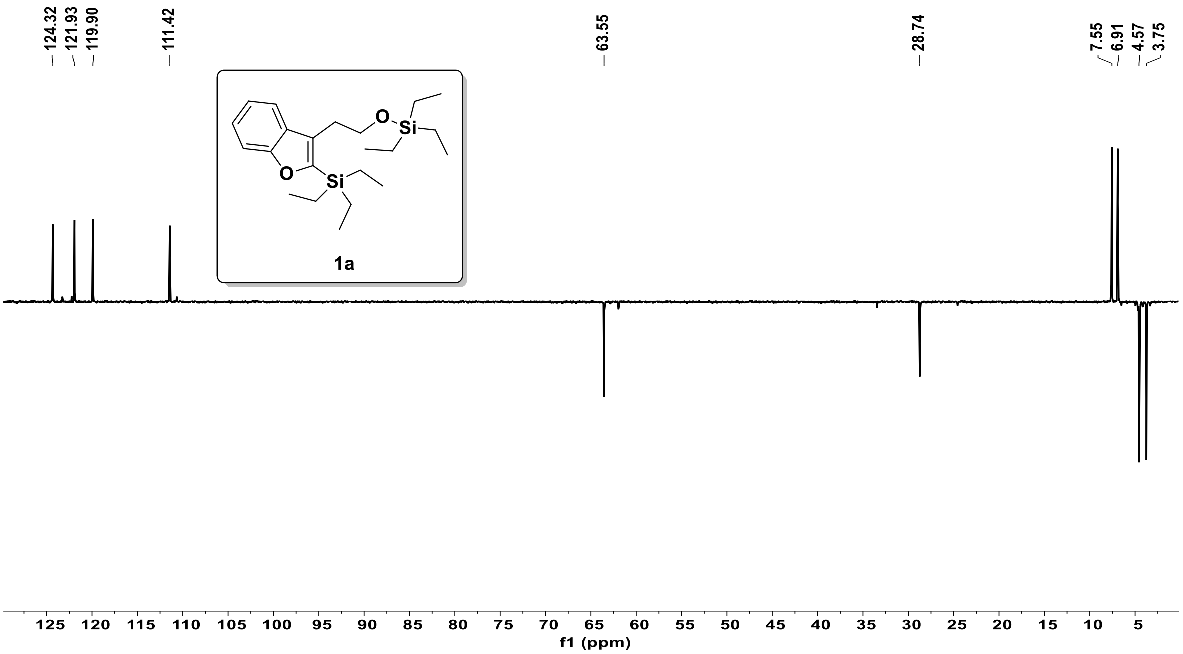


❖ $^1$H NMR (300 MHz, $CDCl_3$) of compound **2** :

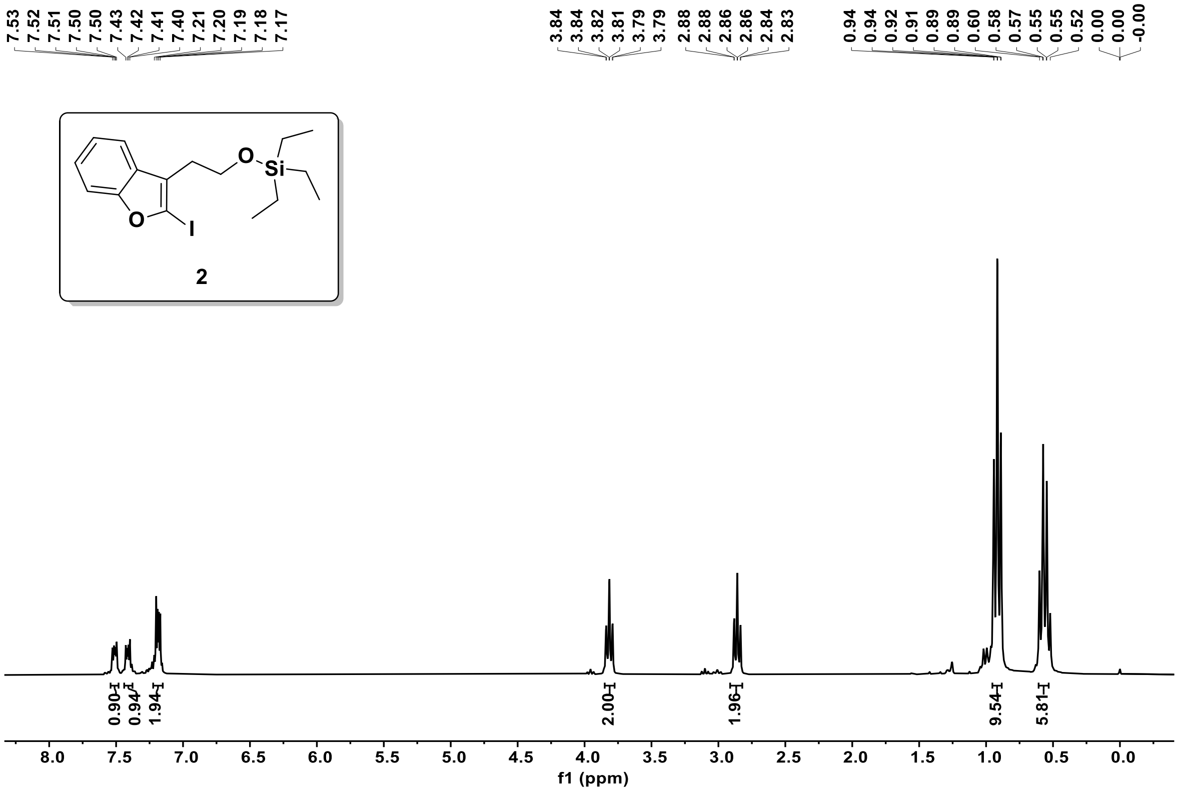

❖ $^{13}$C NMR (75MHz, $CDCl_3$) of compound **2** :

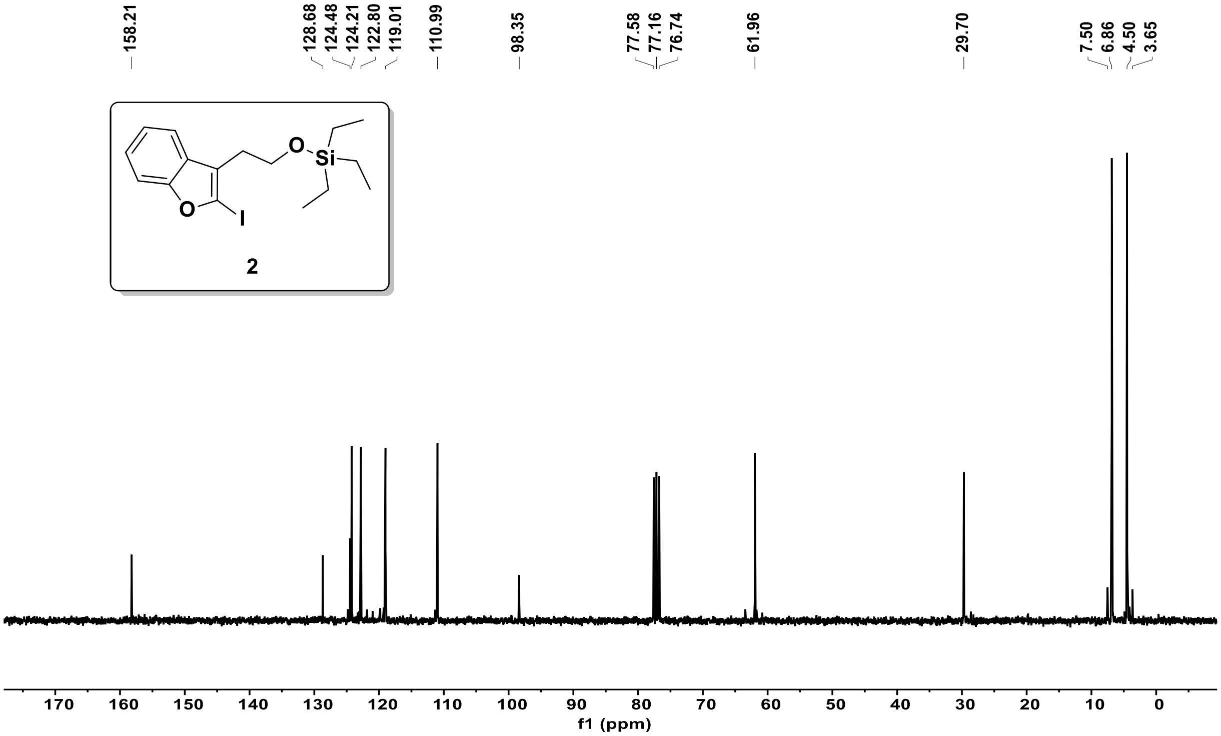


❖ DEPT-135 NMR (75MHz, $CDCl_3$) of compound **2**:

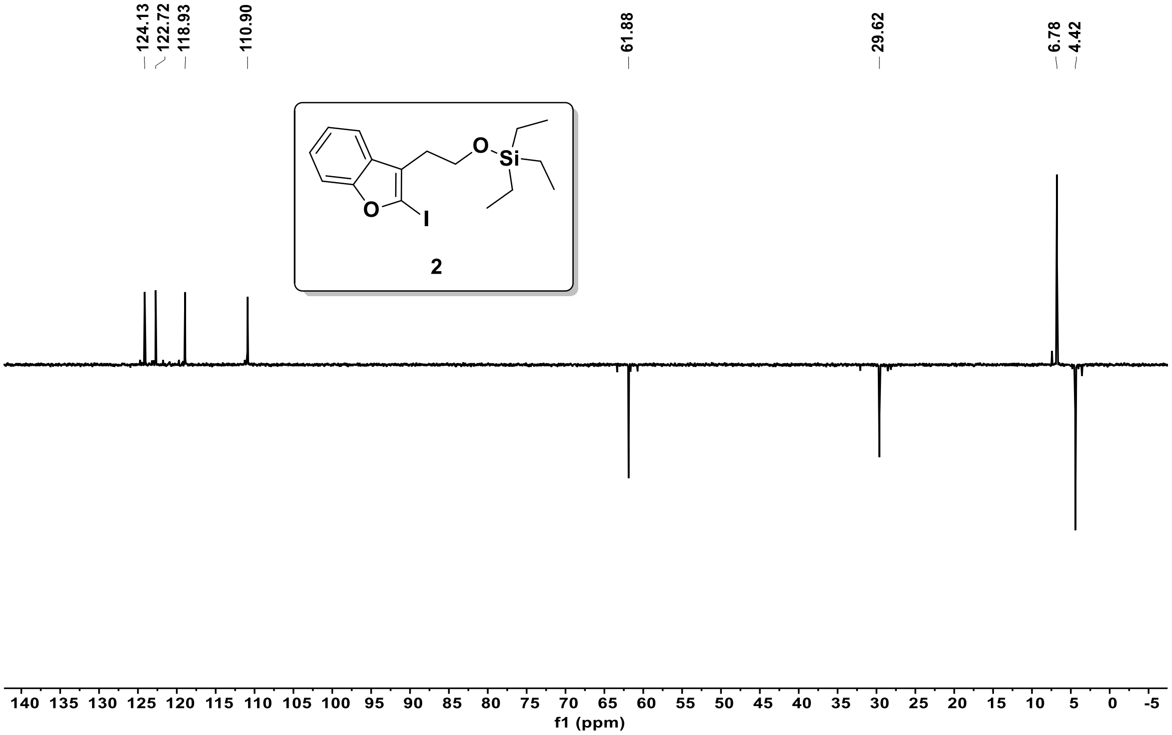

❖ $^{1}H$ NMR (300 MHz, $CDCl_3$) of compound **3**:

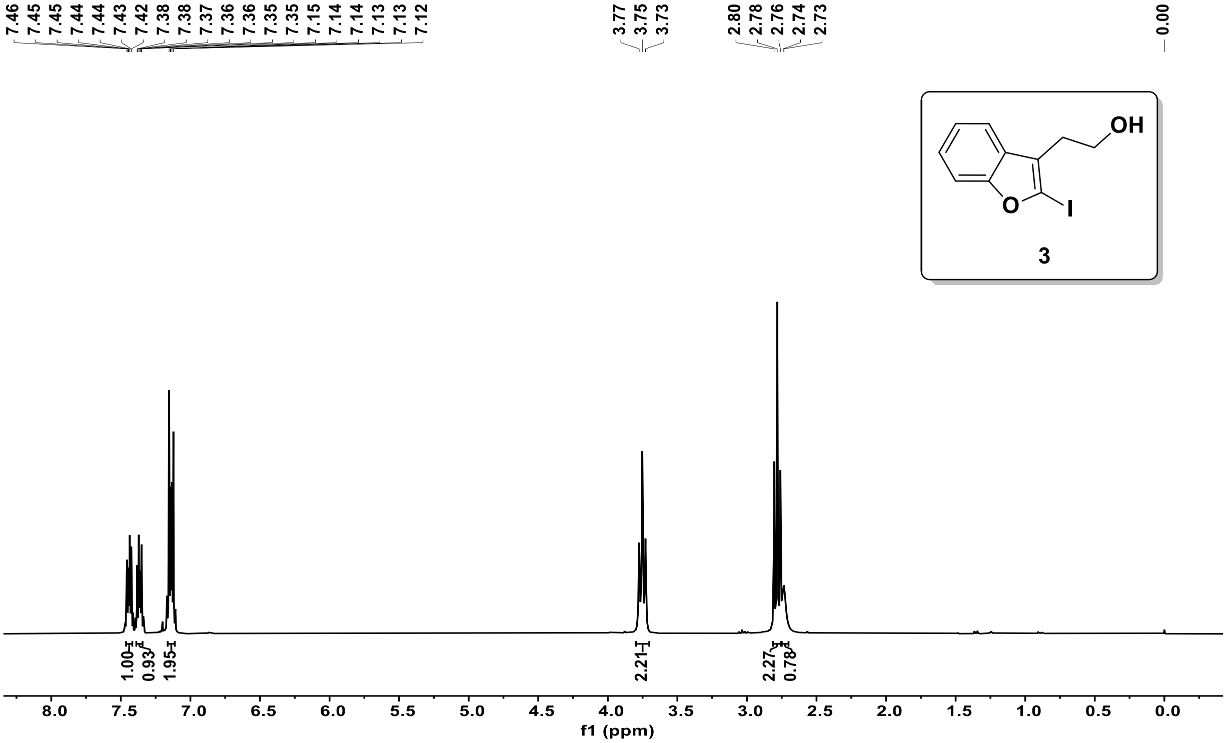


❖ $^{13}C$ NMR (75MHz, $CDCl_3$) of compound **3** :

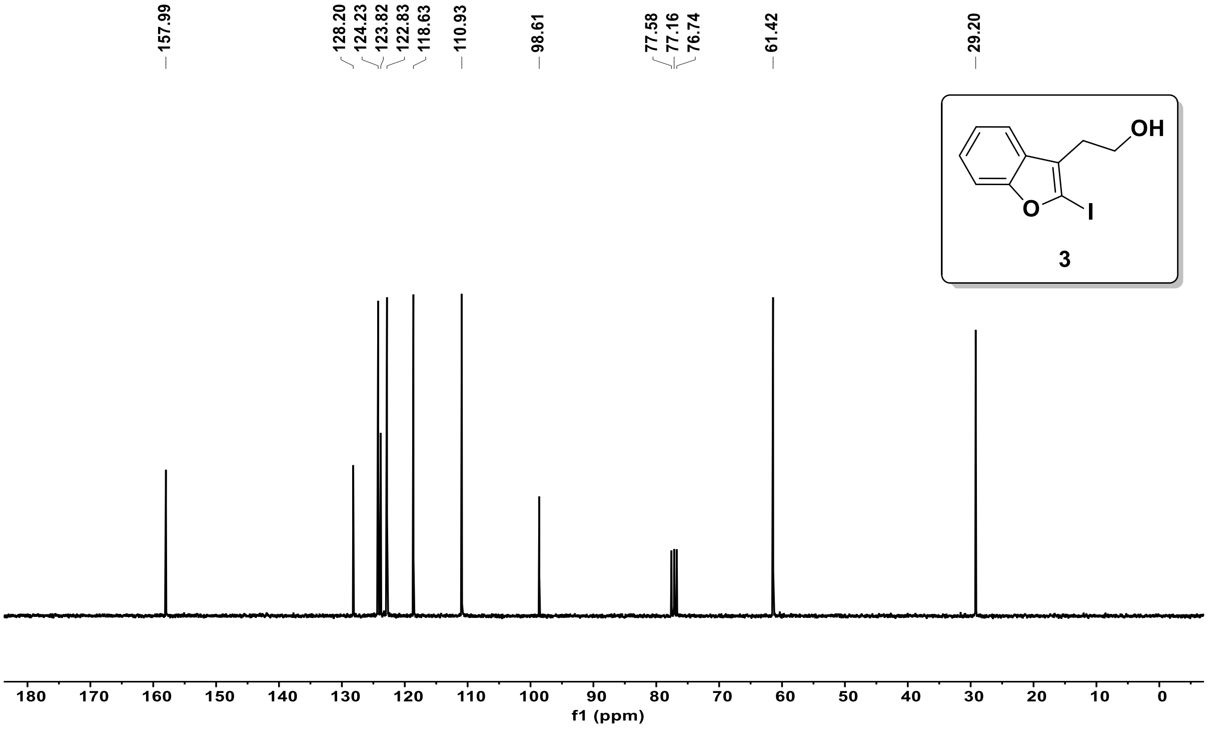

❖ DEPT-135 NMR (75MHz, $CDCl_3$) of compound **3** :

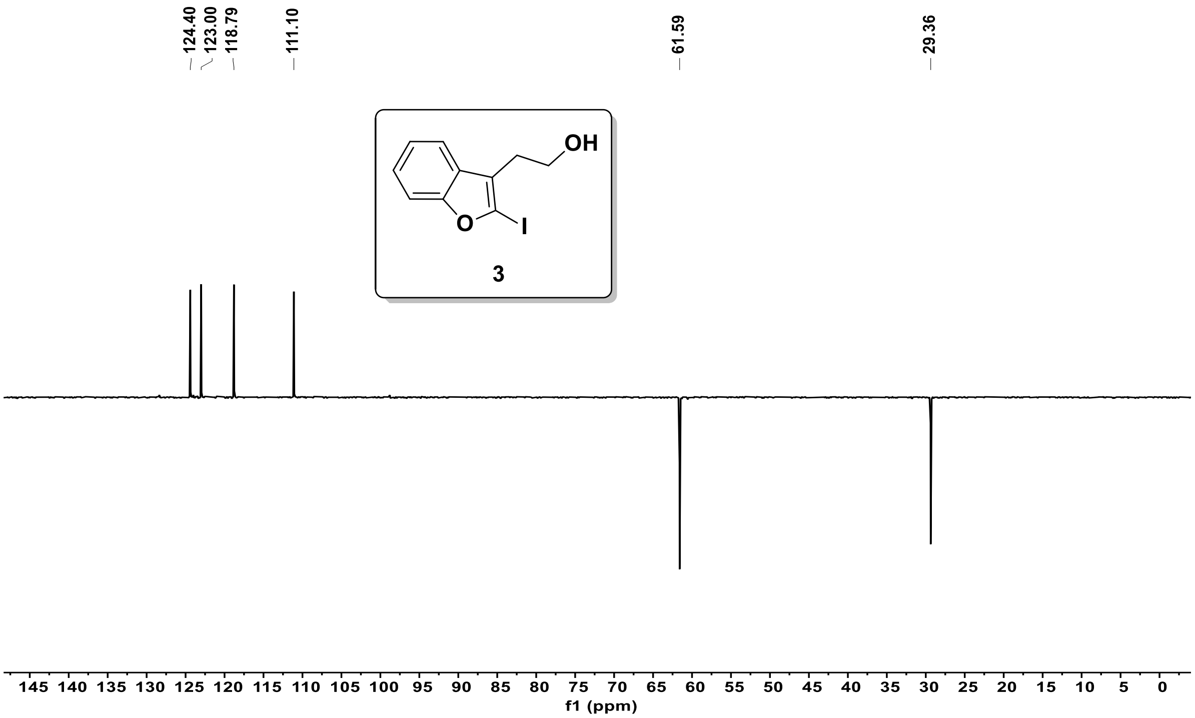


❖ $^1$H NMR (300 MHz, $CDCl_3$) of compound **4** :

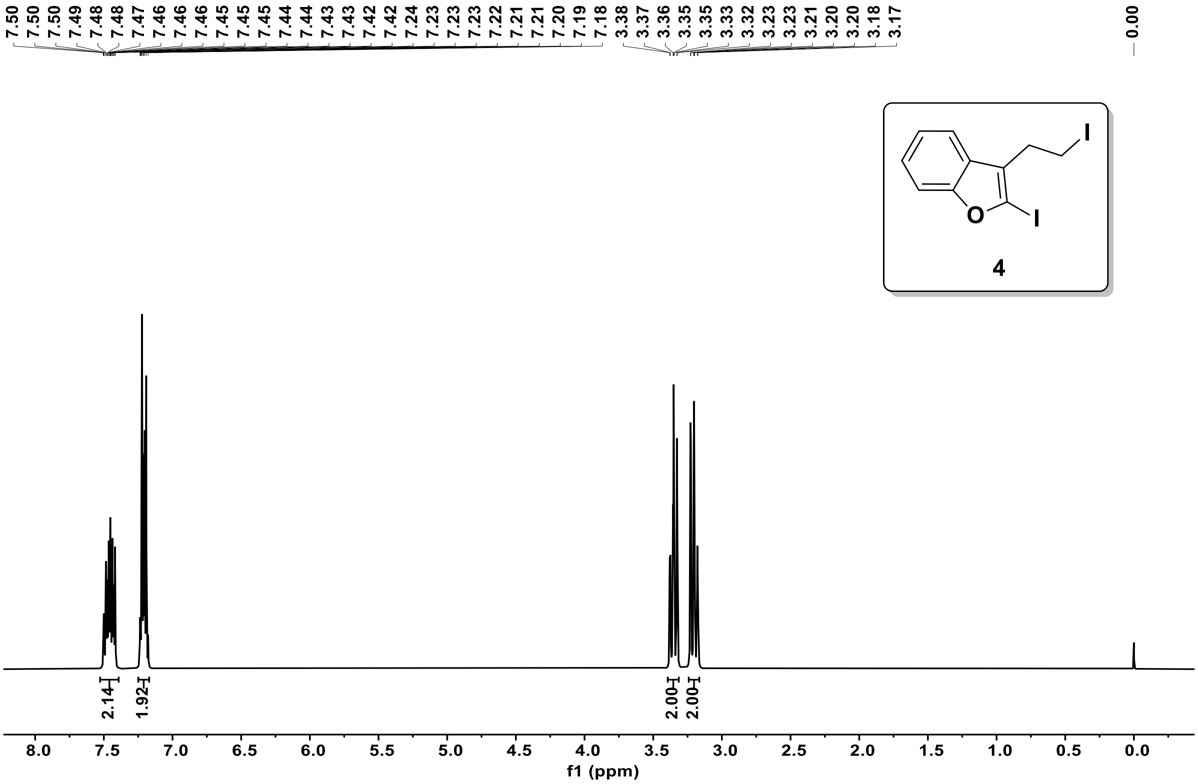

❖ $^{13}$C NMR (75MHz, $CDCl_3$) of compound **4** :

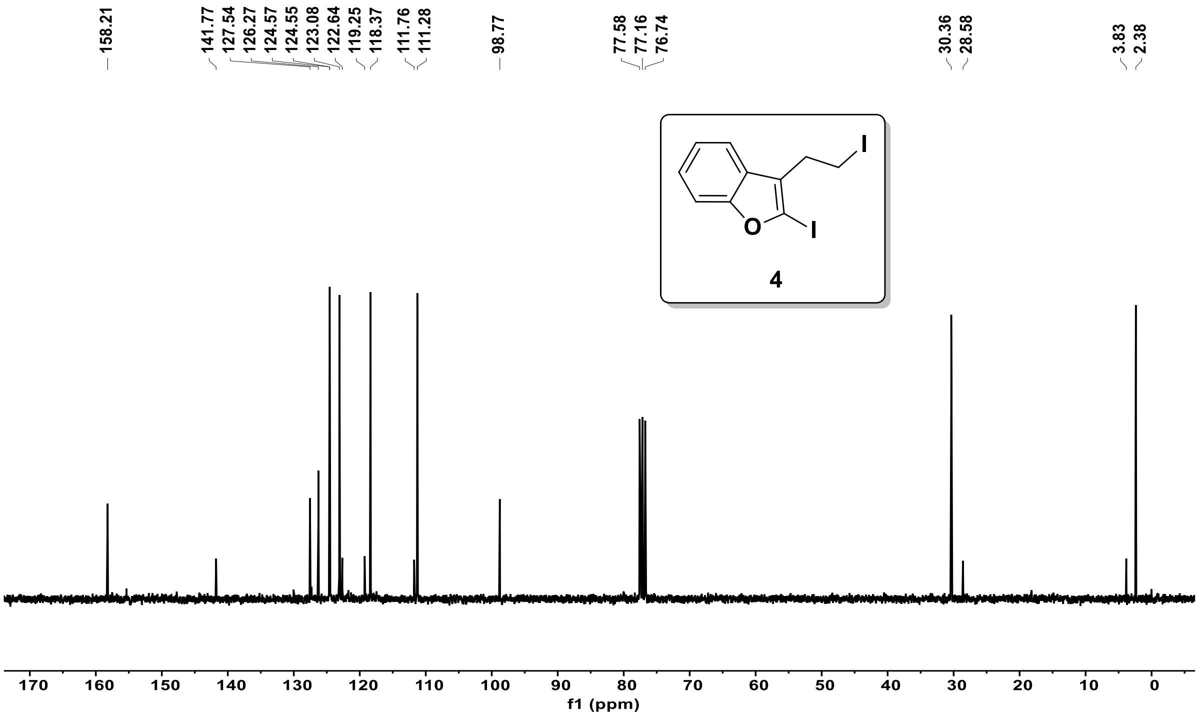


❖ DEPT-135 NMR (75MHz, $CDCl_3$) of compound **4** :

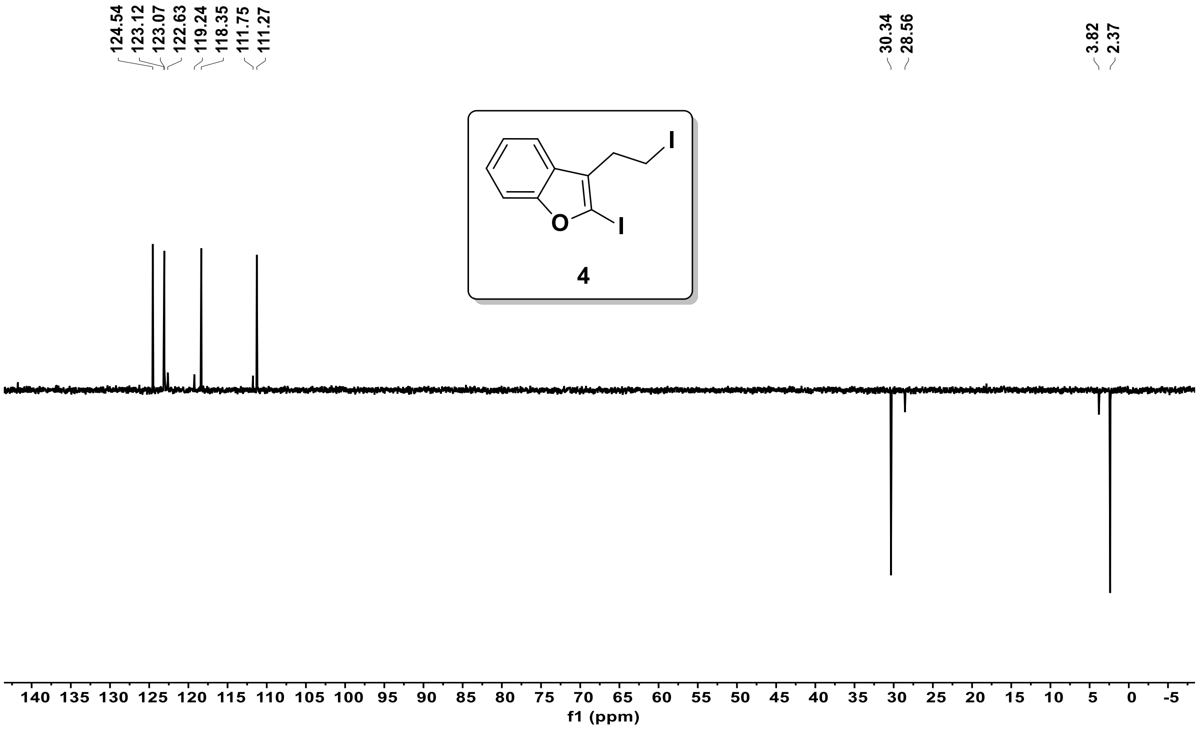

❖ $^{1}$H NMR (300 MHz, $CDCl_3$) of compound **5 (*Exo* + *Endo*)** :

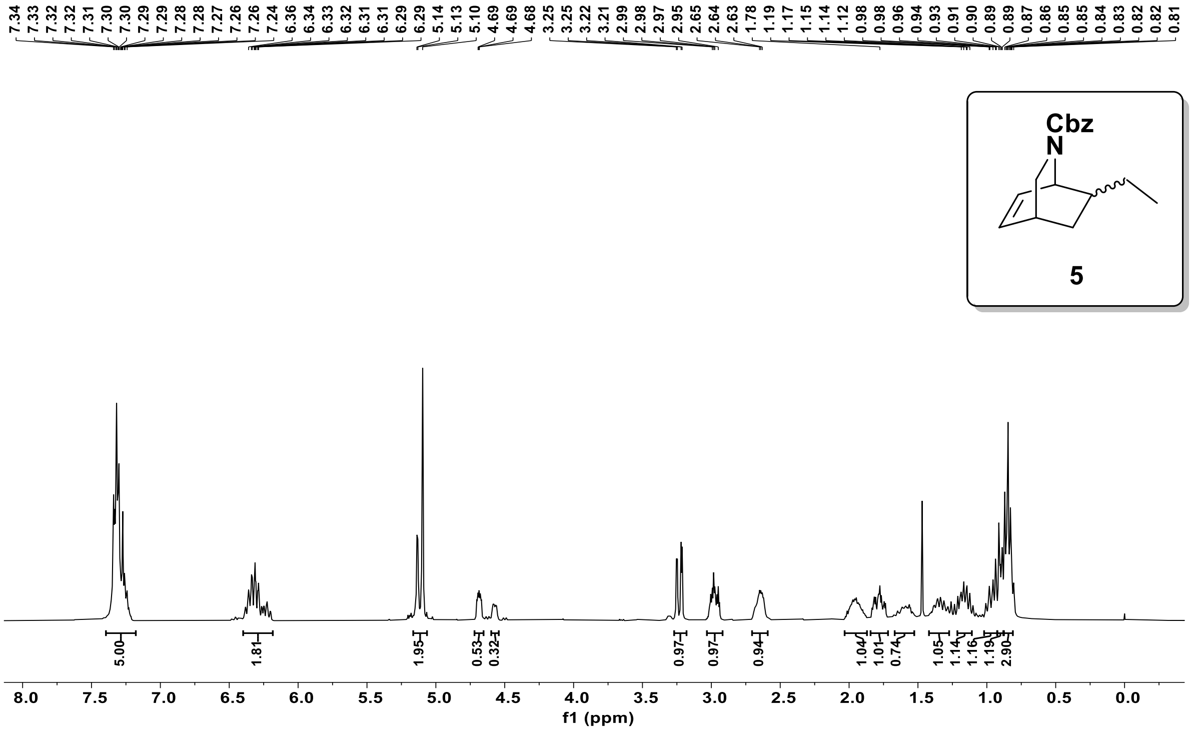


❖ $^{13}$C NMR (75MHz, $CDCl_3$) of compound **5 (*Exo* + *Endo*)** :

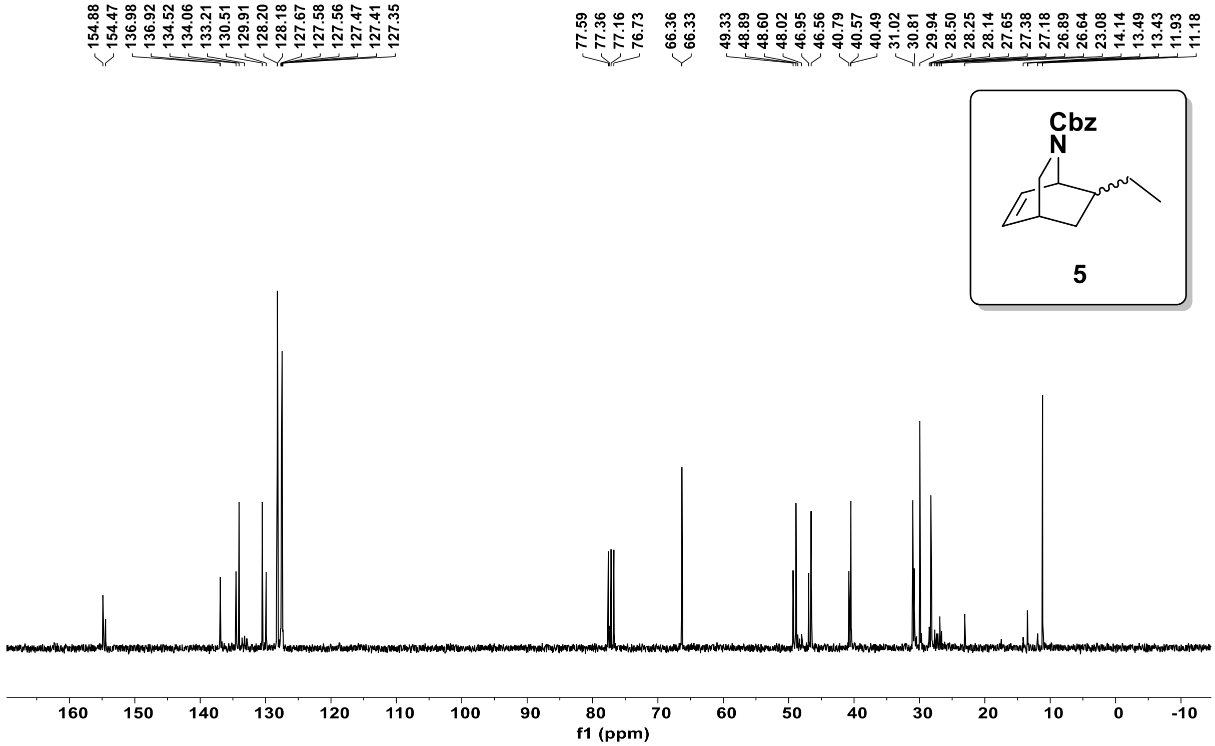

❖ DEPT-135 NMR (75MHz, $CDCl_3$) of compound **5 (*Exo* + *Endo*)** :

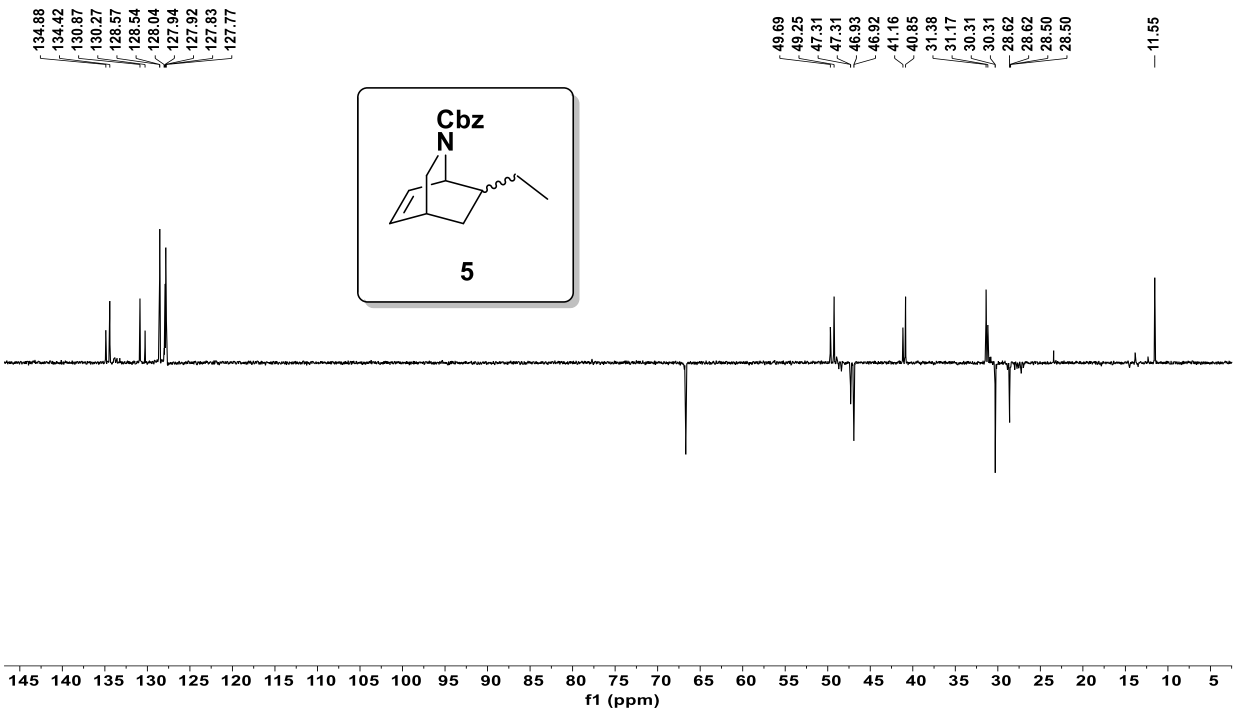


❖ $^1$H NMR (300 MHz, $CDCl_3$) of compound **C3** :

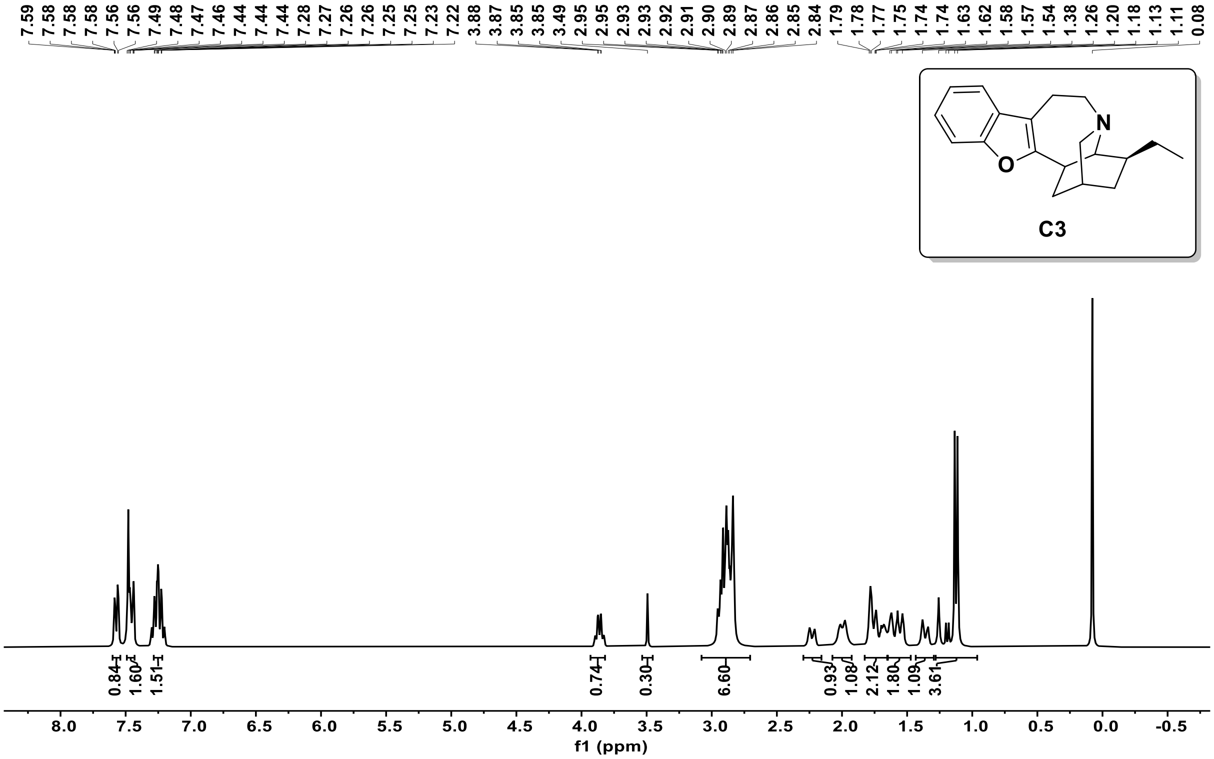

❖ $^{13}$C NMR (75MHz, $CDCl_3$) of compound **C3** :

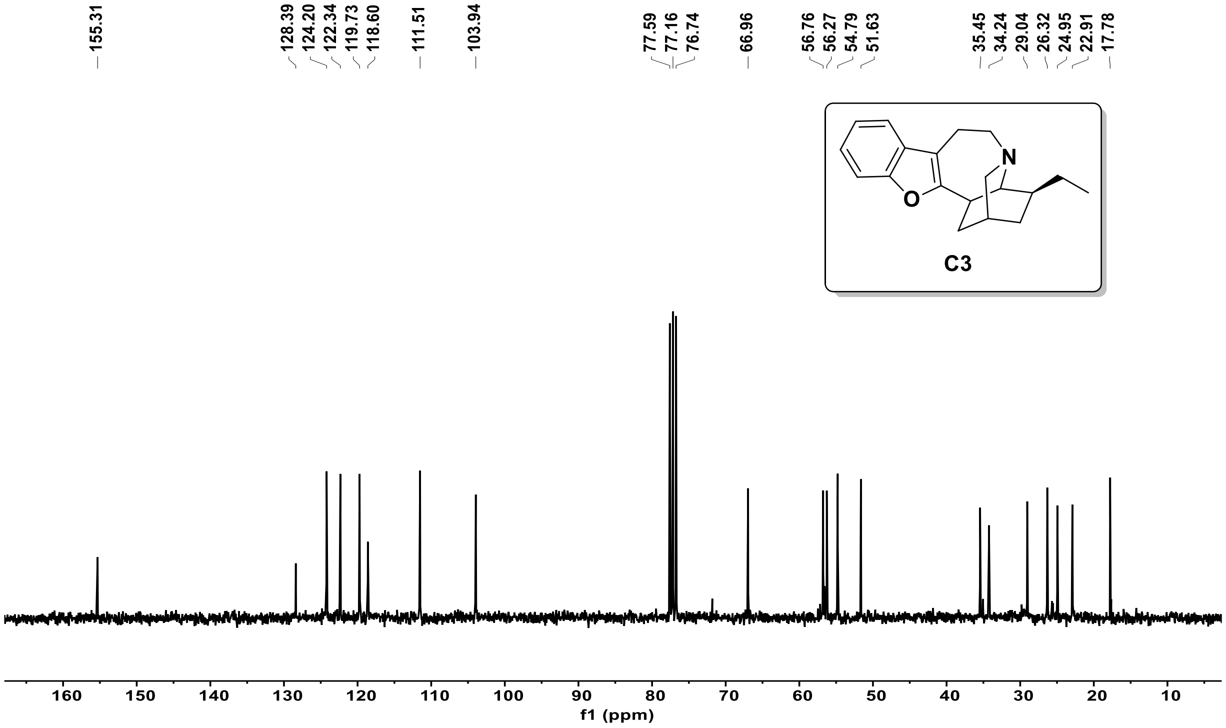


❖ DEPT-135 NMR (75MHz, $CDCl_3$) of compound **C3** :

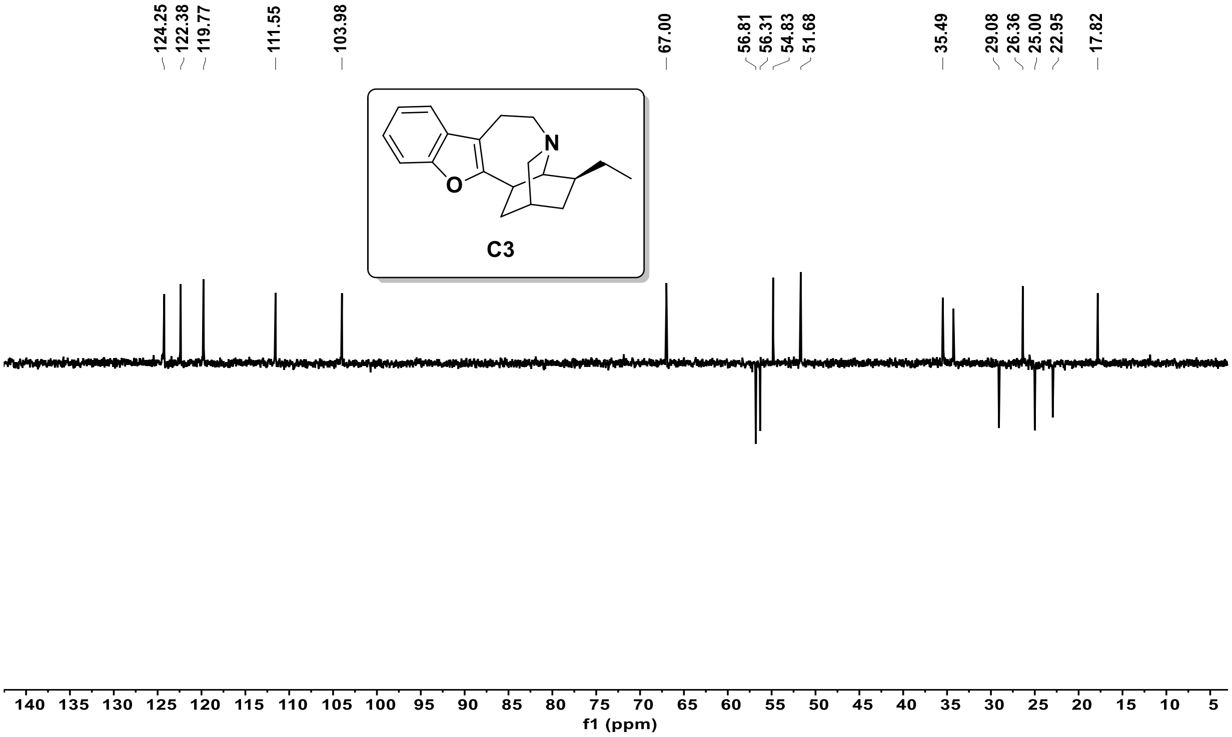

❖ $^{1}$H NMR (300 MHz, $CDCl_3$) of compound **C4** :

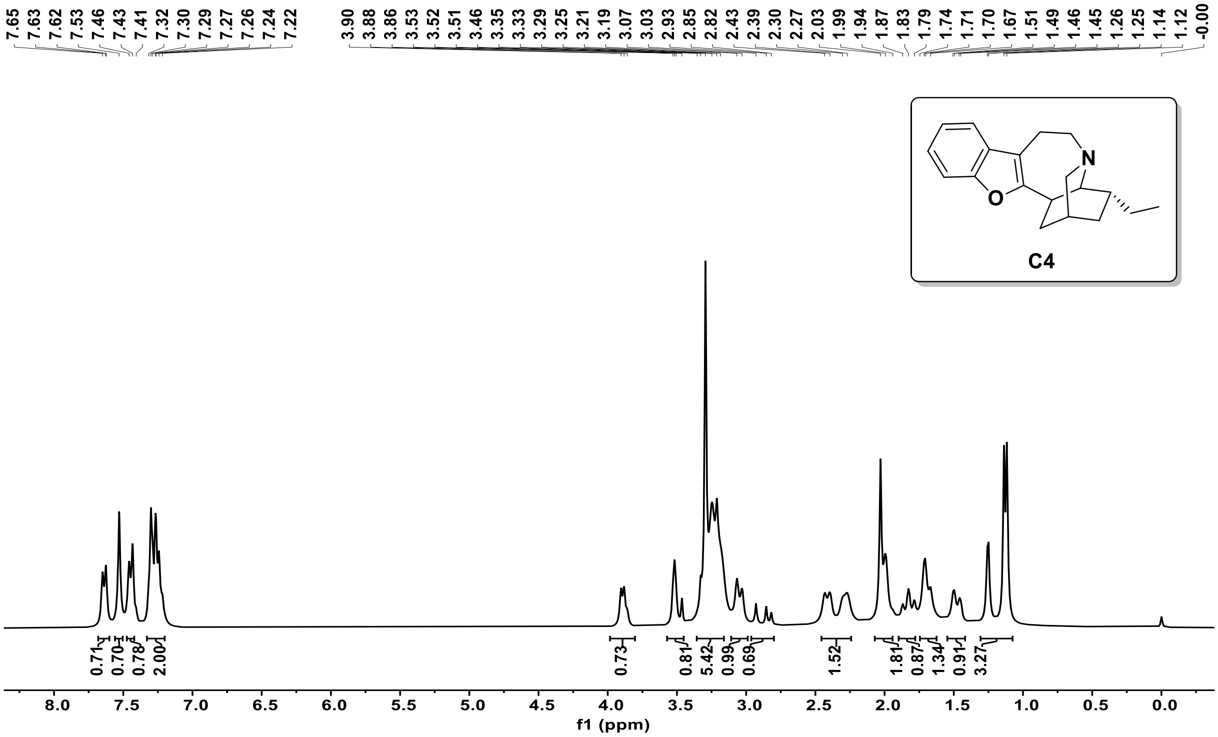


❖ $^{13}$C NMR (75MHz, $CDCl_3$) of compound **C4** :

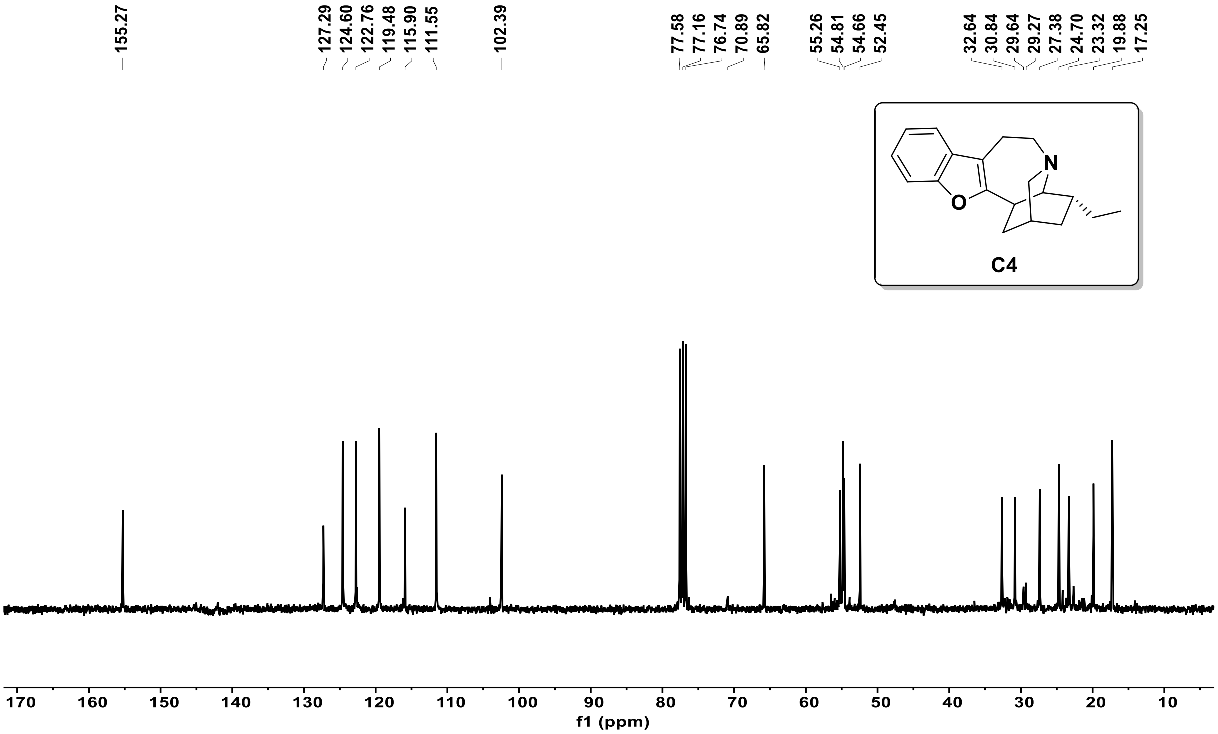

❖ DEPT-135 NMR (75MHz, $CDCl_3$) of compound **C4** :

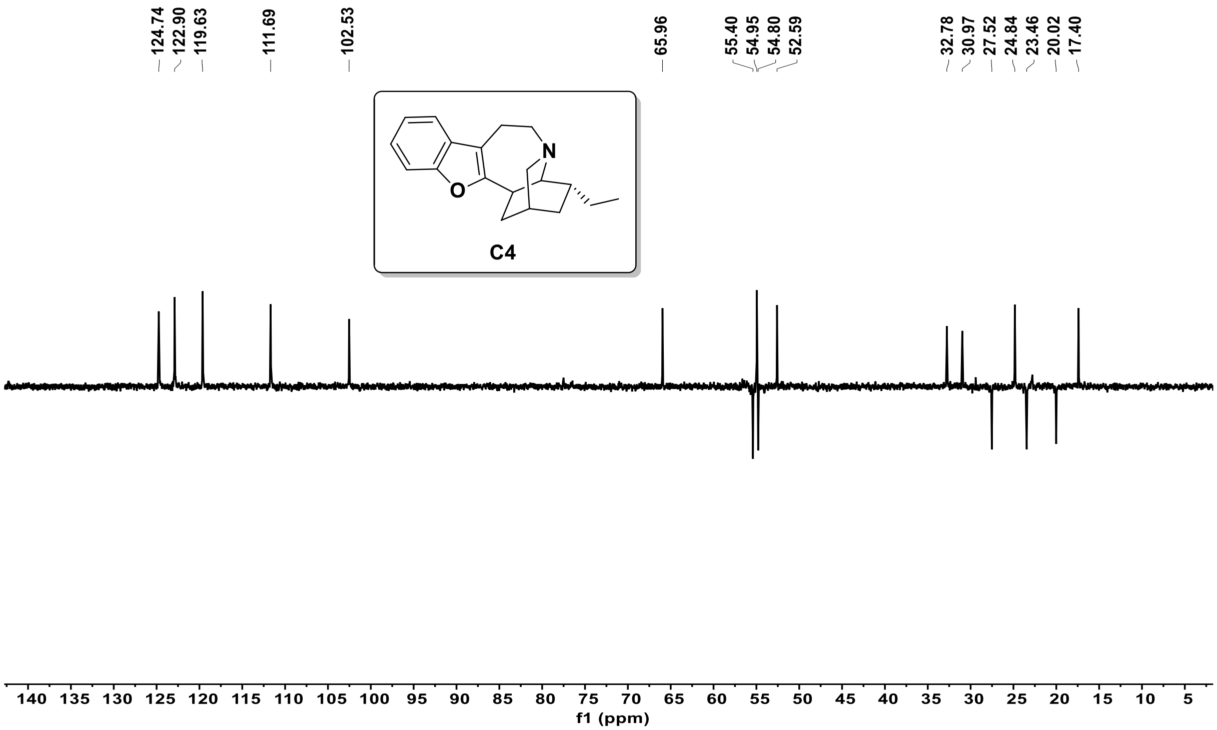


❖ [1]H NMR (300 MHz, $CDCl_3$) of compound **8** :

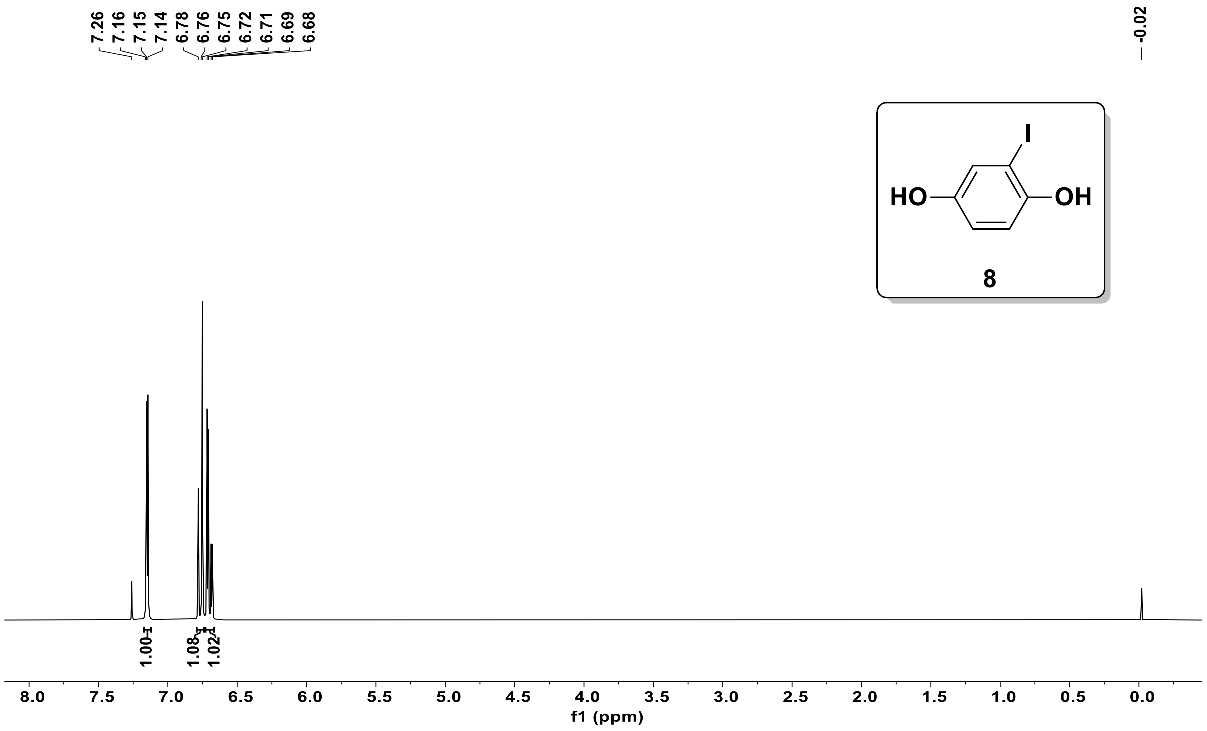

❖ $^{13}$C NMR (75MHz, $CDCl_3$) of compound **8** :

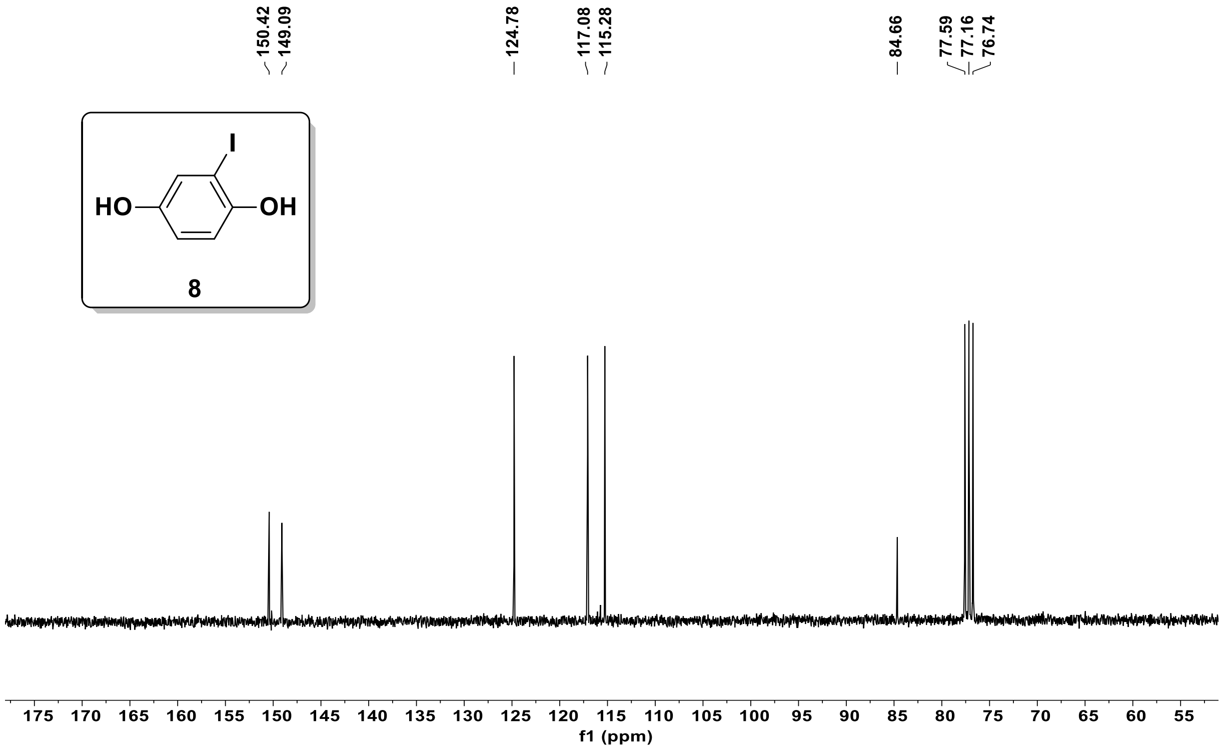


❖ DEPT-135 NMR (75MHz, $CDCl_3$) of compound **8** :

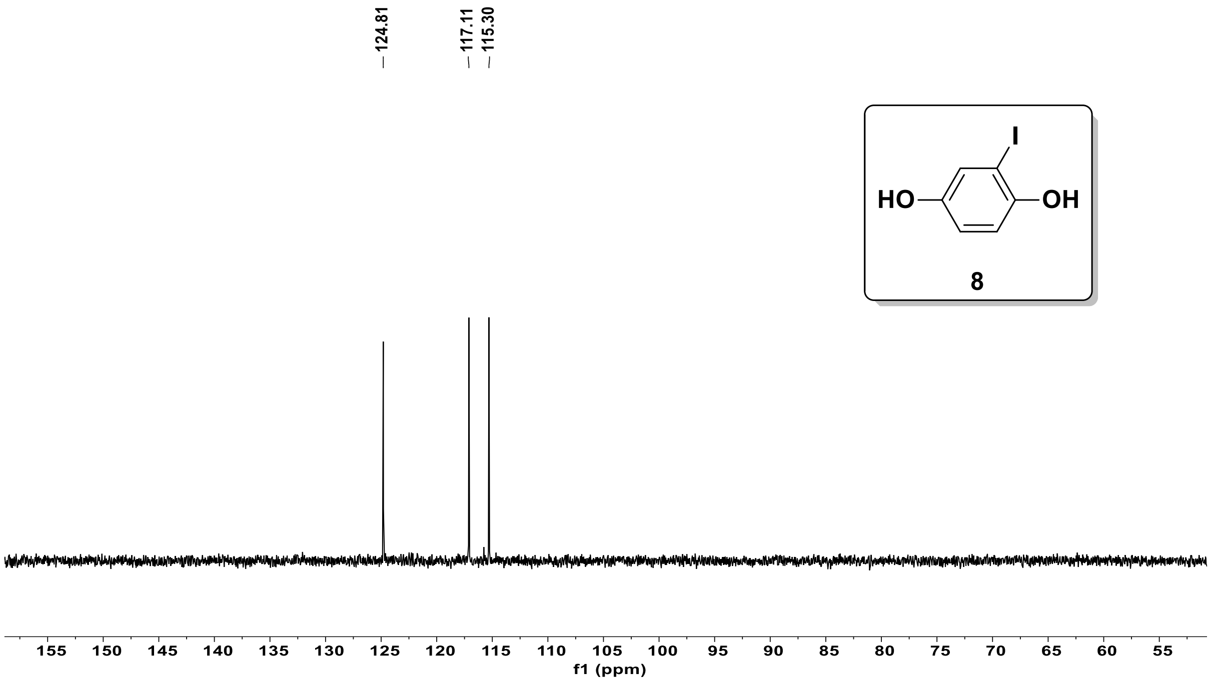

❖ $^{1}$H NMR (300 MHz, $CDCl_3$) of compound **9** :

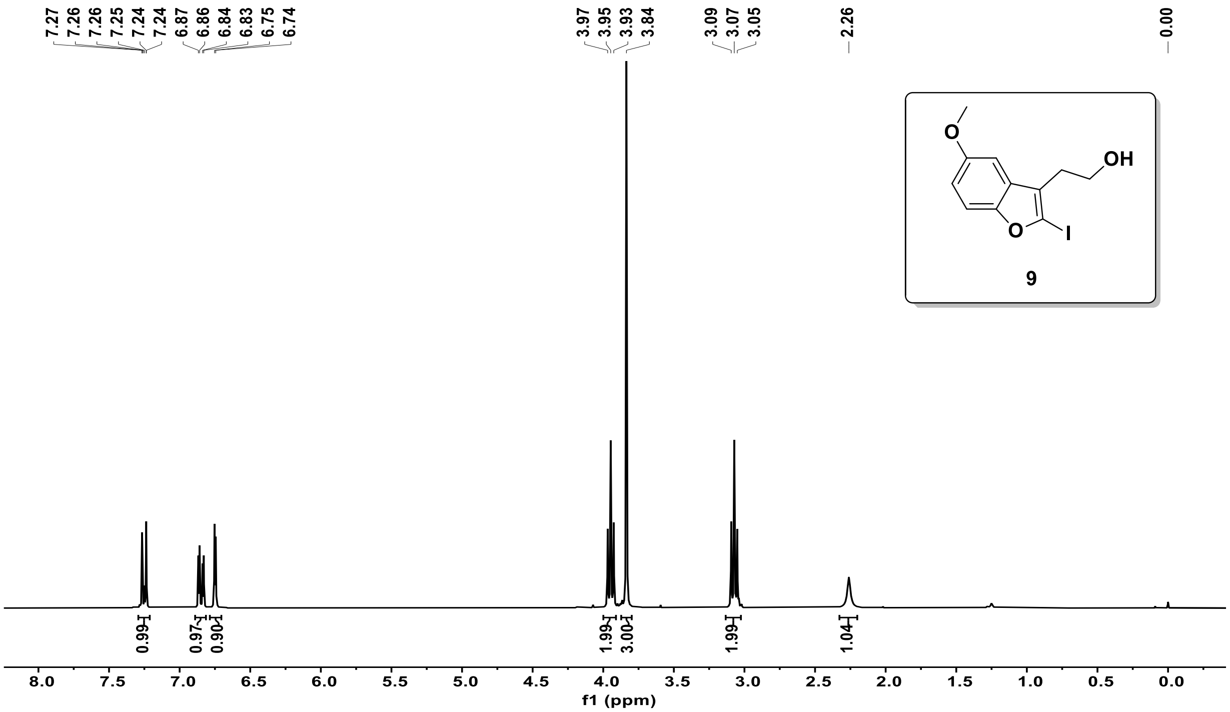


❖ $^{13}$C NMR (75MHz, $CDCl_3$) of compound **9** :

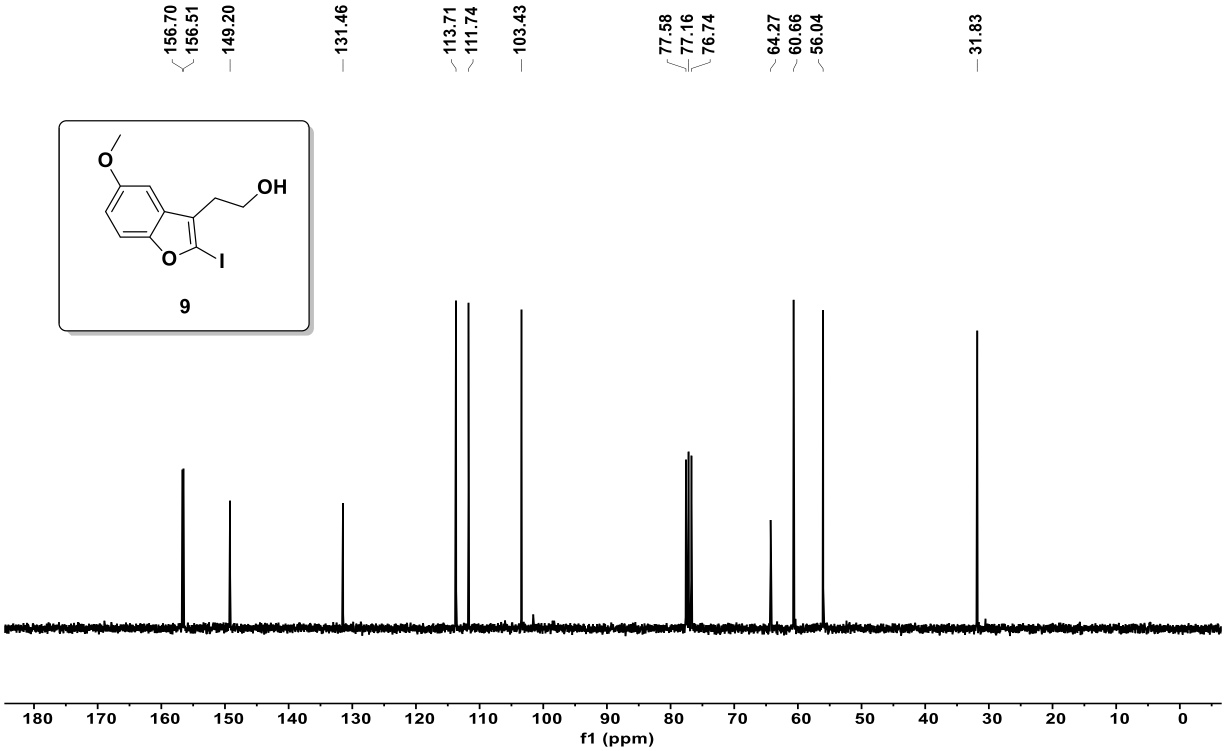

❖ DEPT-135 NMR (75MHz, $CDCl_3$) of compound **9** :

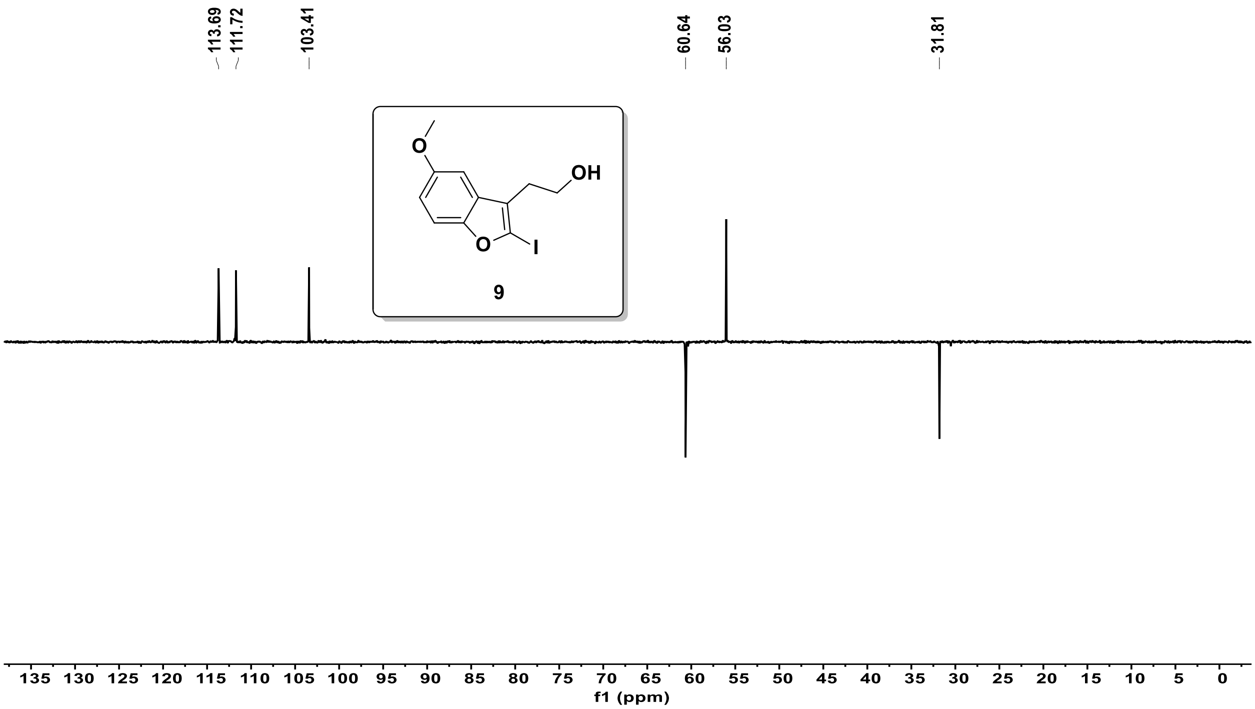


❖ [1]H NMR (300 MHz, $CDCl_3$) of compound **10** :

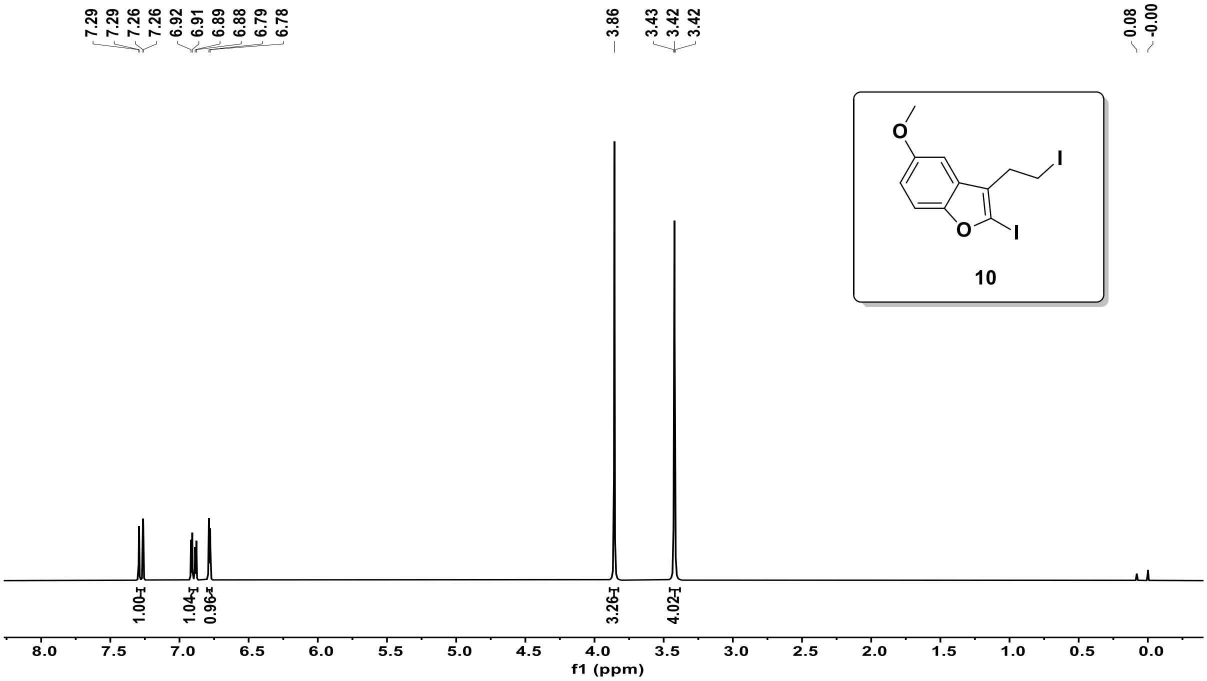

❖ $^{13}$C NMR (75MHz, $CDCl_3$) of compound **10** :

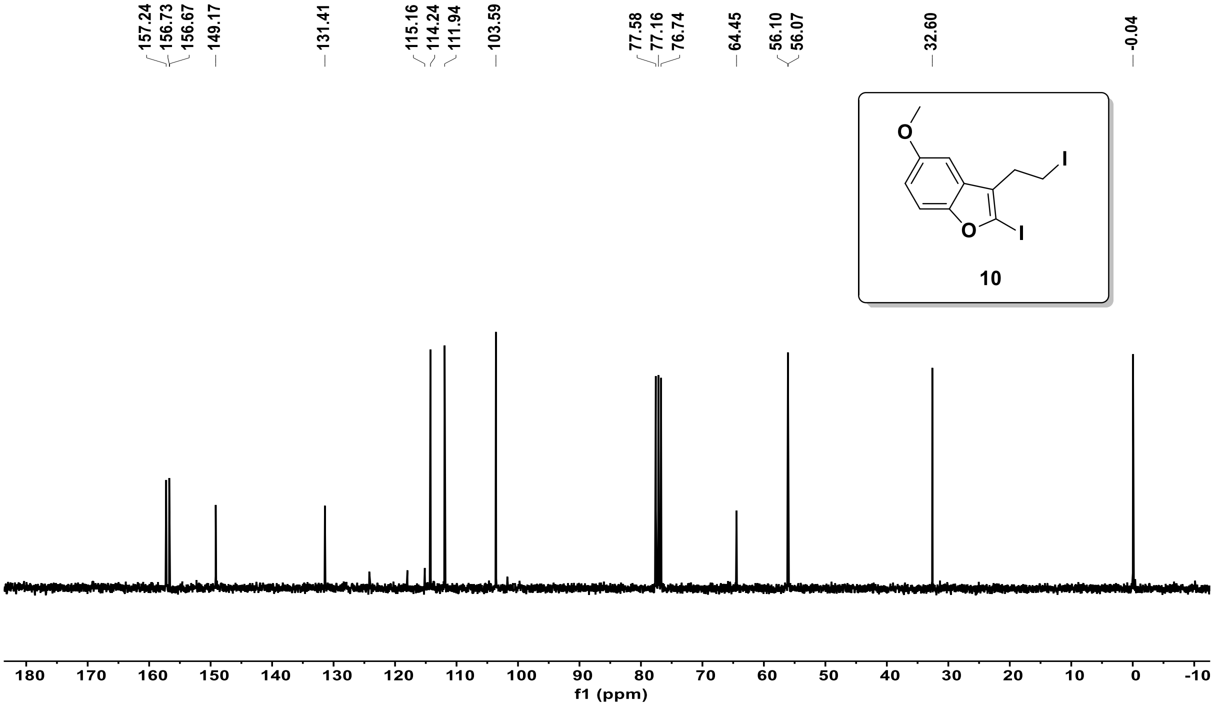


❖ DEPT-135 NMR (75MHz, $CDCl_3$) of compound **10** :

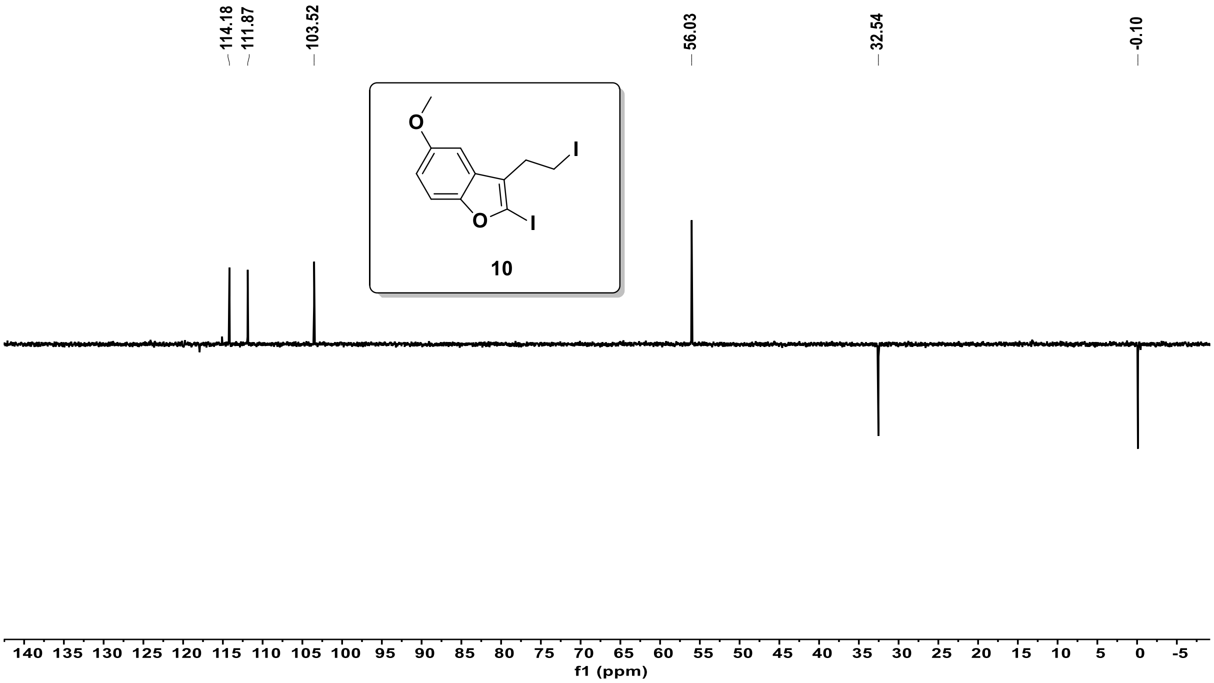

❖ $^{1}$H NMR (300 MHz, $CDCl_3$) of compound **C1** :

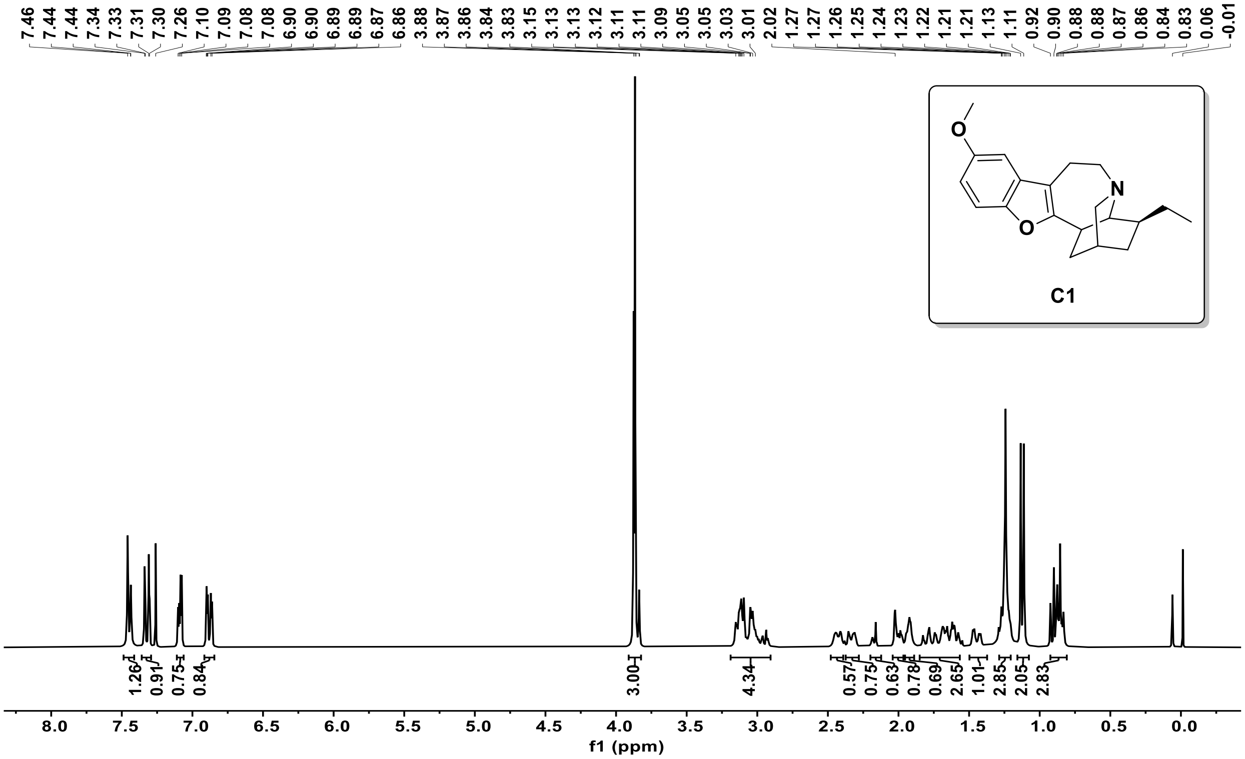


❖ $^{13}$C NMR (75MHz, $CDCl_3$) of compound **C1** :

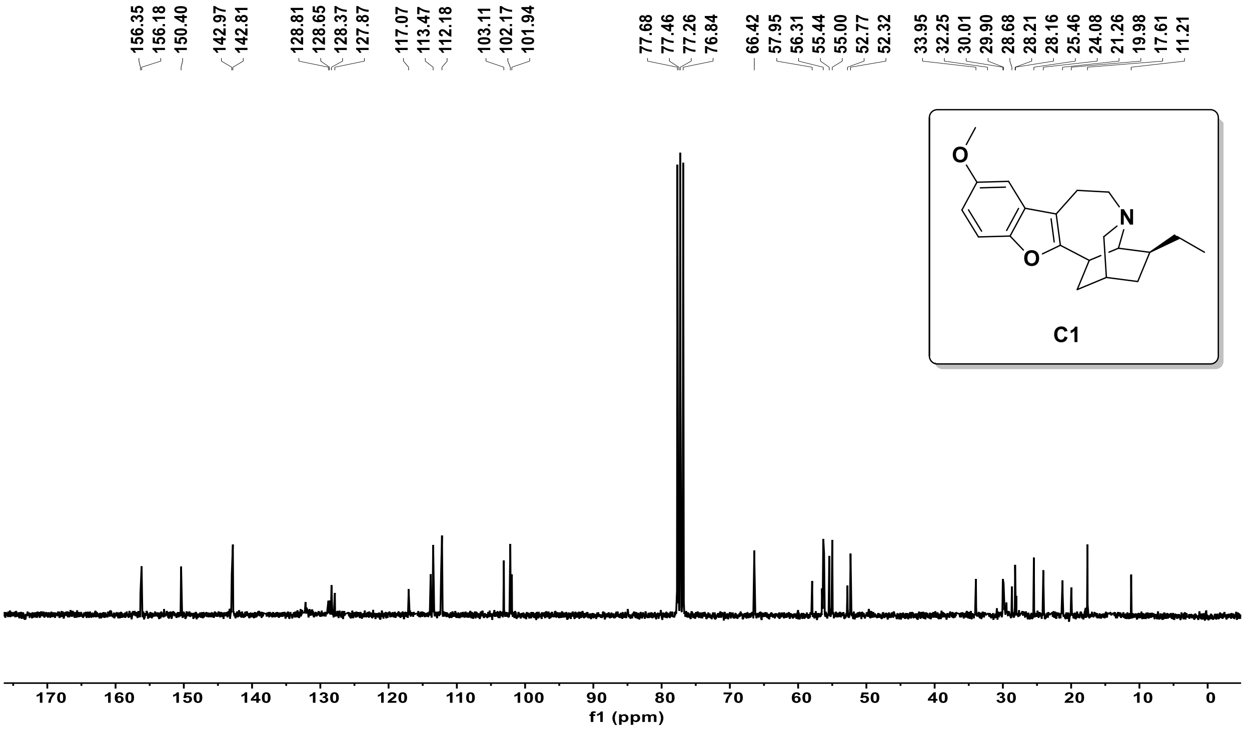

❖ DEPT-135 NMR (75MHz, $CDCl_3$) of compound **C1** :

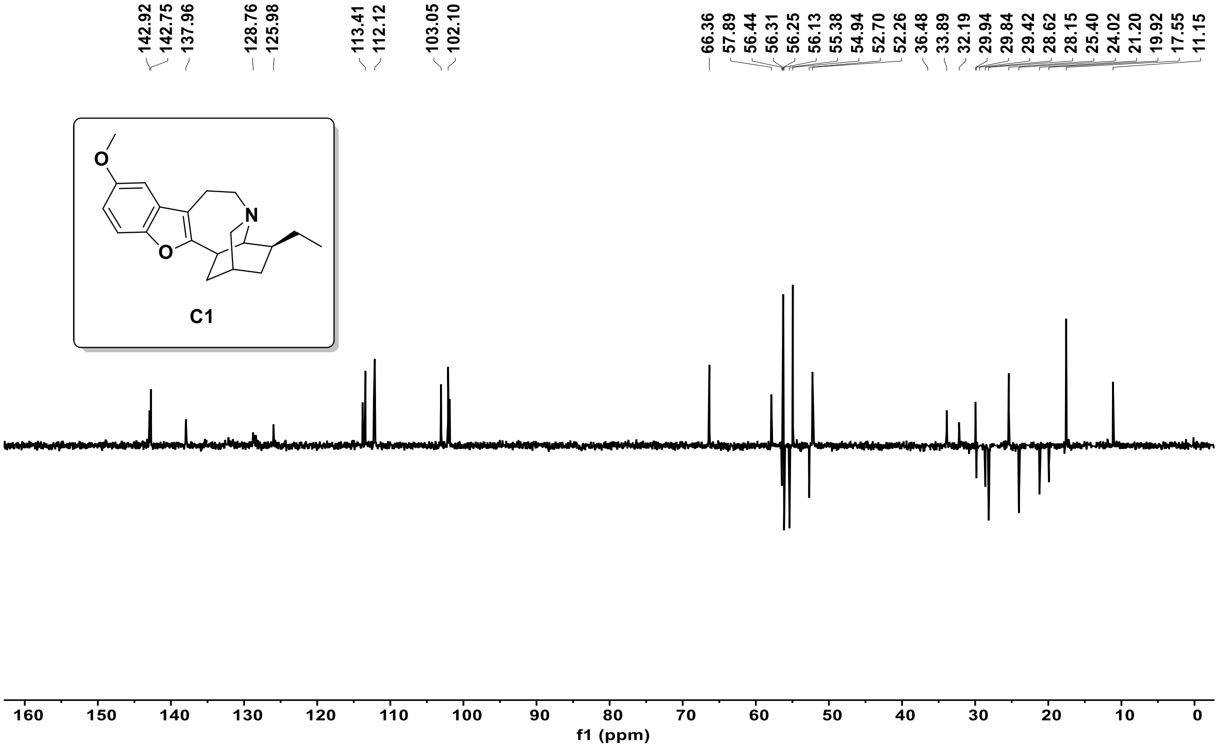


❖ [1]H NMR (300 MHz, $CDCl_3$) of compound **C2** :

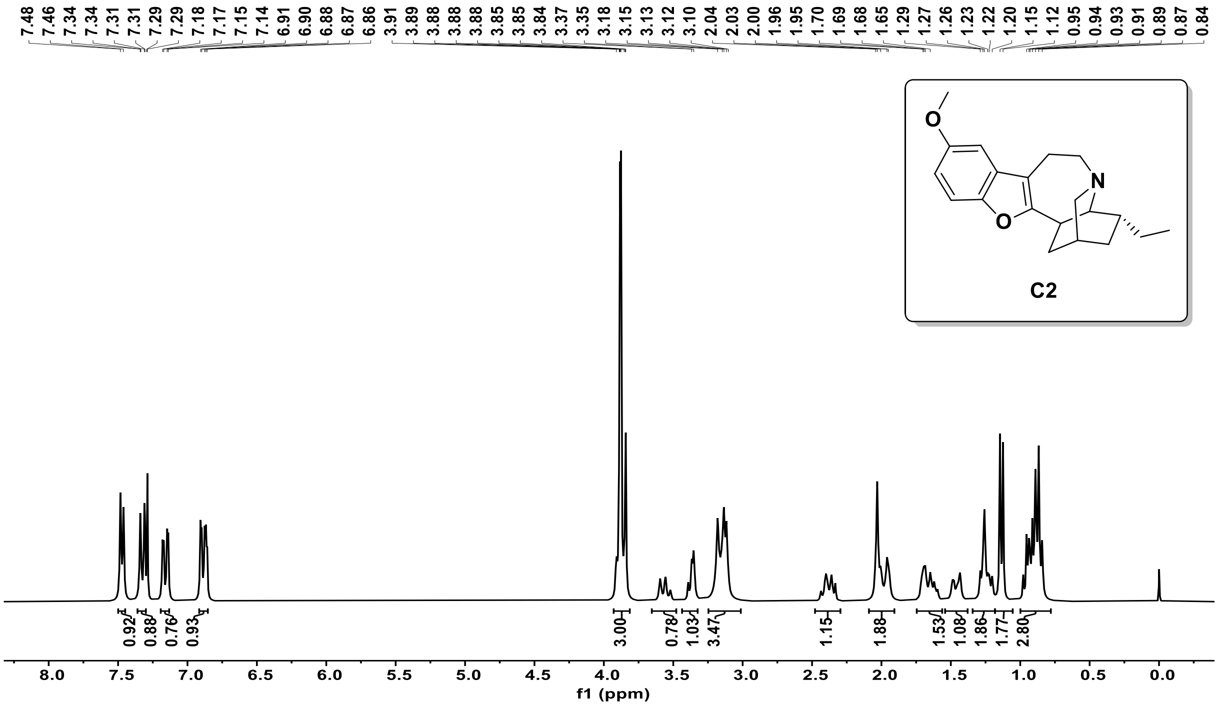

❖ $^{13}$C NMR (75MHz, $CDCl_3$) of compound **C2** :

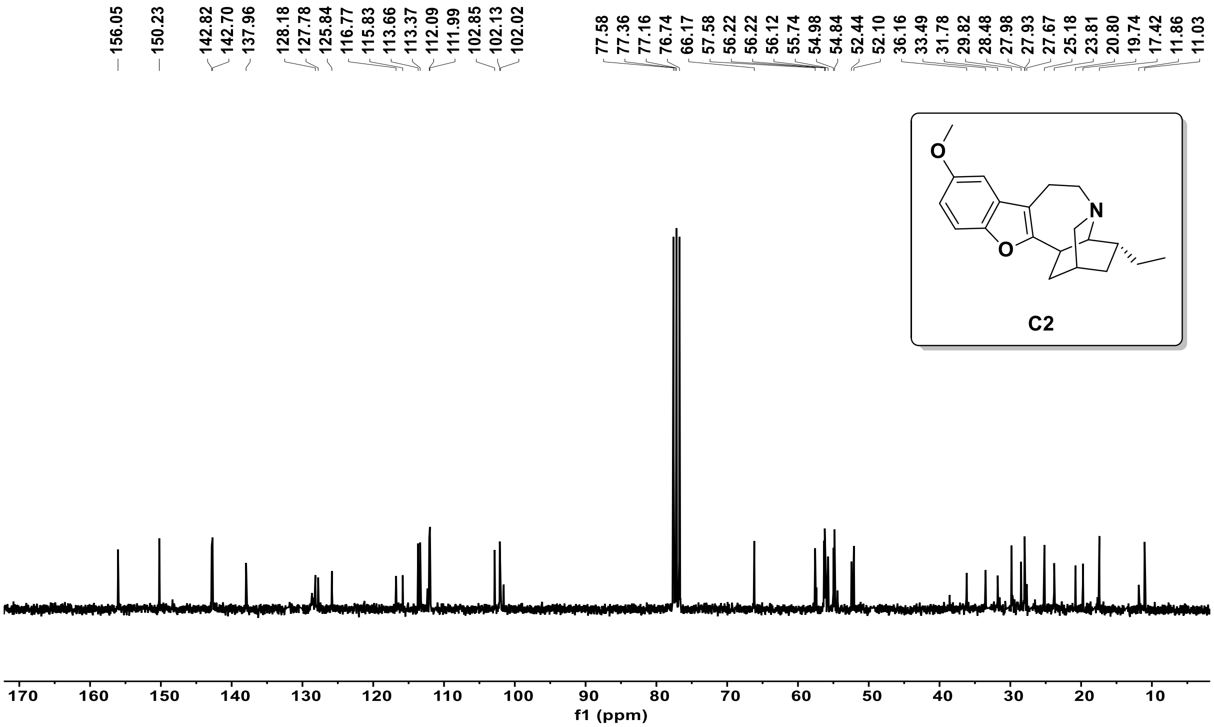


❖ DEPT-135 NMR (75MHz, $CDCl_3$) of compound **C2** :

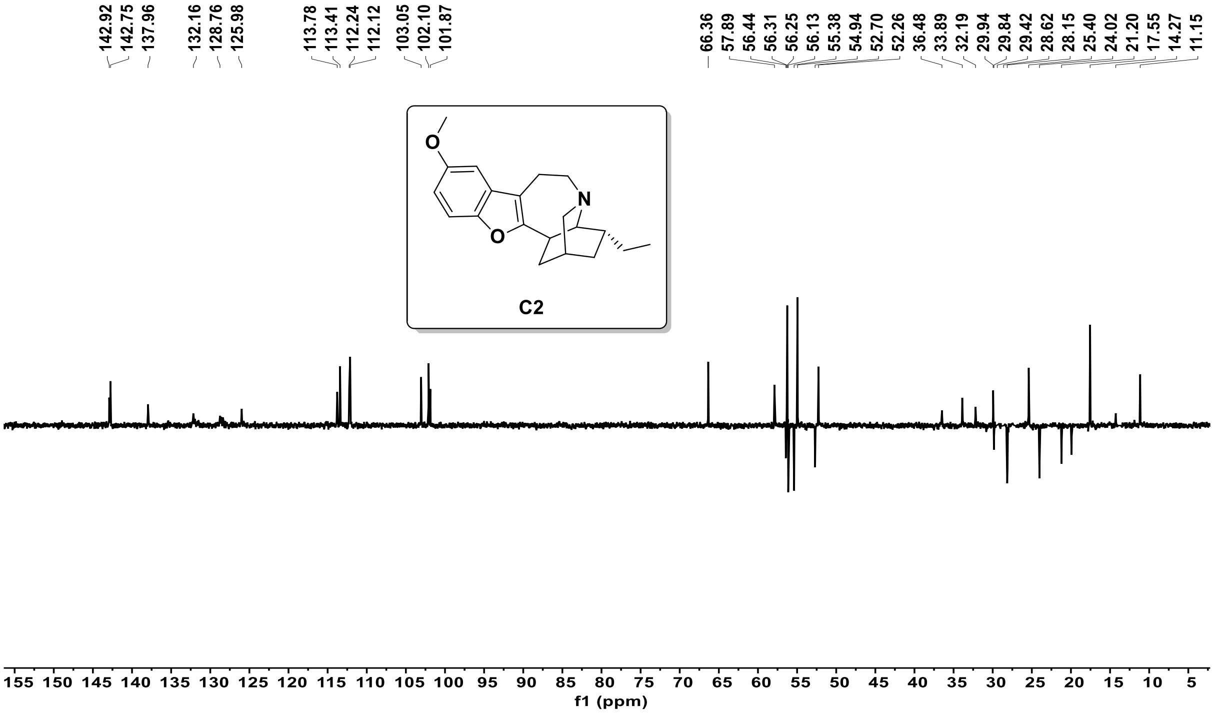

❖ HRMS (ESI, +ve ion mode) of compound **1a** :

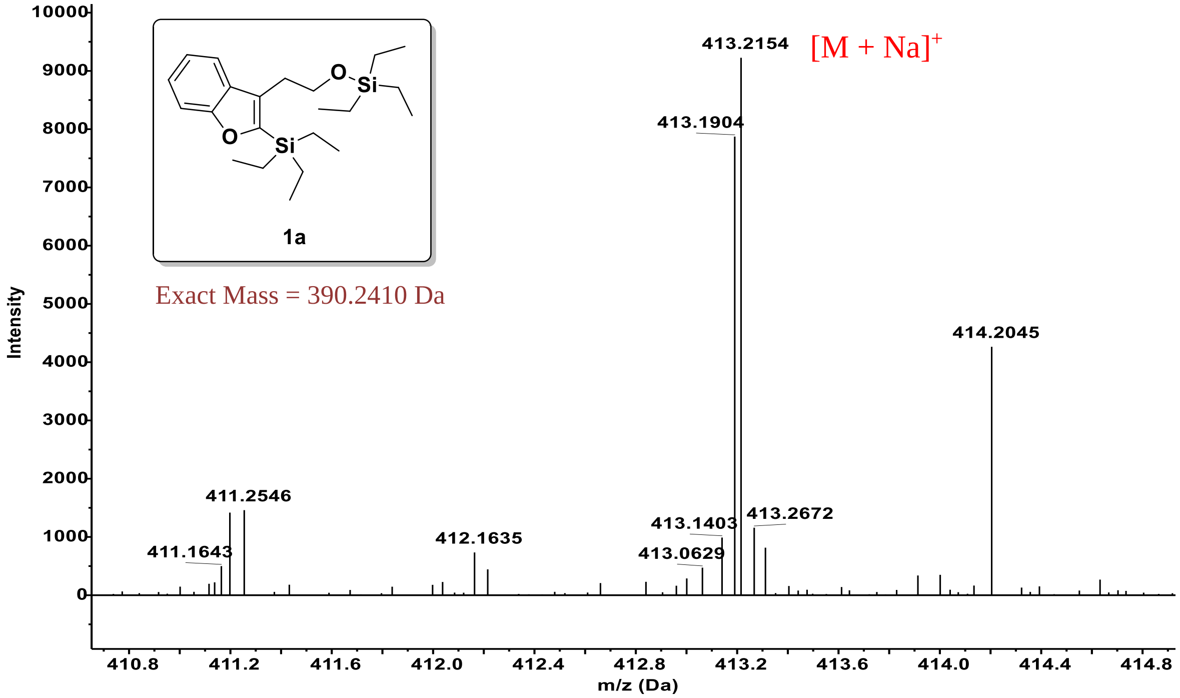


❖ HRMS (ESI, +ve ion mode) of compound **2** :

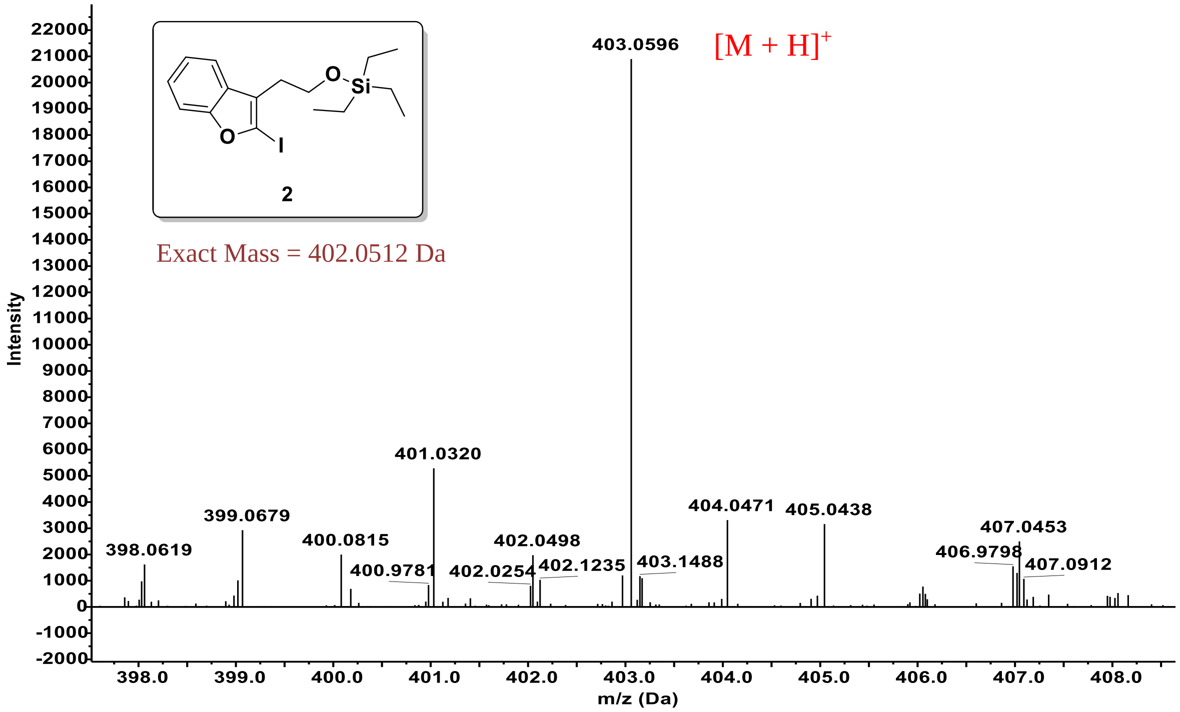

❖ HRMS (ESI, +ve ion mode) of compound **3** :

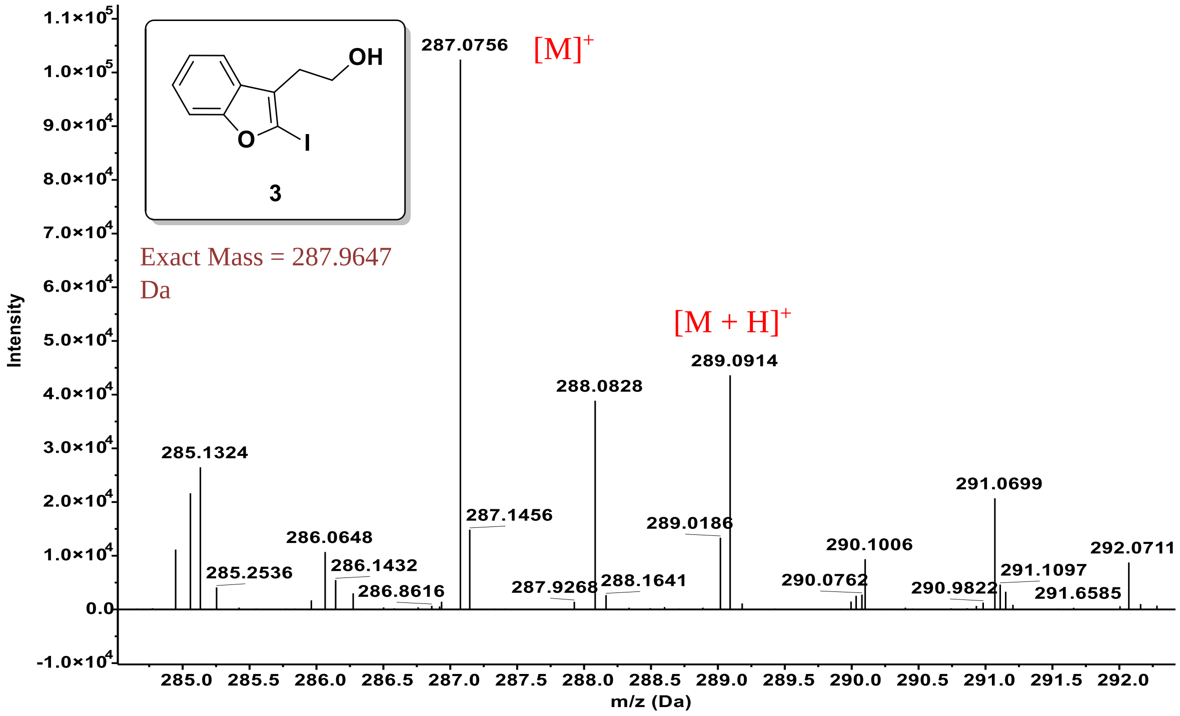


❖ HRMS (ESI, +ve ion mode) of compound **6a** & **6b** :

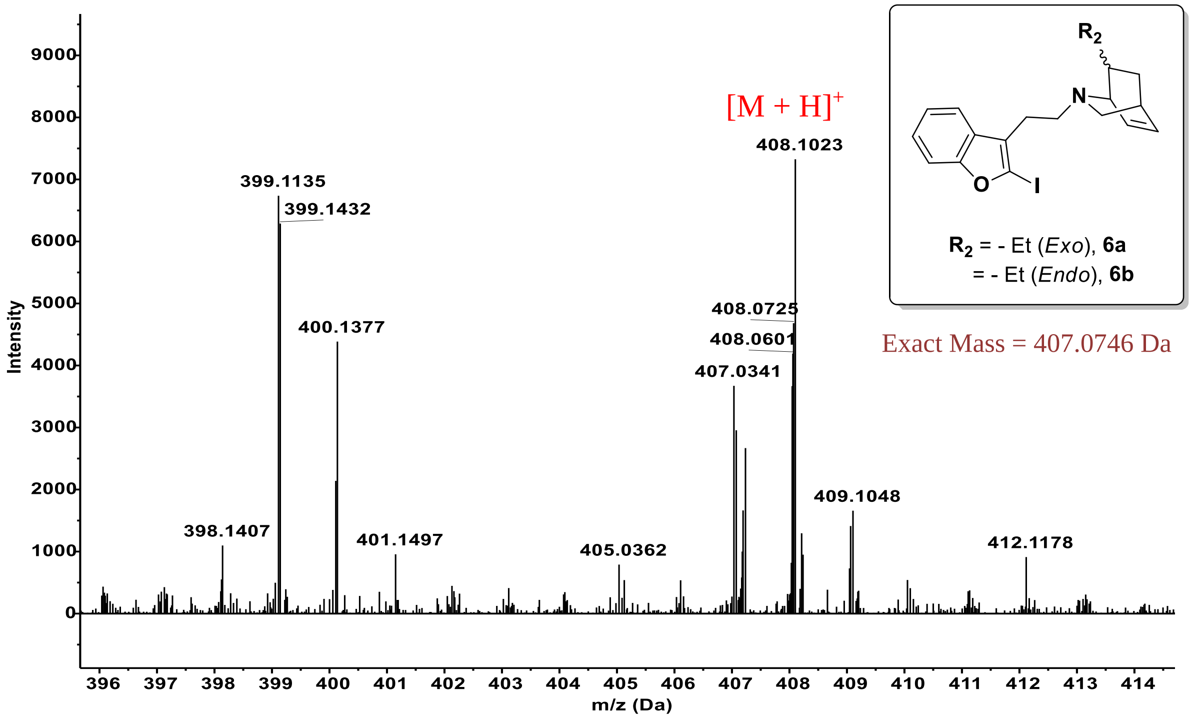

❖ HRMS (ESI, +ve ion mode) of compound **C3** :

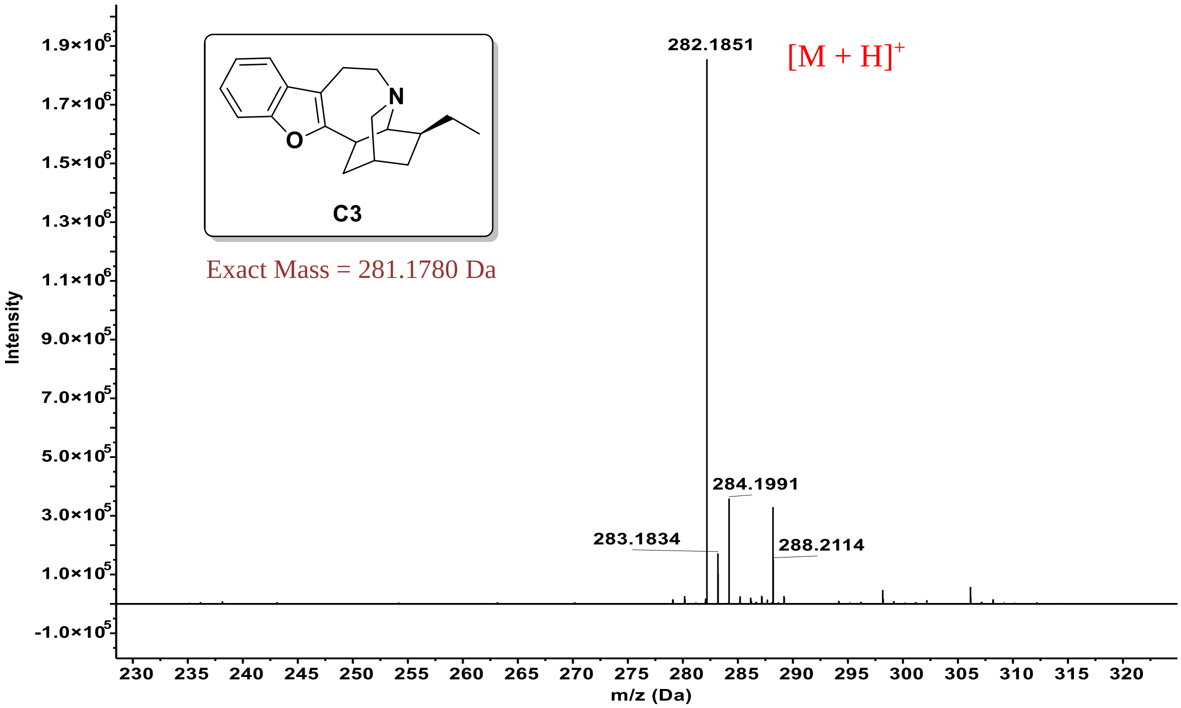


❖ HRMS (ESI, +ve ion mode) of compound **C4** :

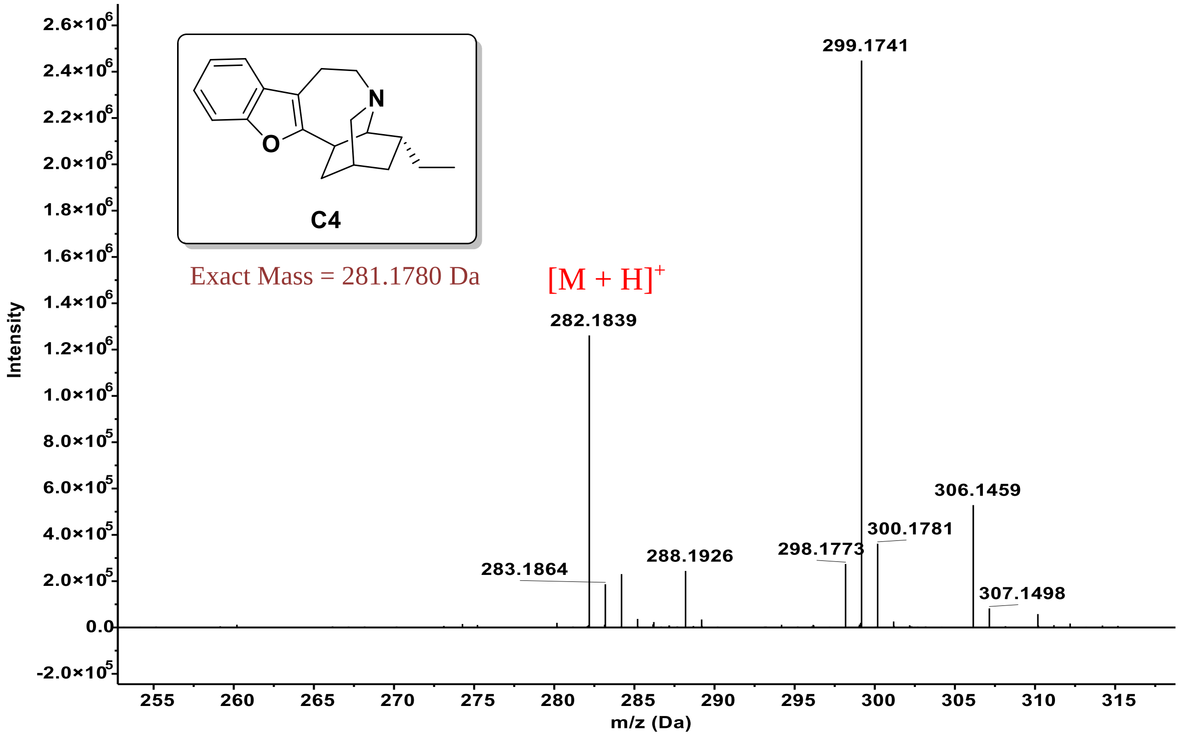

❖ HRMS (ESI, +ve ion mode) of compound **9** :

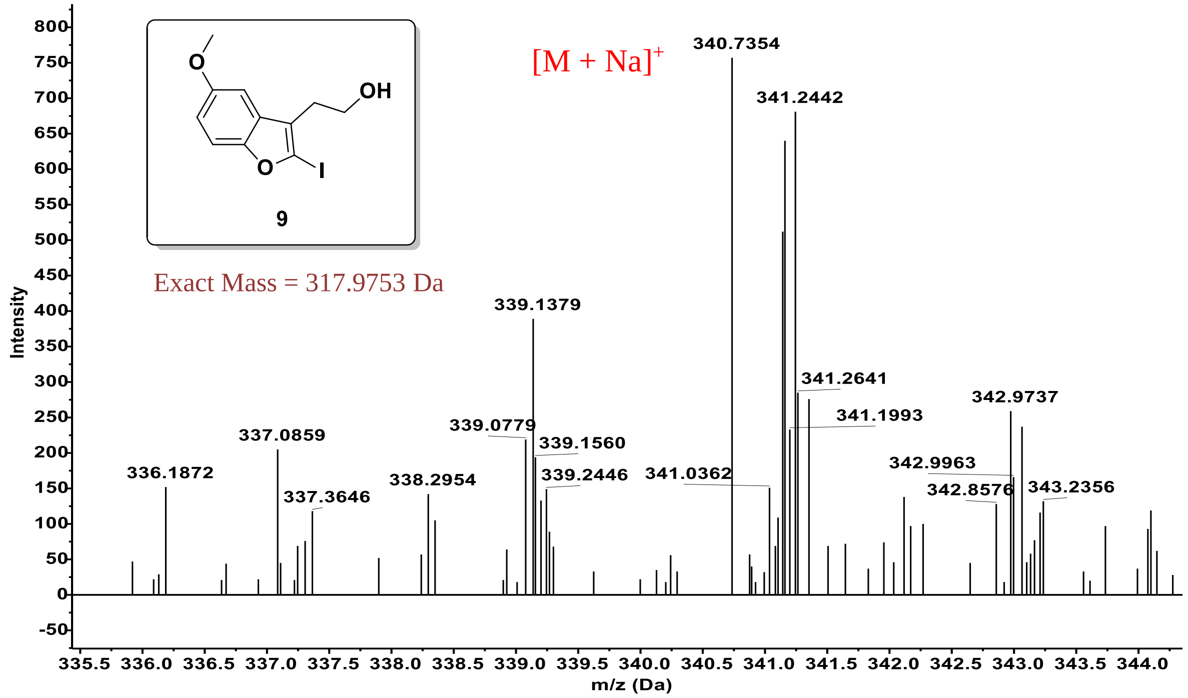


❖ HRMS (ESI, +ve ion mode) of compound **11a & 11b** :

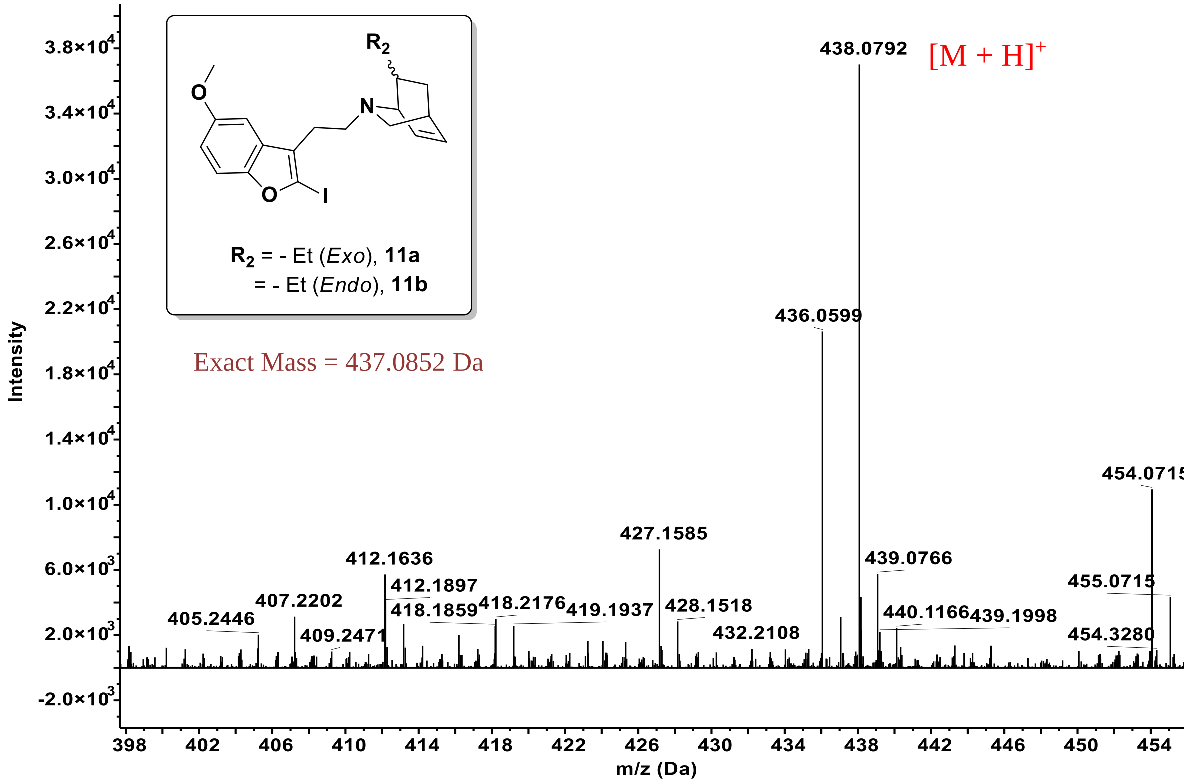

❖ HRMS (ESI, +ve ion mode) of compound **C1** :

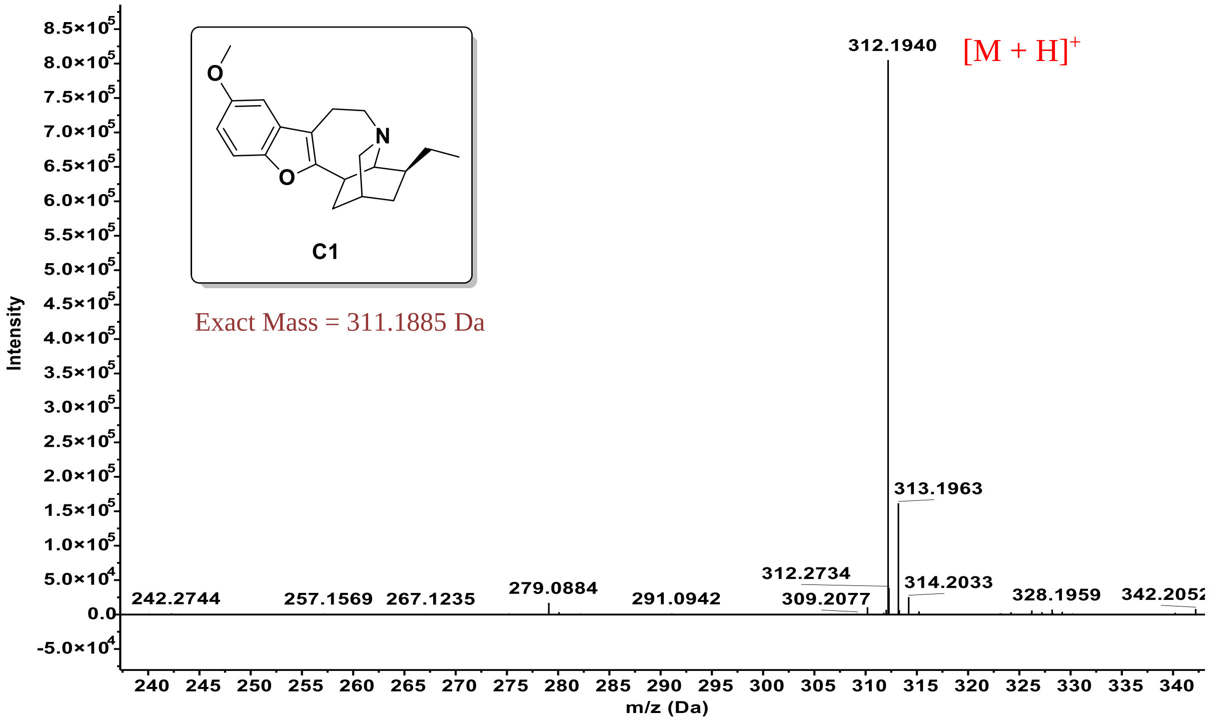


❖ HRMS (ESI, +ve ion mode) of compound **C2** :

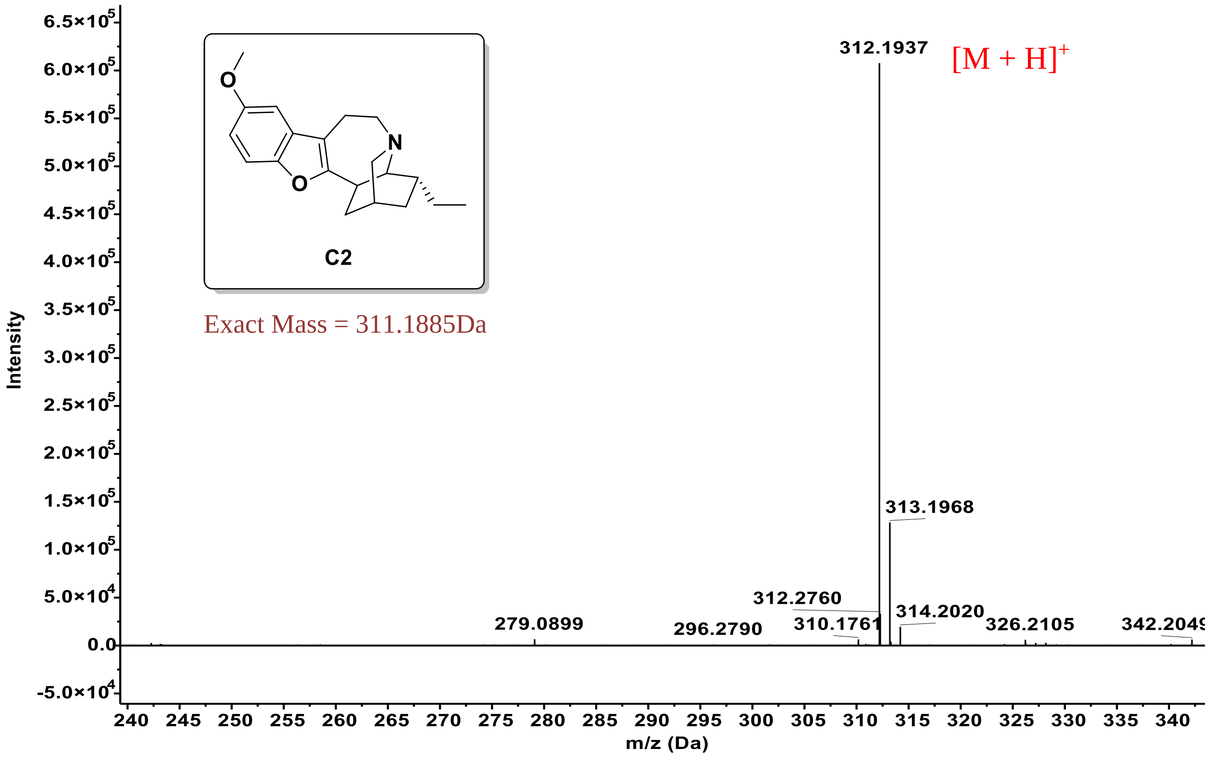

❖ HPLC of compound **C1-C4** :

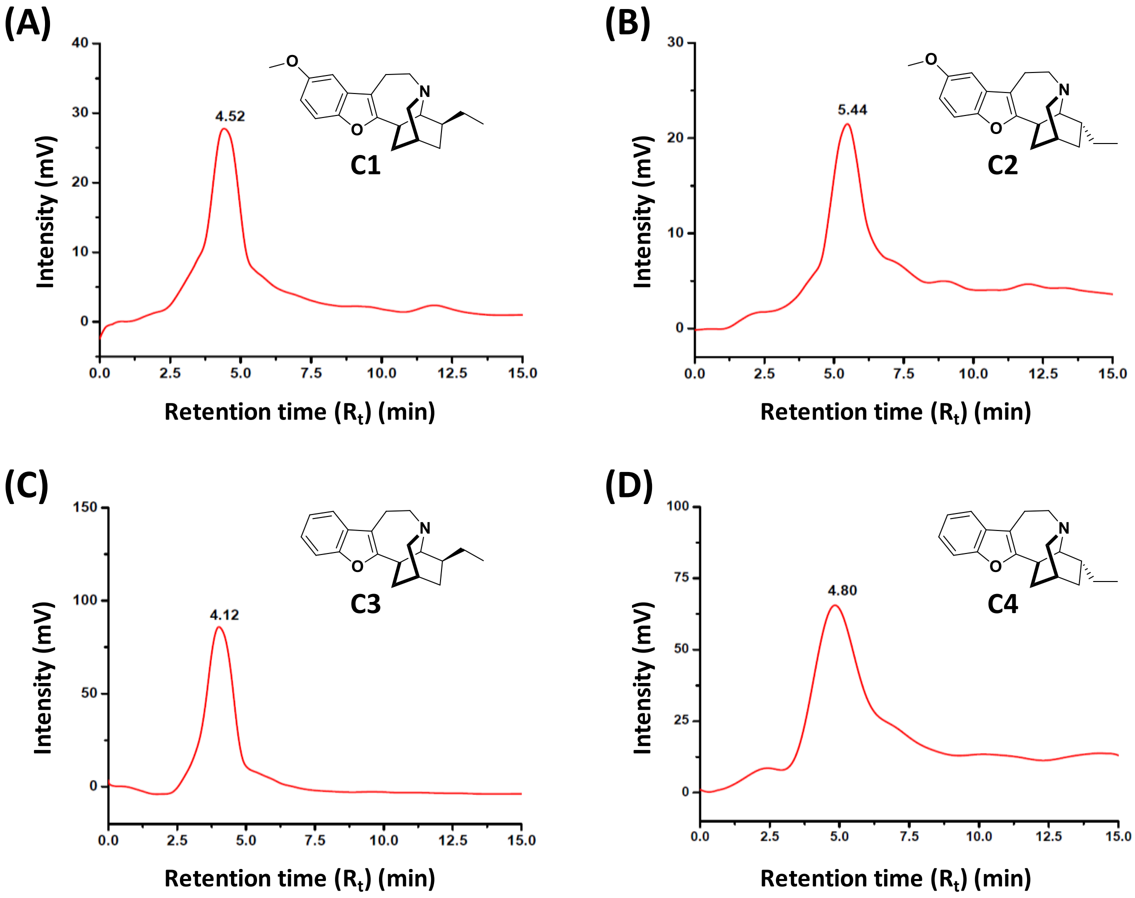


*<u>Gradient Used:</u>* 5% MeOH in $CH_3CN$ (Isocratic solution) for 15 min, *<u>Flow Rate:</u>* 1.0 ml/min.
*<u>Column used:</u>* Waters[R] SPHERISORB[R] ODS2 RP C18 Column (Analytical); 5 µm, 4.6 I.D × 250 mm.

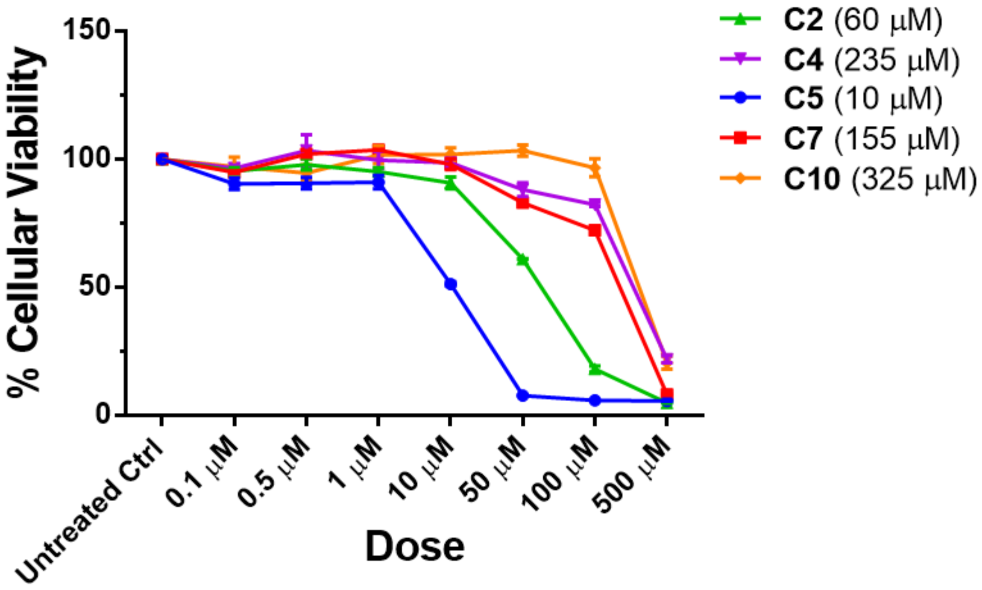


**Figure S1**: MTT-based cellular viability assay of iboga-compounds (**C2**, **C4**, **C5**, **C7** & **C10**) on C2C12 cells 96 h post-treatment. The $IC_{50}$ values were denoted within the parentheses of the legend. Results were replicated for at least three independent set of experiments. Values were represented as Mean ± SEM.

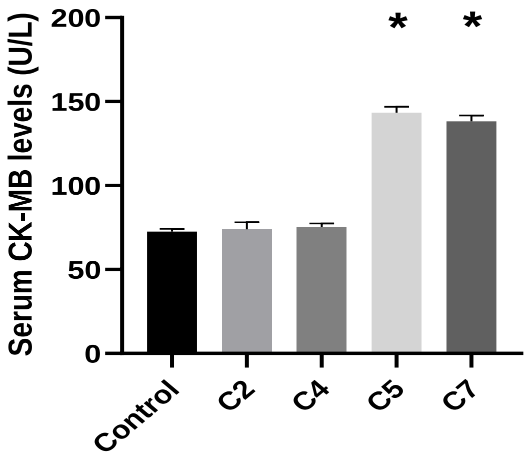


**Figure S2:** Examination of Serum CK-MB (creatine kinase myocardial band) levels to assess toxicity upon cardiomyocytes after iboga-treatments. Data were represented as Mean ± SEM (n = 3). *$P$ < 0.05, * indicated significant difference with the control group.

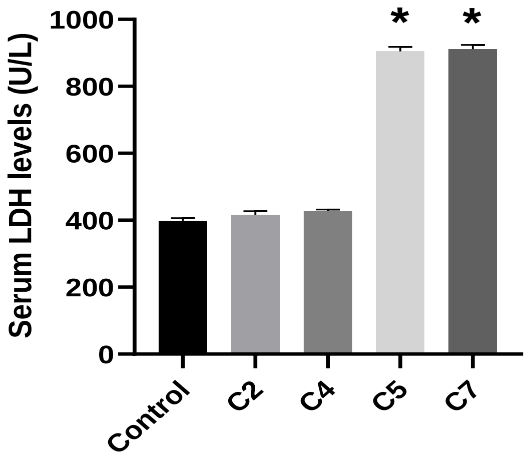


**Figure S3:** Investigation of Serum LDH (Lactate dehydrogenase) levels after iboga-treatments. Values were represented as Mean ± SEM (n = 3). *$P$ < 0.05, * indicated significant difference with the control group.

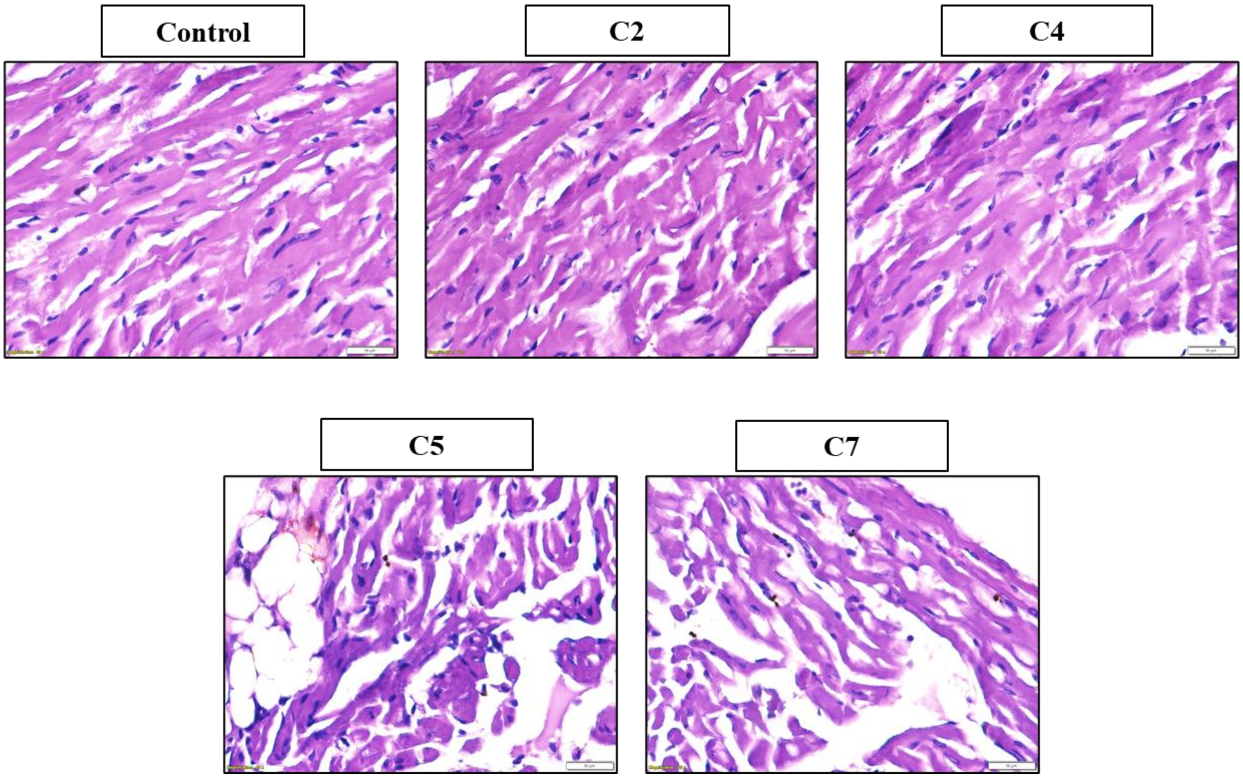


**Figure S4:** Analysis of histological samples of cardiac tissue by Hematoxylin and Eosin (H & E) staining after iboga-treatments. The degeneration of the ideal syncytial arrangement of myocytes was noted in the groups treated with natural ibogaine and ibogamine (**C5** and **C7**), as compared to the bioisosteric oxa-iboga analogs (**C2** and **C4**) and control-treated groups, clearly indicating a state of cardiomyopathy. All images were captured at an optical magnification of 400X using an Olympus bright field microscope. For visual interpretation of images, at least three images per group were considered.

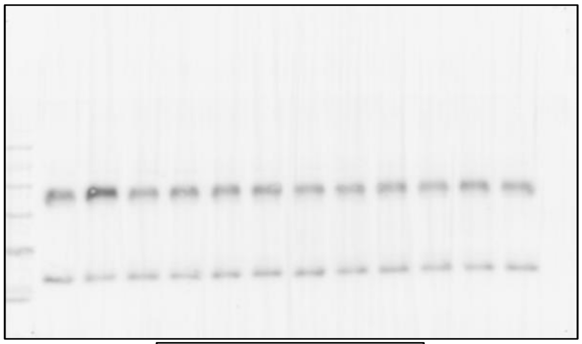

**p65 (nuclear)**

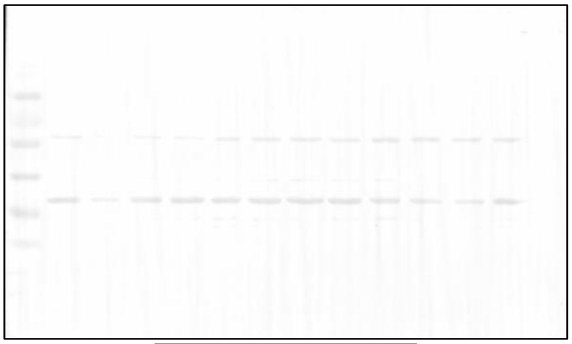

**BDNF**

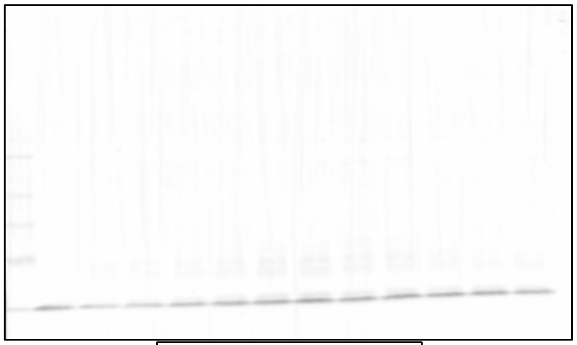

**H3 (nuclear)**

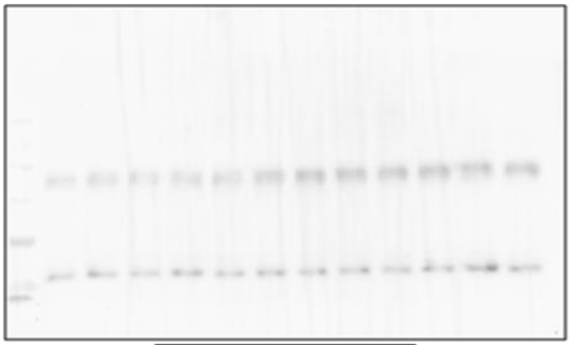

**β-actin**

Whole blot images of all the represented western blots. Samples from left to right direction are arranged as follows: (1) Molecular weight marker, (2) Control, (3) Formalin, (4) Formalin+**C1**, (5) Formalin+**C2**, (6) Formalin+**C3**, (7) Formalin+**C4**, (8) Formalin+**C5**, (9) Formalin+**C6**, (10) Formalin+**C7**, (11) Formalin+**C8**, (12) Formalin+**C9**, (13) Formalin+**C10**.

A

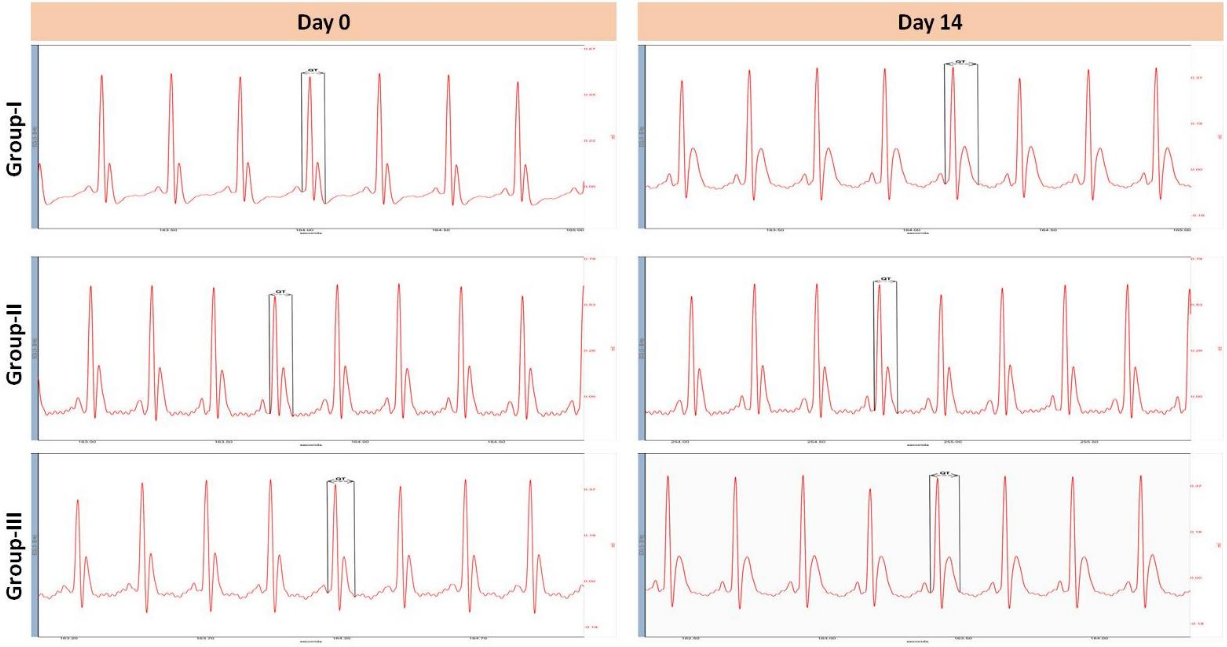


B

| Groups | | QT (sec) | RR (sec) | QTc (sec) |
|---|---|---|---|---|
| Group-I | Day 0 (Untreated Control) | 0.069 ± 0.001 | 0.246 ± 0.002 | 0.140 ± 0.005 |
| | Day 14 | 0.077 ± 0.001 | 0.253 ± 0.001 | 0.154 ± 0.004*** |
| Group-II | Day 0 (Untreated Control) | 0.068 ± 0.001 | 0.255 ± 0.003 | 0.135 ± 0.003 |
| | Day 14 | 0.067 ± 0.001 | 0.244 ± 0.004 | 0.137 ± 0.005 |
| Group-III | Day 0 (Untreated Control) | 0.072 ± 0.004 | 0.237 ± 0.002 | 0.148 ± 0.009 |
| | Day 14 | 0.075 ± 0.001 | 0.230 ± 0.003 | 0.157 ± 0.006* |

**Figure S5:** ECG-based cardiovascular toxicity assessment of different treated groups. (**A**)**.** ECG profile and (**B**)**.** Tabular representation of QTc and RR interval measured at both the start (Day 0) and the end (Day 14) of the experiment. Group-I, Group-II and Group-III represented **C5**, **C4** & **C2**-treated groups respectively. All compounds were administered at 15 mg/kg dose by oral gavage technique and continued for 14 consecutive days. Data represented as Mean ± SEM. *$P < 0.05$, ***$P < 0.001$, *** indicated significant difference between Day 0 & Day 14 treatment.
